# Genetic Variability, N-Glycosylation Sites, and Recombination Events in the ORF5 (GP5) Gene of Lineage 1A (NADC34) *Betaarterivirus americense* Strains from Lima, Peru Molecular Variability of the GP5 Glycoprotein in *Betaarterivirus americense*

**DOI:** 10.64898/2026.01.16.699519

**Authors:** Rony Cotaquispe Nalvarte, Felicita Karina Camargo Paredes, Hilda Victoria Coila De la Cruz, Edgard De la Cruz Vásquez, Julio Cesar Ecos Espino, Miriam Legua Barrios, Erick Llona García, Ayda Liliana Reyes Ruiz, Merici Ingrid Medina Guerrero, Elvia Mejía Vargas, Cesar Augusto Mendoza Yáñez

**Affiliations:** Research Professor, Universidad Científica del Sur, Lima, Peru; Professor, Universidad Privada Autónoma de Ica, Ica, Peru; Professor, Universidad Privada San Juan Bautista, Ica, Peru; Professor, Universidad Nacional San Luis Gonzaga, Ica, Peru; Research and Development Group in Molecular Biology and Clinical Diagnosis of Emerging Diseases, Universidad Privada San Juan Bautista, Ica, Peru; Animal Nutrition and Health Business Unit, Innovation and Development Area, Corporación Montana S.A., Lima, Peru

**Keywords:** genetic diversity, recombination, ORF5 (GP5), N-glycosylation, sublineage 1A (NADC34), RDP v4.1

## Abstract

Lima is a major swine production hub in Peru; therefore, these data provide an initial overview of GP5 variability in a high-risk epidemiological setting. This study aimed to characterize the genetic variability, N-glycosylation sites, and recombination events in the ORF5 (GP5) gene of lineage 1A (NADC34) *Betaarterivirus americense* strains circulating in Lima, Peru. Bioinformatics servers and software were employed, including Nextclade v3, NetNGlyc 1.0, RDP v4.1, DnaSP v6, and MEGA6. Nextclade v3 supported the classification of 24 strains within lineage 1, sublineage 1A (NADC34), which exhibited high divergence based on GP5 phylogenetic analysis. Significant amino acid substitutions were identified in the hypervariable regions HVR1 and HVR2, which are associated with the induction of neutralizing and non-neutralizing antibodies: 16/24 at N32 (S/G/R/E), 3/24 at S34 (N/T), 6/24 at S35 (N/I), 7/24 at L39 (F), 6/24 at Q40 (L/R), 3/24 at L41 (Y/V), 15/24 at K58 (E/V/R), and 8/24 at S59 (H/R/N), located within GP5 epitopes A, B, and C. Nine N-glycosylation patterns (A–I) were identified across the 24 strains, comprising nine putative sites at N30, N32, N33, N34, N35, N44, N50, N51, and N57. Patterns A, B, E, and G exhibited five to six glycosylation sites in 12/24 strains. A statistically robust recombination event was detected in strain 42_Montana2019 (lineage 1A), with a putative major parent MH719138_1 (lineage 1A, Peru-2016/5 variant; 95.3% sequence similarity) and an unknown minor parent, although strain 36_Montana2019 showed the closest phylogenetic affinity. Genetic diversity analysis revealed 530 polymorphic sites, and Tajima’s D test yielded a value of −0.78746, indicating high genetic variability within the Lima cohort. Overall, this study provides the first comprehensive molecular characterization of ORF5 (GP5) genetic variability in sublineage 1A (NADC34) *Betaarterivirus americense* strains circulating in swine farms in Lima. Broader and temporally structured sampling is warranted to assess nationwide evolutionary patterns.

## INTRODUCTION

Porcine reproductive and respiratory syndrome (PRRS) remains one of the most economically impactful diseases affecting the global swine industry due to its substantial health and production losses [11]. The etiological agent, currently classified by the International Committee on Taxonomy of Viruses (ICTV; MSL39.v3, 2023) as Betaarterivirus americense (PRRSV-2) and Betaarterivirus europensis (PRRSV-1), has been extensively characterized at the molecular and structural levels. These species were named according to the geographic origin of their prototype strains, VR-2332 and Lelystad, respectively [3,15,38].

The viral genome consists of a positive-sense single-stranded RNA molecule containing at least eleven open reading frames (ORFs): ORF1a, ORF1b, ORF2a, ORF2b, ORF3, ORF4, ORF5, ORF5a, ORF6, ORF7, and ORF2TF, in addition to a 3′ untranslated region (UTR) [3,38]. ORF1a and ORF1b occupy approximately three-quarters of the genome and encode the replicase polyproteins pp1a and pp1ab. The ORFs located in the 3′ terminal region encode eight structural proteins: four membrane-associated glycoproteins (GP2a, GP3, GP4, and GP5), three non-glycosylated membrane proteins (E, ORF5a, and M), and the nucleocapsid protein (N) [15,38].

Among these, ORF5 encodes the GP5 glycoprotein, the major envelope protein and a key determinant of viral infectivity. GP5 is a glycosylated transmembrane protein exposed on the virion surface, involved in receptor binding and recognized as one of the principal targets of the humoral immune response [13,39]. It harbors critical antigenic domains and multiple N-glycosylation sites associated with viral neutralization. N-glycosylation acts as an immune evasion mechanism by modulating epitope accessibility to neutralizing antibodies [13]. The high genetic variability of ORF5, driven by point mutations and recombination, promotes the emergence of novel B. americense variants and contributes to the expanding diversity of circulating genotypes [9,13]. Based on ORF5 sequence analysis, PRRSV-1 is classified into four subtypes, whereas PRRSV-2 comprises 11 monophyletic lineages (L1–L11) and multiple sublineages, including 1A–1F and 1H– 1J within L1, 5A–5B in L5, 8A–8E in L8, and 9A–9E in L9 [14,15,35,37].

In the present study, bioinformatic approaches were employed to assess genetic diversity and detect recombination events within ORF5. Recombination analysis was conducted using RDP v4.1, which integrates multiple algorithms (RDP, GENECONV, BootScan, MaxChi, Chimaera, SiScan, and 3Seq), and only events supported by multiple methods were retained to ensure statistical robustness [21,24]. DnaSP v6 was used to estimate genetic diversity parameters and polymorphic sites [27], while MEGA was applied for comparative phylogenetic analyses [9].

In Lima, Peru, the molecular detection of lineage 1, sublineage 1A (IA/2014/NADC34-like variant) in 2019 and again in 2025 has raised concerns regarding the extent of genetic diversification and the potential contribution of recombination to the evolution of circulating field strains. However, detailed molecular characterization of ORF5 variability in this epidemiological setting remains limited. Therefore, the aim of this study was to characterize the genetic diversity and evaluate recombination events shaping the molecular architecture of GP5 in field strains from Lima, Peru. These findings provide novel insights into the evolutionary dynamics and molecular epidemiology of PRRSV-2 in a high-density swine production region and offer relevant evidence for understanding current challenges in vaccine efficacy and viral control strategies.

## MATERIALS AND METHODS

### Biological Material

A total of 24 nucleotide sequences obtained from field strains collected from commercial swine farms in Lima, Peru, were analyzed in this study [6]. These sequences corresponded to the ORF5 gene of *Betaarterivirus americense*.

### RNA Extraction, Quantification, and Conventional RT-PCR of ORF5

Viral RNA extraction and complementary DNA (cDNA) synthesis of *B. americense* were performed as previously described by Cotaquispe [6]. Conventional RT-PCR targeting the ORF5 gene was conducted under the same conditions reported in that study.

### Electrophoresis, Purification, and Sequencing

Amplified products were visualized using a horizontal electrophoresis system (multiSUB™ Choice, Cleaver Scientific) with 1% molecular-grade agarose gels (SCL AG500, Cleaver Scientific). Fluorescent Dye-DNA/Safe-Green™ (ABM®) was used for nucleic acid staining, and a 100 bp Opti-DNA ladder (50 bp–1.5 kb, ABM®) served as the molecular size marker. Gels were documented using a UVP UV Solo Touch imaging system (Analytik Jena). Gel fragments containing amplicons of the expected size were excised and purified using the innuPREP Nucleic Acid Purification Kit (Analytik Jena). Purified PCR products were sequenced bidirectionally by Macrogen (Seoul, South Korea) using the BigDye® Terminator v3.1 Cycle Sequencing Kit (Life Technologies) and analyzed on an ABI 3137 XL Genetic Analyzer (Life Technologies), as previously described [5].

### Phylogenetic Analysis of GP5

Raw chromatograms were inspected and edited using Chromas Lite® v2.6.6. Lineage assignments were determined exclusively based on GP5 nucleotide phylogeny. Sequences were analyzed using Nextclade v3 (https://clades.nextstrain.org/) [2] to refine local phylogenetic relationships and improve lineage resolution. The dataset was combined with 46 reference sequences retrieved from GenBank, including nine Peruvian isolates (MH791386.1; MH791385.1; MH791383.1; MH791388.1; MH791379.1; MH791378.1; MH791376.1; MH791391.1; MH791390.1).

### Antigenic Structure and N-Glycosylation Analysis of GP5

Amino acid sequences were analyzed using MEGA v6.0 [30] to characterize structural and immunologically relevant domains, including the signal peptide, decoy epitopes I and II, hypervariable regions I and II (HVR1 and HVR2), the primary neutralizing epitope (PNE), transmembrane regions 1–3, T-cell epitopes 1 and 2, and B-cell epitopes. This analysis enabled the identification of amino acid substitutions within key functional domains that may influence viral infectivity, pathogenicity, and persistence [24].

The full length of the GP5 protein was determined for each strain, and potential N-glycosylation sites (N-X-S/T motifs) were predicted using the NetNGlyc 1.0 server (NetNGlyc 1.0 - DTU Health Tech - Bioinformatic Services) [10,24]. Signal peptides were predicted using SignalP v4.0 [25].

### Genetic Diversity Analysis: Tajima’s D and DnaSP v6

Genetic diversity was evaluated under the framework of the neutral theory of molecular evolution. Tajima’s D statistic [31] was calculated to compare two estimates of nucleotide diversity (θ) within the sequence dataset. The statistic has an expected mean of zero under neutrality and reflects deviations that may suggest selection or demographic effects.

D values were interpreted according to the appropriate confidence intervals based on sample size [32]. Negative values indicate an excess of low-frequency polymorphisms, consistent with purifying selection or population expansion, whereas positive values may suggest balancing selection or population bottlenecks. Tajima’s D and additional diversity parameters were calculated using DnaSP v6 [27].

### Recombination Detection Analysis of ORF5 Using RDP v4.1

The ORF5 alignment dataset was initially screened for recombination signals. Recombination detection was performed using seven independent methods implemented in RDP v4.1 (http://web.cbio.uct.ac.za/~darren/rdp.html): RDP [18], GENECONV [23], BootScan [19], MaxChi [29], Chimaera [26], SiScan [8], and 3Seq [17], with default parameters [21].

Recombination events were considered reliable only when supported by at least four independent methods with associated p-values < 0.05 [15,33], thereby ensuring statistical robustness and minimizing false-positive signals.

## RESULTS

### ORF5 Amplification and Sequencing

Serum samples were analyzed using an in-house RT-PCR assay targeting the ORF5 gene, which encodes the GP5 glycoprotein of *Betaarterivirus americense*. Primers were validated and standardized using vaccine strains of both PRRSV species as positive controls, along with a commercial IDEXX positive control mixture. Amplification yielded a 723 bp product (Figure 1). Sequencing of all partial amplicons confirmed the presence of the complete ORF5 coding region, comprising 603 nucleotides and encoding a functional protein of 201 amino acids.

**Figure 1.**
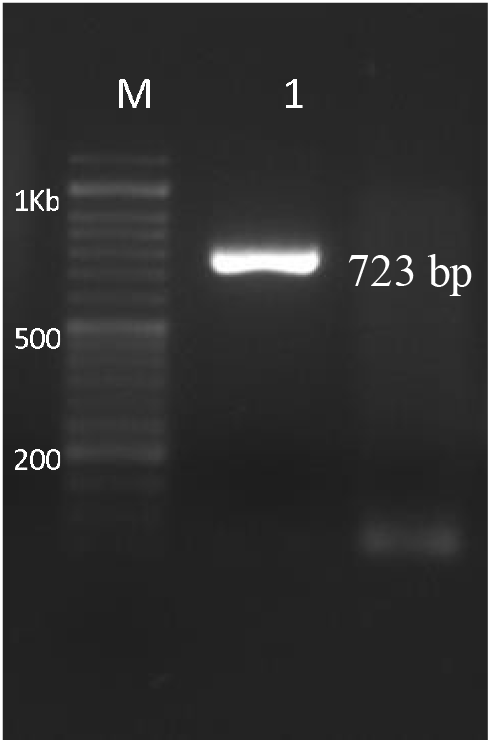
In-house conventional RT-PCR of the ORF5 gene designed in the laboratory. Lane 1 (ORF5-SKA): cDNA from a PRRS field-positive sample; Lane 2 (avian bronchitis): negative control cDNA from IBV; Lane M (ladder): 100 bp Plus Opti-DNA Marker ABM® (50 bp–1.5 kb).

### Phylogenetic Analysis of GP5

Phylogenetic relationships among the 24 study strains were inferred from GP5 nucleotide sequence alignments combined with representative GenBank reference strains from lineages 1, 5, and 8 using Nextclade v3. The resulting topology is shown in Figure 2. Nine previously reported Peruvian isolates belonging to sublineage 1A (1-7-4/NADC34-like variant) were included for comparative analysis. The phylogeny resolved multiple well-supported branches corresponding to PRRSV-2 lineages, including: Lineage 1: sublineage 1E (USA strains 99-3584, 99-3298, 98-3403), 1A (NADC34), 1B (NADC31), 1C (HENAN), and 1F (MN184A); Lineage 5: sublineage 5A (VR-2332-like); Lineage 8: sublineages 8A (Ingelvac) and 8E (JXA1). All study strains clustered within lineage 1, sublineage 1A (NADC34-like), forming a well-defined clade together with Peruvian isolates reported between 2015 and 2017, indicating strong phylogenetic affinity and local evolutionary continuity (Figure 2).

**Figure 2.**
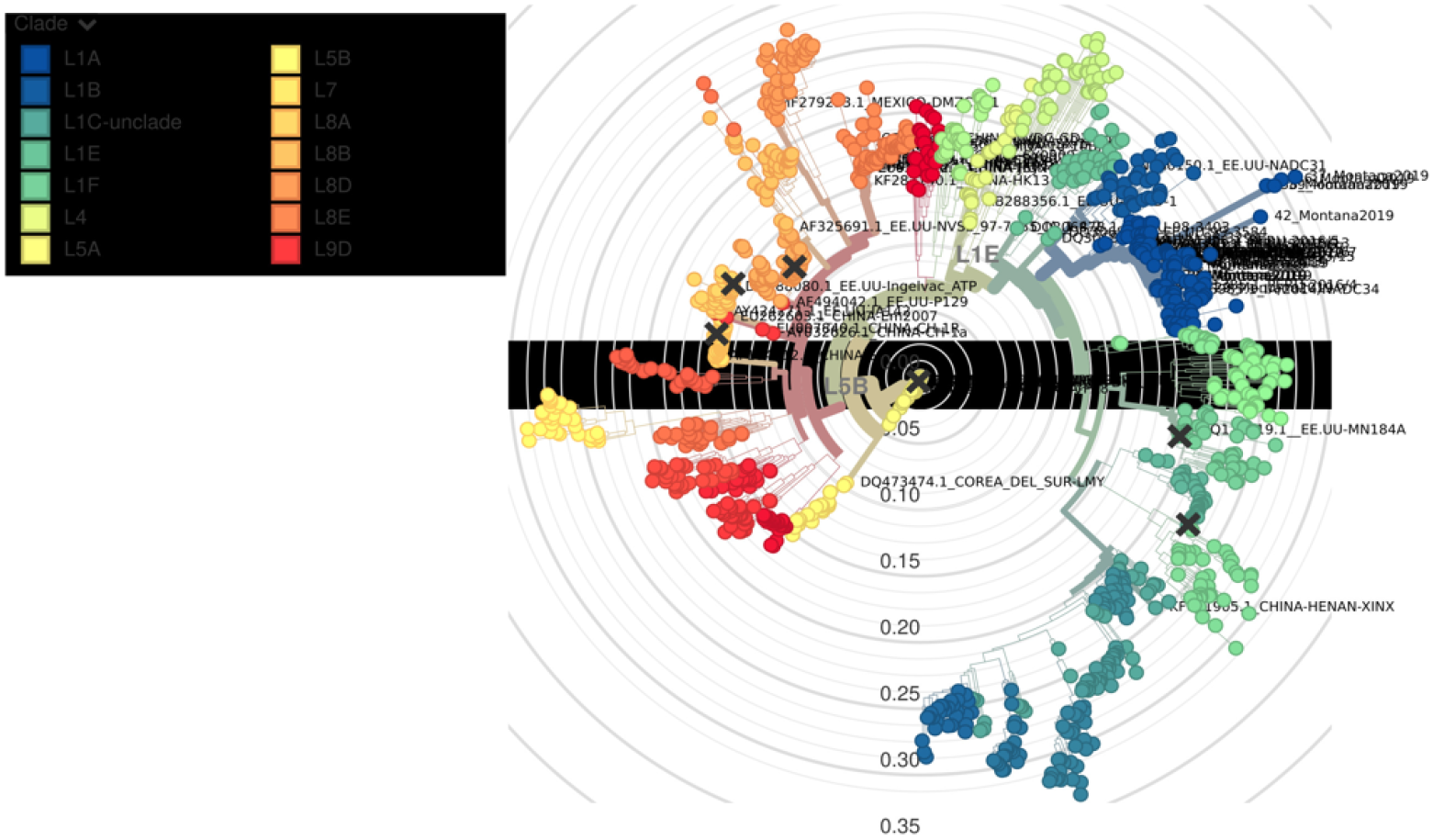
Phylogenetic tree of the species *Betaarterivirus americense* based on the glycoprotein 5 (GP5) sequence, generated using the Nextclade v3 server (https://clades.nextstrain.org/) (Aksamentov et al., 2021). Branches correspond to existing lineages, color-coded as follows: Lineage 1 — sublineage 1E (turquoise) USA (99-3584), sublineage 1A NADC34 (dark blue), sublineage 1B (NADC31) (light blue), sublineage 1C (HENAN) (sky blue), sublineage 1F (MN184A) (forest green); Lineage 5 — sublineage 5A (VR2332) (yellow); and Lineage 8 — sublineage 8A (Ingelvac) (light orange) and sublineage 8E (JXA1) (dark orange). The 24 strains analyzed in this study belong to sublineage 1A (dark blue), with divergent variants observed within the group.

### Amino Acid Diversity of GP5

The translated GP5 sequences (201 amino acids) exhibited the expected structural organization: signal peptide (aa 1–31), decoy epitope I (aa 27–36), decoy epitope II (aa 177–197), primary neutralizing epitope (PNE, epitope B; aa 37–44), neutralizing epitope C (aa 51–60), transmembrane regions 1 (aa 64–80), 2 (aa 94–102), and 3 (aa 108–125), T-cell epitopes (aa 115–129 and 148–161), B-cell epitope (aa 177–200), and a stop codon at position 201 (Figure 3). Multiple amino acid substitutions were identified within antigenically relevant domains: **Signal peptide (aa 1–26):** several neutral substitutions were observed. **Decoy epitope I / HVR1:** substitutions included 12/24 at A27 (V/S), 16/24 at N32 (S/G/R/E), 3/24 at S34 (N/T), and 6/24 at S35 (N/I). **Decoy epitope II:** 2/24 substitutions at T190P and 1/24 at K177N, potentially affecting cross-neutralization dynamics. Within the primary neutralizing epitope (PNE, epitope B), which contains key antigenic determinants at positions 39, 40, and 43, substitutions were observed in 7/24 strains at L39F, 6/24 at Q40L/R, 4/24 at N43I, and 4/24 at T46A. In neutralizing epitope C and the adjacent HVR2 region, substitutions were detected in 15/24 strains at K58E/V/R and 8/24 at S59H/R/N. These changes are consistent with previously described mechanisms facilitating escape from neutralizing antibodies.

**Figure 3.**
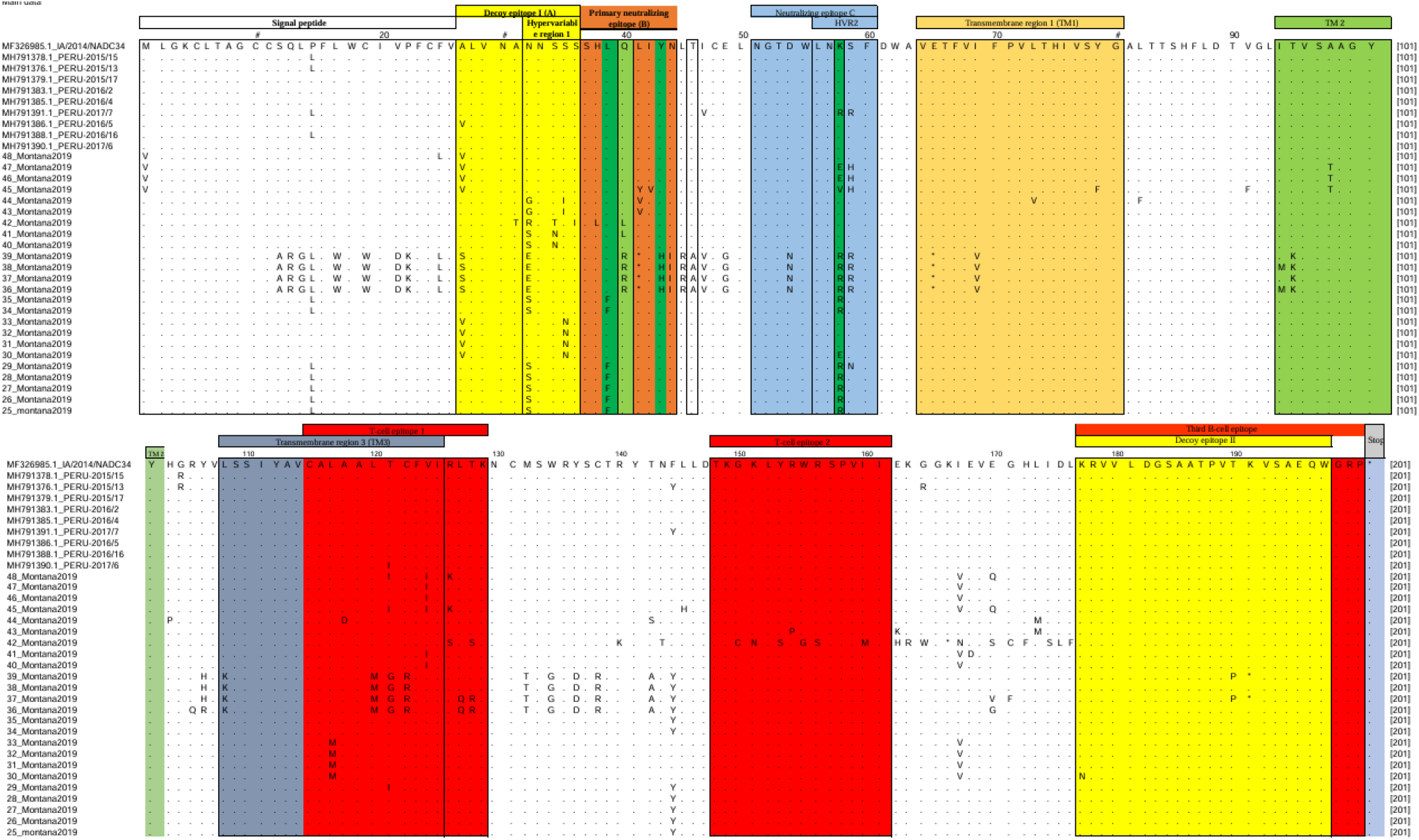
Alignment of 24 amino acid sequences of the GP5 antigenic regions in *Betaarterivirus americense* sublineage 1A (NADC34). The boxed and color-coded regions depict the structure of glycoprotein 5 with antibody-neutralizing epitopes. The 24 field strains, the reference lineage MF326985.1_IA/2014/NADC34, and nine representative Peruvian variants of sublineage 1A (NADC34) were used as reference patterns in the alignment. Dots indicate positions of conserved amino acids, while letters denote amino acid substitutions.

### N-Glycosylation Pattern Analysis of GP5

Prediction analysis identified nine distinct N-glycosylation patterns (A–I). In total, nine potential N-glycosylation sites (N-X-S/T motifs) were detected: N30, N32, N33, N34, N35, N44, N50, N51, and N57. The major neutralization-associated sites N33, N44, and N51 were present in most strains. Notably, 20/24 isolates harbored N44 and N51. All isolates contained at least three predicted N-glycosylation sites. A previously unreported glycosylation site at N57 was identified. Strains 40 and 41 exhibited the highest number of predicted glycosylation sites (Table 1). Signal peptide cleavage prediction using SignalP 4.0 indicated enzymatic cleavage between residues 31 and 32, consistent with canonical GP5 processing (Table 1).

**Table 1.** Locations and patterns of potential N-glycosylation sites in GP5 among 24 sequences of *Betaarterivirus americense* from Lima, Peru.

| Strains | GP5 (aa) | Signal peptide | Glycosylation pattern | N-Glycosylation sites |  |  |  |  |  |  |  |  | % of total | Sublineages |
| --- | --- | --- | --- | --- | --- | --- | --- | --- | --- | --- | --- | --- | --- | --- |
|  |  |  |  | N30 | N32 | N33 | N34 | N35 | N44 | N50 | N51 | N57 |  |  |
| 25-48 | 201 | 1-31 | A | + |  | + |  |  | + |  | + | + | 25% (6/24) | L1A |
|  |  |  | B |  | + |  |  | + | + |  | + | + | 16.67% (4/24) | L1A |
|  |  |  | C |  |  | + |  |  |  | + |  |  | 16.67% (4/24) | L1A |
|  |  |  | D |  | + | + |  |  | + |  | + |  | 12.5% (3/24) | L1A |
|  |  |  | E | + |  | + | + |  | + |  | + | + | 8.33% (2/24) | L1A |
|  |  |  | F |  |  |  |  |  | + |  | + | + | 8.33% (2/24) | L1A |
|  |  |  | G |  | + | + |  |  | + |  | + | + | 4.16% (1/24) | L1A |
|  |  |  | H |  |  | + |  |  | + |  | + | + | 4.16% (1/24) | L1A |
|  |  |  | I | + |  | + |  |  | + |  | + |  | 4.16% (1/24) | L1A |

### Genetic Diversity and Tajima’s D Analysis

Genetic diversity analysis using DnaSP v6 revealed substantial nucleotide heterogeneity among GP5 sequences, with 530 polymorphic sites and 61 conserved sites (Table 2).

**Table 2.** Results of the genetic diversity of 24 isolates of *Betaarterivirus americense* analyzed using the DnaSP and MEGA 6 programs from blood samples of pigs from the Lima region, Peru.

| Strains | Sequence | ORF 5 (GP5) | C.S. | Variable sites | P. I. S | U. V. S | D Tajima |
| --- | --- | --- | --- | --- | --- | --- | --- |
| 24 | Nt (aa) | 588 (196) | 61 | 530 | 48 | 82 | -0,78746 |

**Nt:** nucleotide; **aa:** amino acid; **C.S.:** conserved sites; **P.I.S.:** parsimony-informative sites; **U.V.S:** unique variable sites.

**Table 3.**
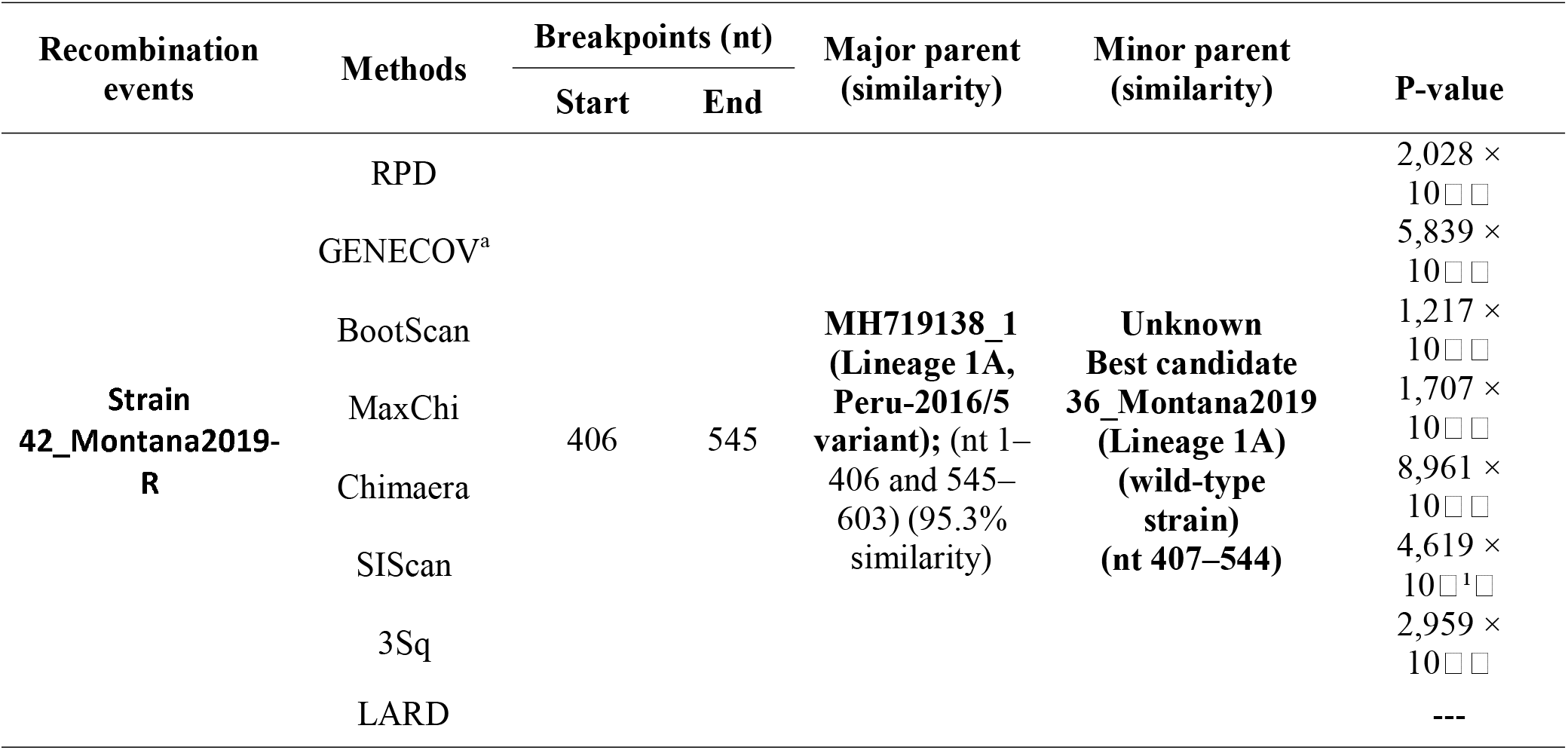
Genetic recombination events of the 42_Montana2019-R strain detected using RDP software version 4.1.

Tajima’s D test yielded a value of −0.78746, with no statistically significant deviation from neutrality (P > 0.10). The negative value suggests an excess of low-frequency polymorphisms, potentially consistent with purifying selection or population expansion, although the result did not reach statistical significance.

### Recombination Analysis of ORF5 Using RDP v4.1

Recombination analysis performed using RDP v4.1 identified a statistically significant recombination event in strain 42_Montana2019 (lineage 1A). The event was consistently detected by all seven implemented algorithms (RDP, GENECONV, BootScan, MaxChi, Chimaera, SiScan, and 3Seq), fulfilling conservative criteria for recombination inference. Topological incongruence between phylogenies reconstructed from non-recombinant regions (nt 1–406 and 545–603) and the recombinant region (nt 407–544) supported this event (Figure 4): The major parental sequence corresponded to GenBank strain MH719138_1 (lineage 1A, Peru-2016/5 variant) in the non-recombinant regions. The minor parental sequence was not directly identified in the dataset; however, strain 36_Montana2019 (lineage 1A) exhibited the closest phylogenetic affinity in the recombinant region. Statistical support for the recombination signal was highly significant across methods: RDP (2.028 × 10□□), GENECONV (5.839 × 10□□), BootScan (1.217 × 10□□), MaxChi (1.707 × 10□□), Chimaera (8.961 × 10□□), SiScan (4.619 × 10□^1^□), and 3Seq (2.959 × 10□□) (Table 4). Detection by seven independent algorithms markedly reduces the likelihood of a false-positive inference and supports the robustness of the identified recombination event.

**Figure 4.**
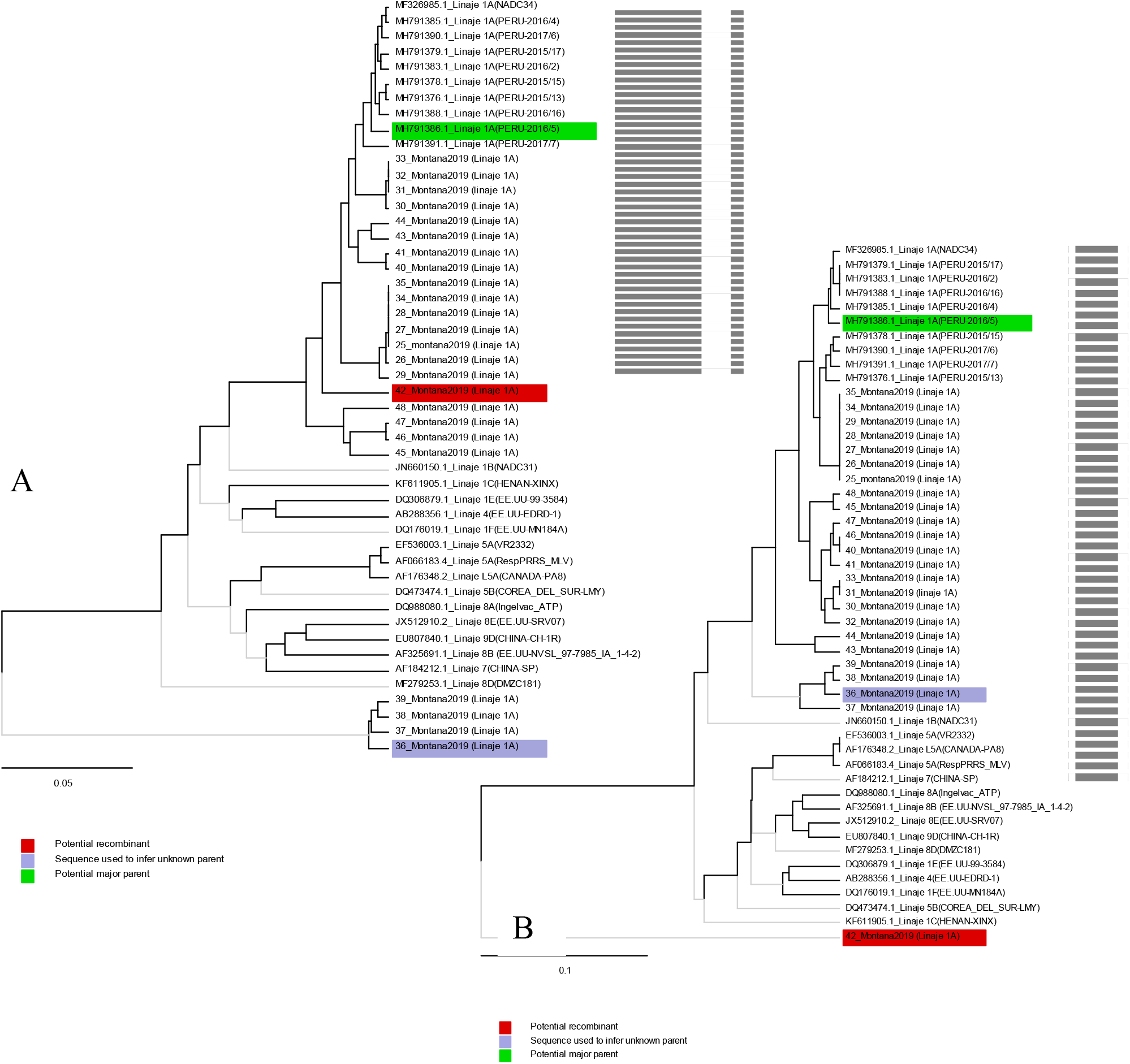
Genomic recombination analysis of the 42_Montana2019-R strain of porcine reproductive and respiratory syndrome virus (PRRSV), Lima, Peru, 2026. A) UPGMA tree of the region derived from the major parent MH719138_1 (Lineage 1A, Peru-2016/5 variant) (nt 1–406 and 545–603). B) UPGMA tree of the region derived from the minor parent 36_Montana2019 (Lineage 1A) (nt 407–544). Phylogenies of the parental strains were identified using RDP software version 4.101 (http://web.cbio.uct.ac.za/~darren/rdp.html). Red indicates the recombinant strain (42_Montana2019-R); green indicates the major parental strain from GenBank (MH719138_1) (Lineage 1A, Peru-2016/5 variant); translucent blue indicates the best candidate for the minor parental strain 36_Montana2019 (Lineage 1A) (wild-type strain). Scale bars indicate nucleotide substitutions per site.

## DISCUSSION

All 24 sequences analyzed in this study belonged to the species *Betaarterivirus americense*. Phylogenetic inference was performed using FASTA-formatted sequences and aligned with globally significant epidemic strains available in the Nextclade v3 database [2], facilitating lineage assignment for lineages 1, 5, and 8. The GP5 phylogeny generated in this study confirmed that all study strains clustered within lineage 1, sublineage 1A (NADC34-like variant), with notable variability observed among strains 36, 37, 38, 39, and 42.

Structural analyses of GP5 indicate that antigenic domains play crucial roles in immune recognition. Both synonymous and nonsynonymous substitutions occur in key immunological regions, including decoy epitopes I (A) and II, as well as primary and secondary neutralizing epitopes (B and C), influencing non-neutralizing and neutralizing antibody responses and contributing to immune evasion [9,16]. Compared with the reference strain MF326985.1_IA/2014/NADC34, varying degrees of substitution were observed across the signal peptide and antigenic domains. Within decoy epitope I and the HVR1 region, substitutions were particularly frequent (N32 S/G/R/E in 16/24 strains; S34 N/T in 3/24; S35 N/I in 6/24), consistent with rapid induction of non-neutralizing antibody responses [12,34]. These decoy epitopes (aa 27– 36 and aa 177–196) likely delay the production of effective neutralizing antibodies, facilitating viral persistence [39].

Within the primary neutralizing epitope B, substitutions at L39F (7/24), Q40L/R (6/24), and L41Y/V (3/24), along with four nonsense mutations due to deletions, indicate considerable variability. Prior studies identified aa38, aa42, aa43, and aa44 as critical neutralizing determinants, and aa39–41 as antibody-binding sites, with aa39 often used to classify viral variants [7,39]. Mutations in these regions may reduce PRRSV-2 sensitivity to neutralizing antibodies in vivo, potentially affecting viral clearance and contributing to vaccine failure and increased morbidity [4,22]. While neutralization assays were not performed here, functional studies targeting these domains are warranted to evaluate their effect on pathogenesis and vaccine escape.

For neutralizing epitope C and HVR2, substitutions at K58E/V/R (15/24) and S59H/R/N (8/24) may also facilitate escape from neutralizing antibodies [22]. Counts of substitutions in epitopes A, B, and C are descriptive, highlighting positional variability in GP5; formal site-specific selection analyses were not conducted in this study but will be addressed in future work.

N-glycosylation of GP5, predicted using NetNGlyc [10], revealed nine potential glycosylation sites (N30, N32, N33, N34, N35, N44, N50, N51, N57) across the 24 strains, forming nine distinct glycosylation patterns (A–I). Glycosylation at neutralization-associated sites (N33, N44, N51) was common, and a previously unreported site at N57 was observed. Strains 40 and 41 displayed the highest number of potential glycosylation sites, consistent with findings from Chen [4] and Jacab [13]. High levels of N-glycosylation can shield neutralizing epitopes and reduce immunogenicity, potentially enabling immune evasion and vaccine escape [1,36]. However, functional studies are required to determine the impact of glycosylation on epitope exposure, immune recognition, and viral pathogenesis.

Recombination analysis identified a statistically robust event in strain 42_Montana2019 (lineage 1A) between nucleotide positions 407–544, corresponding to the mid–C-terminal region of ORF5, which encompasses functionally relevant GP5 domains. The major and minor parental sequences were inferred as MH719138_1 (lineage 1A, Peru-2016/5) and 36_Montana2019 (lineage 1A), respectively. While overall nucleotide similarity with the major parent was ∼95.3%, the recombinant segment (407–544 nt) displayed higher phylogenetic affinity to the minor parent, supporting genomic fragment exchange [24,33].

UPGMA trees constructed for non-recombinant (1–406 nt) and recombinant (407–544 nt) regions demonstrated clear topological incongruence: in the 1–406 region, 42_Montana2019 clustered with the major parent, whereas in the 407–544 region, it grouped with 36_Montana2019, consistent with a mosaic genome structure and intra-lineage homologous recombination. Although homoplasy and convergent mutation cannot be fully excluded, the consistent detection across seven independent algorithms, well-defined breakpoints, and topological evidence strongly support a genuine recombination event.

Genetic diversity analysis revealed high variability within the Lima cohort, with 530 polymorphic nucleotide sites. Tajima’s D value (−0.78746) suggests a significant incidence of population subdivision or purifying selection [27,34,39].

Overall, these findings indicate substantial GP5 genetic diversity in circulating PRRSV-2 strains in Lima, highlighting the need for broader and temporally resolved sampling to assess national evolutionary patterns. This variability may partly explain suboptimal vaccine compatibility, though vaccine efficacy was not evaluated in this study.

## CONCLUSIONS

✓ The Nextclade v3 server successfully resolved the phylogenetic relationships of the 24 study strains within lineage 1, sublineage 1A (NADC34), revealing high divergence in strains 36, 37, 38, 39, and 42 based on GP5 sequence alignment.
✓ Amino acid substitution analysis using MEGA6 identified significant changes in key antigenic regions (signal peptide, decoy epitope I [A], HVR1, HVR2, primary neutralizing epitope B, and neutralizing epitope C) associated with both neutralizing and non-neutralizing antibody responses: N32 S/G/R/E (16/24), S34 N/T (3/24), S35 N/I (6/24), L39 F (7/24), Q40 L/R (6/24), L41 Y/V (3/24), K58 E/V/R (15/24), and S59 H/R/N (8/24) within epitopes A, B, and C of GP5.
✓ NetNGlyc analysis identified nine N-glycosylation patterns (A–I) across the 24 strains, with nine putative glycosylation sites at N30, N32, N33, N34, N35, N44, N50, N51, and N57. Patterns A, B, E, and G exhibited 5–6 glycosylation sites in 12/24 strains.
✓ Recombination analysis using RDP v4.101 detected a statistically robust event in strain 42_Montana2019 (lineage 1A). The inferred major parent was MH719138_1 (lineage 1A, Peru-2016/5; 95.3% similarity), while the minor parent was undetermined but showed highest phylogenetic affinity with 36_Montana2019.
✓ Genetic diversity analysis of 24 ORF5 sequences (603 nt) using DnaSP v6 identified 530 polymorphic sites. Tajima’s D test indicated high genetic variability within the Lima cohort, with a D value of −0.78746.

## Supporting information

Archivos complementarios

## SUPPLEMENTARY MATERIAL

✓ **Supplementary Material 1:** ORF5 sequences of 24 study strains, 9 Peruvian reference strains, and 46 sequences from GenBank.
✓ **Supplementary Material 2:** ORF5 lineage 1A sequences used for RDP v4.101 recombination analysis.

## FUNDING

This research was conducted and financially supported at the facilities of the Animal Nutrition and Health Business Unit, Innovation and Development Area, Corporation Montana S.A., Lima, Peru.

## ACKNOWLEDGMENTS

The authors sincerely thank Montana S.A. for funding and logistical support, and Bach. Segundo Del Águila Soto for technical assistance and valuable suggestions throughout this study.

---

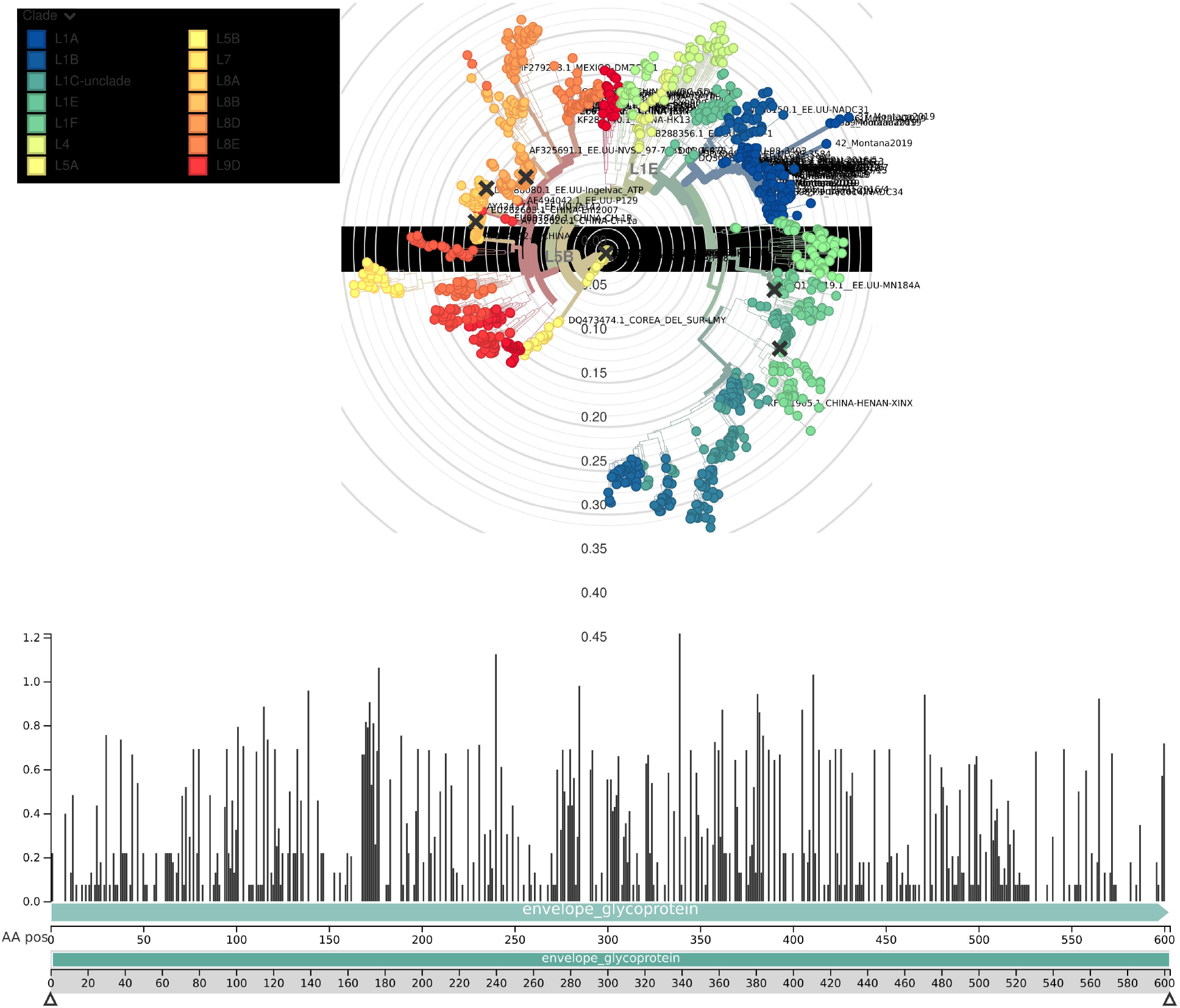

## Notes

### Competing Interest Statement

The authors have declared no competing interest.

### Summary of Updates

The article was improved thanks to the participation of two members, who provided valuable support in improving new techniques, writing, and structure.

## REFERENCES

[1] Ansari, I. H., Kwon, B., Osorio, F. A., & Pattnaik, A. K. (2006). Influence of N-linked glycosylation of porcine reproductive and respiratory syndrome virus GP5 on virus infectivity, antigenicity, and ability to induce neutralizing antibodies. Journal of Virology, 80(8), 3994–4004. 10.1128/JVI.80.8.3994-4004.2006

[2] Aksamentov, I., Roemer, C., Hodcroft, E. B., & Neher, R. A. (2021). Nextclade: Clade assignment, mutation calling, and quality control for viral genomes. Journal of Open Source Software, 6(67), 3773. 10.21105/joss.03773

[3] Brinton, M. A., Gulyaeva, A. A., Balasuriya, U., Dunowska, M., Faaberg, K. S., Goldberg, T., Leung, F., Nauwynck, H. J., Snijder, E. J., Stadejek, T., & Gorbalenya, A. E. (2021). ICTV virus taxonomy profile: Arteriviridae 2021. Journal of General Virology, 102(8), 001632. 10.1099/jgv.0.001632

[4] Chen, X. W., Li, L., Yin, M., Wang, Q., Luo, W. T., Ma, Y., Pu, Z. H., & Zhou, J. L. (2017). Cloning and molecular characterization of the ORF5 gene from a PRRSV-SN strain from Southwest China. Microbial Pathogenesis, 112, 295–302. 10.1016/j.micpath.2017.09.011

[5] Cotaquispe, N. R., De la Cruz, V. E., Quispe, C. E., Sitaras, N., & Toskano, H. J. (2022). Molecular characterization of porcine reproductive and respiratory syndrome (PRRS) based on the ORF7 nucleocapsid gene in swine farms in Lima, Peru. Revista de Investigaciones Veterinarias del Perú, 33, e21638. 10.15381/rivep.v33i6.21638

[6] Cotaquispe, N. R., Legua, B. M., Escajadillo, L. P., et al. (2025). Molecular characterization of the ORF5 (GP5) gene of the causative agent of porcine reproductive and respiratory syndrome (PRRS) detected in swine farms in Lima, Peru. Revista Argentina de Microbiología. 10.1016/j.ram.2025.07.005

[7] Faaberg, K. S., Hocker, J. D., Erdman, M. M., Harris, D. L., Nelson, E. A., Torremorell, M., & Plagemann, P. G. (2006). Neutralizing antibody responses of pigs infected with natural GP5 Nglycan mutants of porcine reproductive and respiratory syndrome virus. Viral Immunology, 19(2), 294–304. 10.1089/vim.2006.19.294

[8] Gibbs, M. J., Armstrong, J. S., & Gibbs, A. J. (2000). Sister-scanning: A Monte Carlo procedure for assessing signals in recombinant sequences. Bioinformatics, 16(7), 573–582. 10.1093/bioinformatics/16.7.573

[9] Guo, Z., Chen, X. X., Li, R., Qiao, S., & Zhang, G. (2018). The prevalent status and genetic diversity of porcine reproductive and respiratory syndrome virus in China: A molecular epidemiological perspective. Virology Journal, 15(1), 2. 10.1186/s12985-017-0910-6

[10] Gupta, R., & Brunak, S. (2002). Prediction of glycosylation across the human proteome and the correlation to protein function. In Pacific Symposium on Biocomputing (pp. 310–322). PMID: 11928486.

[11] Holtkamp, D. J., Kliebenstein, J. B., Neumann, E. J., Zimmerman, J. J., Rotto, H. F., Yoder, T. K., Wang, C., Yeske, P. E., Mowrer, C. L., & Haley, C. A. (2013). Assessment of the economic impact of porcine reproductive and respiratory syndrome virus on United States pork producers. Journal of Swine Health and Production, 21, 72–84.

[12] Hong, S., Wei, Y., Lin, S., He, W., Li, J., Chen, N., & Li, X. (2019). Genetic analysis of porcine reproductive and respiratory syndrome virus between 2013 and 2014 in southern parts of China: Identification of several novel strains with amino acid deletions or insertions in nsp2. BMC Veterinary Research, 15, 171. 10.1186/s12917-019-1906-9

[13] Jakab, S., Kaszab, E., Marton, S., Bányai, K., Bálint, Á., Nemes, I., & Szabó, I. (2022). Genetic diversity of imported PRRSV-2 strains, 2005–2020, Hungary. Frontiers in Veterinary Science, 9, 986850. 10.3389/fvets.2022.986850

[14] Jantafong, T., Karnbunchob, N., Tanomsridachchai, W., Mutthi, P., & Pavasutthipaisit, S. (2025). Nested open reading frame (ORF) 7 reverse transcription polymerase chain reaction and ORF5 phylogenetic refinement for enhanced detection and genetic classification of porcine reproductive and respiratory syndrome virus-2 in Thailand. Veterinary World, 18(9), 2850–2866.

[15] Jeong, H., Eo, Y., Lee, D., Jang, G., Min, K.-C., Choi, A. K., Won, H., Cho, J., Kang, S. C., & Lee, C. (2025). Comparative genomic and biological investigation of NADC30- and NADC34-like PRRSV strains isolated in South Korea. Transboundary and Emerging Diseases, 72, Article 9015349. 10.1155/tbed/9015349

[16] Jones, D. T., Taylor, W. R., & Thornton, J. M. (1992). The rapid generation of mutation data matrices from protein sequences. Computer Applications in the Biosciences, 8, 275–282.

[17] Lam, H. M., Ratmann, O., & Boni, M. F. (2017). Improved algorithmic complexity for the 3SEQ recombination detection algorithm. Molecular Biology and Evolution, 35(1), 247–251. 10.1093/molbev/msx263

[18] Martin, D., & Rybicki, E. (2000). RDP: Detection of recombination among aligned sequences. Bioinformatics, 16(6), 562–563. 10.1093/bioinformatics/16.6.562

[19] Martin, D., Posada, D., Crandall, K., & Williamson, C. (2005). A modified Bootscan algorithm for automated identification of recombinant sequences and recombination breakpoints. AIDS Research and Human Retroviruses, 21(2), 98–102. 10.1089/aid.2005.21.98

[20] Martin, D. P., Murrell, B., Golden, M., Khoosal, A., & Muhire, B. (2015). RDP4: Detection and analysis of recombination patterns in virus genomes. Virus Evolution, 1, vev003. 10.1093/ve/vev003

[21] Martin, D. P., Varsani, A., Roumagnac, P., Botha, G., Maslamoney, S., Schwab, T., Kelz, Z., Kumar, V., & Murrell, B. (2020). RDP5: A computer program for analyzing recombination in nucleotide sequence datasets and removing recombination signals. Virus Evolution, 7, veaa087. 10.1093/ve/veaa087

[22] Nguyen, N. H., Tran, H., Nguyen, T. Q., Nguyen, P., Le, T., Lai, D. C., & Nguyen, M. N. (2022). Phylogenetic analysis of porcine reproductive and respiratory syndrome virus in Vietnam, 2021. Virus Genes, 58(4), 361–366. 10.1007/s11262-022-01912-w

[23] Padidam, M., Sawyer, S., & Fauquet, C. M. (1999). Possible emergence of new geminiviruses by frequent recombination. Virology, 265(1), 218–225. 10.1006/viro.1999.0056

[24] Pellegrini Ferreira, C., Galina-Pantoja, L., Wagner, M., & Schroeder, D. C. (2025). Recombinants are the key drivers of recent PRRSV-2 evolution. Pathogens, 14(8), 743. 10.3390/pathogens14080743

[25] Petersen, T. N., Brunak, S., von Heijne, G., & Nielsen, H. (2011). SignalP 4.0: Discriminating signal peptides from transmembrane regions. Nature Methods, 8(10), 785–786. 10.1038/nmeth.1701

[26] Posada, D. (2002). Evaluation of methods for detecting recombination from DNA sequences: Empirical data. Molecular Biology and Evolution, 19(5), 708–717. 10.1093/oxfordjournals.molbev.a004129

[27] Rozas, J., Ferrer-Mata, A., Sánchez-DelBarrio, J. C., Guirao-Rico, S., Librado, P., Ramos-Onsins, S. E., & Sánchez-Gracia, A. (2017). DnaSP 6: DNA sequence polymorphism analysis of large data sets. Molecular Biology and Evolution, 34(12), 3299–3302. 10.1093/molbev/msx248

[28] Shin, G., Park, J., Lee, K., Ko, M., Ku, B., Park, C., & Jeoung, H. (2021). Genetic diversity of porcine reproductive and respiratory syndrome virus and evaluation of three one-step real-time RT-PCR assays in Korea. Research Square, 1–20. 10.21203/rs.3.rs-1149203/v1

[29] Smith, J. M. (1992). Analyzing the mosaic structure of genes. Journal of Molecular Evolution, 34(2), 126–129. 10.1007/BF00182389

[30] Tamura, K., Stecher, G., Peterson, D., Filipski, A., & Kumar, S. (2013). MEGA6: Molecular Evolutionary Genetics Analysis version 6.0. Molecular Biology and Evolution, 30(12), 2725–2729.

[31] Tajima, F. (1989). Statistical method for testing the neutral mutation hypothesis by DNA polymorphism. Genetics, 123(3), 585–595. PMC1203831

[32] Tajima, F. (1993). Statistical analysis of DNA polymorphism. Idengaku Zasshi, 68(6), 567–595. 10.1266/jjg.68.567

[33] Wang, A., Chen, Q., Wang, L., Madson, D., Harmon, K., Gauger, P., … Li, G. (2019). Recombination between vaccine and field strains of porcine reproductive and respiratory syndrome virus. Emerging Infectious Diseases, 25(12), 2335–2337. 10.3201/eid2512.191111

[34] Wang, T., Wang, X. A., Zhang, J. Q., Li, Y., Liu, H., Chen, R., & Zhao, K. (2025). Molecular characterization of porcine reproductive and respiratory syndrome virus in Henan and Shanxi, China, during 2023–2024. Archives of Virology, 170, 206. 10.1007/s00705-025-06380-9

[35] Yim-im, W., Anderson, T. K., Paploski, I. A. D., VanderWaal, K., Gauger, P., Krueger, K., Shi, M., Main, R., & Zhang, J. (2023). Refining PRRSV-2 genetic classification based on global ORF5 sequences and investigation of their geographic distributions and temporal changes. Microbiology Spectrum, 11, e02916–23. 10.1128/spectrum.02916-23

[36] Zhang, L., Feng, Y., Martin, D. P., Chen, J., Ma, S., Xia, P., & Zhang, G. (2017). Genetic diversity and phylogenetic analysis of the ORF5 gene of PRRSV from central China. Research in Veterinary Science, 115, 226–234.

[37] Zeller, M. A., Chang, J., Trevisan, G., Main, R. G., Gauger, P. C., & Zhang, J. (2024). Rapid PRRSV-2 ORF5-based lineage classification using Nextclade. Frontiers in Veterinary Science, 11, 1419340. 10.3389/fvets.2024.1419340

[38] Zheng, Y., Li, G., Luo, Q., Sha, H., Zhang, H., Wang, R., Kong, W., Liao, J., & Zhao, M. (2024). Research progress on the N protein of porcine reproductive and respiratory syndrome virus. Frontiers in Microbiology, 15, 1391697. 10.3389/fmicb.2024.1391697

[39] Zhou, L., Kang, R., Ji, G., Tian, Y., Ge, M., Xie, B., Yang, X., & Wang, H. (2018). Molecular characterization and recombination analysis of porcine reproductive and respiratory syndrome virus emerged in southwestern China during 2012–2016. Virus Genes, 54(1), 98–110. 10.1007/s11262-017-1519-y

