## Supplementary material for "Genetic Variability, N-Glycosylation Sites, and Recombination Events in the ORF5 (GP5) Gene of Lineage 1A (NADC34) *Betaarterivirus americense* Strains from Lima, Peru Molecular Variability of the GP5 Glycoprotein in *Betaarterivirus americense*": Archivos complementarios: Supplementary Material 1. ORF5 sequences of 24 study strains, 9 Peruvian reference strains, and 46 sequences from GenBank..pdf

>MF326985.1_IA/2014/NADC34

ATGTTGGGGAAATGCTTGACCGCGGGCTGCTGCTCGCAATTGCCTTTTTTGTGGTGTAT
CGTGCCGTTCTGTTTTGTTGCGCTCGTCAACGCCAACAACAGCAGCAGCTCCCATTTAC
AGTTGATTTATAACCTGACGATATGTGAGCTGAATGGCACAGATTGGCTAAATAAAAG
TTTTGATTGGGCGGTGGAGACCTTTGTTATTTTTCCTGTGTTGACTCATATTGTCTCCTA
TGGCGCCCTCACCACCAGCCATTTCCTTGACACAGTCGGCCTGATCACCGTGTCTGCCG
CCGGATATTACCACGGGCGGTATGTCTTGAGTAGCATTTACGCCGTCTGCGCCCTGGCT
GCGTTAACTTGCTTCGTCATCAGGCTAACAAAAAATTGTATGTCCTGGCGTTACTCATG
CACCAGATATACTAACTTTCTTCTGGACACCAAGGGCAAACTCTATCGTTGGCGGTCTC
CTGTCATCATAGAGAAAGGGGGTAAAATTGAGGTCGAAGGTCACCTGATCGACCTCAA
GAGAGTTGTGCTTGACGGTTCCGCGGCAACCCCTGTAACCAAAGTTTCAGCGGAACAA
TGGGGTCGTCCTTAG

>EF536003.1_EE.UU-VR2332

ATGTTGGAGAAATGCTTGACCGCGGGCTGTTACTCGCAATTGCTTTCTTTGTGGTGTAT
CGTGCCGTTCTGTTTTGCTGTGCTCGTCAACGCCAGCAACGACAGCAGCTCCCATCTAC
AGCTGATTTACAACTTGACGCTATGTGAGCTGAATGGCACAGATTGGCTAGCTAACAA
ATTTGATTGGGCAGTGGAGAGTTTTGTCATCTTTCCCGTTTTGACTCACATTGTCTCCTA
TGGTGCCCTCACTACTAGCCATTTCCTTGACACAGTCGCTTTAGTCACTGTGTCTACCGC
CGGGTTTGTTCACGGGCGGTATGTCCTAAGTAGCATCTACGCGGTCTGTGCCCTGGCTG
CGTTGACTTGCTTCGTCATTAGGTTTGCAAAGAATTGCATGTCCTGGCGCTACGCGTGT
ACCAGATATACCAACTTTCTTCTGGACACTAAGGGCAGACTCTATCGTTGGCGGTCGCC
TGTCATCATAGAGAAAAGGGGCAAAGTTGAGGTCGAAGGTCATCTGATCGACCTCAAA
AGAGTTGTGCTTGATGGTTCCGTGGCAACCCCTATAACCAGAGTTTCAGCGGAACAAT
GGGGTCGTCCTTAG

>AF066183.4_EE.UU-RespPRRS_MLV

ATGTTGGAGAAATGCTTGACCGCGGGCTGTTGCTCGCAATTGCTTTCTTTGTGGTGTAT
CGTGCCGTTCTGTTTTGCTGTGCTCGCCAACGCCAGCAACGACAGCAGCTCCCATCTAC
AGCTGATTTACAACTTGACGCTATGTGAGCTGAATGGCACAGATTGGCTAGCTAACAA
ATTTGATTGGGCAGTGGAGAGTTTTGTCATCTTTCCCGTTTTGACTCACATTGTCTCCTA
TGGTGCCCTCACTACCAGCCATTTCCTTGACACAGTCGCTTTAGTCACTGTGTCTACCGC
CGGGTTTGTTCACGGGCGGTATGTCCTAAGTAGCATCTACGCGGTCTGTGCCCTGGCTG
CGTTGACTTGCTTCGTCATTAGGTTTGCAAAGAATTGCATGTCCTGGCGCTACGCGTGT
ACCAGATATACCAACTTTCTTCTGGACACTAAGGGCGGACTCTATCGTTGGCGGTCGCC
TGTCATCATAGAGAAAAGGGGCAAAGTTGAGGTCGAAGGTCATCTGATCGACCTCAAA

AGAGTTGTGCTTGATGGTTCCGTGGCAACCCCTATAACCAGAGTTTCAGCGGAACAAT
GGGGTCGTCCTTAG
>DQ988080.1_EE.UU-Ingelvac_ATP
ATGTTGGGGAGATGCTTGACCGCGGGCTGTTGCTCGCGATTGCTTTCTTTGTGGTGTAT
CGTGCCATTTTGTTTTGCTGCGCTCGTCAACGCCAACAGCAACAGCAGCTCTCATCTTC
AGTTAATTTACAACTTGACGCTATGTGAGCTGAATGGCACAGATTGGCTGAAAGACAA
ATTTGATTGGGCATTGGAGACTTTTGTCATCTTTCCCGTGTTGACTCACATTGTCTCATA
TAGTGCACTCACCACTAGCCATTTCCTTGACACAGTCGGTCTGGTTACTGTGTCTACTG
CCGGGTTCTACCACGGGCGGTATGTTCTGAGTAGCATCTACGCGGTCTGCGCTCTGGCC
GCATTGACTTGCTTCGTCATTAGGCTTGCGAAGAACTGCATGTCCTGGCGCTACTCTTG
TACCAGATATACTAACTTCCTTCTGGACACTAAGGGCAGACTCTATCGCTGGCGGTCGC
CCGTTATCATAGAGAAAGGGGGTAAGGTTGAGGTCGAAGGTCACCTGATCGACCTCAA
AAGAGTTGTGCTTGATGGTTCCGTGGCAACCCCTTTAACCAGAGTTTCAGCGGAACAAT
GGGGTCGTCTTTAG
>JN660150.1_EE.UU-NADC31
ATGTTGGGGAAATGCTCGACCGCGGGCTGCTGCTCGCCATTGCTTTTTTTGTGGTGTAT
CGTGCCGTCCTGTTTAGTTGCGCTCGTCAACGCCAGCAAAAACAACAGCTCCCATTTAC
AGTCGATTTATAACCTGACGATATGTGAGCTGAATGGCACAGATTGGCTAAATAAAAA
ATTTGACTGGGCAGTGGAGACCTTTGTCATTTTTCCTGTATTGACTCATATCGTCTCCTA
TGGTGCCCTCACCACCAGCCATTTCCTTGACGCAGTCGGTTTGGTCATCGTGTCCACCG
CCGGATACTATCACGGGCGGTATGTCCTGAGCAGCATTTACGCTGTCTGCGCCTTGGCC
GCGCTGATTTGCTTCGCCATCAGGTTAACGAAAAACTGCATGTCCTGGCGCTACTCATG
TACTAGGTATACTAACTTTCTTCTAGACACCAAGGGCAAACTCTATCGTTGGCGGTCTC
CCGTCATCATAGAAAAAAAAGGGAAAATCGAGGTCAACGGTCACTTGATCGACCTCAA
GAGAGTTGTGCTTGATGGTTCCGCAGCAACCCCTGTAACCAAAGTTTCAGCGGAACAA
TGGGGTCATCCTTAG
>EF484033.1_EE.UU-pMLV
ATGTTGGAGAAATGCTTGACCGCGGGCTGTTGCTCGCAATTGCTTTCTTTGTGGTGTAT
CGTGCCGTTCTGTTTTGCTGTGCTCGCCAACGCCAGCAACGACAGCAGCTCCCATCTAC
AGCTGATTTACAACTTGACGCTATGTGAGCTGAATGGCACAGATTGGCTAGCTAACAA
ATTTGATTGGGCAGTGGAGAGTTTTGTCATCTTTCCCGTTTTGACTCACATTGTCTCCTA
TGGTGCCCTCACTACCAGCCATTTCCTTGACACAGTCGCTTTAGTCACTGTGTCTACCGC
CGGGTTTGTTCACGGGCGGTATGTCCTAAGTAGCATCTACGCGGTCTGTGCCCTGGCTG
CGTTGACTTGCTTCGTCATTAGGTTTGCAAAGAATTGCATGTCCTGGCGCTACGCGTGT

ACCAGATATACCAACTTTCTTCTGGACACTAAGGGCGGACTCTATCGTTGGCGGTCGCC
TGTCATCATAGAGAAAAGGGGCAAAGTTGAGGTCGAAGGTCATCTGATCGACCTCAAA
AGAGTTGTGCTTGATGGTTCCGTGGCAACCCCTATAACCAGAGTTTCAGCGGAACAAT
GGGGTCGTCCTTAG
>U87392.3_EE.UU-VIRUS2
ATGTTGGAGAAATGCTTGACCGCGGGCTGTTGCTCGCGATTGCTTTCTTTGTGGTGTAT
CGTGCCGTTCTGTTTTGCTGTGCTCGCCAACGCCAGCAACGACAGCAGCTCCCATCTAC
AGCTGATTTACAACTTGACGCTATGTGAGCTGAATGGCACAGATTGGCTAGCTAACAA
ATTTGATTGGGCAGTGGAGAGTTTTGTCATCTTTCCCGTTTTGACTCACATTGTCTCCTA
TGGTGCCCTCACTACCAGCCATTTCCTTGACACAGTCGCTTTAGTCACTGTGTCTACCGC
CGGGTTTGTTCACGGGCGGTATGTCCTAAGTAGCATCTACGCGGTCTGTGCCCTGGCTG
CGTTGACTTGCTTCGTCATTAGGTTTGCAAAGAATTGCATGTCCTGGCGCTACGCGTGT
ACCAGATATACCAACTTTCTTCTGGACACTAAGGGCAGACTCTATCGTTGGCGGTCGCC
TGTCATCATAGAGAAAAGGGGCAAAGTTGAGGTCGAAGGTCATCTGATCGACCTCAAA
AGAGTTGTGCTTGATGGTTCCGTGGCAACCCCTATAACCAGAGTTTCAGCGGAACAAT
GGGGTCGTCCTTAG
>MF279253.1_MEXICO-DMZC181
ATGTTGGGGAATTGCTTGACAGCTGTTTGTTGCTCGCGATTGTTTTCTTTGTGGTATATC
GTGCCGTTTTGTTTTGCTGTGCTCGTCAACGCCAACARCAGCAACAGCTCTCATTTTCA
GTTGATTTATAACTTGACGATATGCGAGCTGAATGGCACAGACTGGCTGAGCCACAAC
TTTGATTGGGCCGTGGAGACTTTTGTCATTTTTCCCGTGCTAACTCACATCGTTTCCTAT
GGTGCACTCACCACCAGCCACTTCCTCGACACGGTTGGTCTGGTTACAGTGTCCACCGC
CGGGTATTACCACGGGCGGTATGTCTTGAGCAGCATCTATGCAATCTGTGCCTTAGCAG
CATTGACTTGCTTCTTCATTAGGCTTACGAAAAATTGCATGTCCTGGCGTTACTCTTGTA
CCAGATATACCAACTTCCTTCTAGACACTAAGGGCAGGCTCTATCGTTGGCGGTCTCCC
GTCATCATAGAGAAAGGGGGTAAGGTTGAGGTTGAAGGTCACCTAATCGACCTCAAAA
GAGTCGTGCTTGATGGTTCCGCGGCAACYCCTTTAACCAAAGTTTCAGCGGAGCAATG
GGGTCGTCCCTAG
>KF611905.1_CHINA-HENAN-XINX
ATGTTGGTGAAATGCTTGGCCGCGGGTTGTTGCTCGCAATTGCCTTTTTTGTGGTGTATC
GTGCCGTTCTATTTTGCTGCGCTCGTCAACGCCAACAGCAACAGCAGCTCCCATCTACA
GTTGATTTATAACCTGACGATATGTGAGCTGAATGGTACGGATTGGCTGGGATCAAATT
TTGATTGGGCAGTGGAGACTTTCGTCATCTTTCCTGTATTGACTCATATTGTCTCTTACG
GCGCCCTTACCACTAGCCATTTTCTTGACACGGTCGGCCTGATCACTGTGTCCACCGCC

GGATATTTTCACAAGCGGTATGTGTTGAGTAGCATCTACGCTGTCTGTGCCCTGGCTGC
GTTGGTTTGCTTCGCCATTAGGTTGGCAAAAAATTGCATGTCCTGGCGCTACTCATGCA
CCAGATATACCAATTTTCTTCTGGACACTAAGGGCAAACTCTACCGCTGGCGGTCACCC
GTCATCATAGAGAAGGAGGGTAAAGTTGATGTGGGTGGTCACTTAATCGACCTCAAGA
GAGTTGTGCTTGATGGTTCCGCGGCAACCCCTGTAACCAAGATTTCAGCGGAACAATG
GGGTCGTCCATAG

>KF287140.1_CHINA-HK13

ATGTTGGGGAAATGCTTGACCGCGGGCTGTTGCTCGCGATTGCTTTCTTTGTGGTGTAT
CGTGCCGTTCTATCTTGCTGTGCTCGTCAACGCCAACAACAACAACAGCTCTCATATTC
AGTTGATTTATAACTTGACGCTATGTGAGCTGAATGGCACAGATTGGCTGGCACAAAA
ATTTGACTGGGCAGTGGAGACTTTTGTCATCTTCCCCGTGTTGACTCACATTGTTTCCTA
TGGGGCACTCACCACCAGCCATTTCCTTGACACAGTTGGTCTGGTCACTGTGTCCACCG
CCGGATATTATCACAGGCGGTATGTCTTGAGTAGCATTTACGCAGTCTGTGCTCTGGCT
GCGCTGATTTGCTTTGTCATTAGGCTTGCGAAGAACTGCATGTCCTGGCGCTACTCTTG
TACCAGATATACCAACTTCCTTCTGGACACTAAGGGCAGACTCTATCGTTGGCGGTCGC
CCGTCATTGTGGAGAAAGGGGGTAAGGTTGAGGTCGAAGGTCACCTGATCGACCTCAA
GAGAGTTGTGCTTGATGGTTCCGTGGCAACCCCTTTAACCAGAGTTTCAGCGGAACAAT
GGGGTCGTCTCTAG

>KC862577.1_DINAMARCA-DK-1

ATGTTGGAGAAATGCTTGACCGCGGGCTGTTGCTCGCAATTGCTTTCTTTGTGGTGTAT
CGTGCCGTTCTGTTTTGCTGTGCTCGCCAACGCCAGCAACGACAACAGCTCCCATCTAC
AGCTGATTTACAACTTGACGCTATGTGAGCTGAATGGCACAGATTGGCTAGCTAACAA
ATTTGATTGGGCAGTGGAGAGTTTTGTCATCTTTCCCGTTTTGACTCACATTGTTTCCTA
TGGTGCCCTCACTACTAGCCATTTTCTTGACACAGTCGCTTTAGTCACTGTGTCTACCGC
CGGGTTTGTTCACGGGCGGTATGTCCTAAGTAGCATCTACGCGGTCTGTGCCCTGGCTG
CGTTGACTTGCTTCGTCATTAGGTTTGCAAAGAATTGCATGTCCTGGCGCTACGCGTGT
ACCAGATATACCAACTTTCTTCTGGACACTAAGGGCAGACTCTATCGTTGGCGGTCGCC
TGTCATCATAGAGAAAAGGGGCAAAGTTGAGGTCGAAGGTCATCTGATCGACCTCAAA
AGAGTTGTGCTTGATGGTTCCGTGGCAACCCCTATAACCAGAGTTTCAGCGGAACAAT
GGGGTCGTCCTTAG

>JX512910.2_EE.UU-SRV07

ATGTTGGGGAAGTGCTTGACCGCGTGCTGTTGCTCGCGATTGCTTTTTTTGTGGTGTATC
GTGCCGTTCTATCTTGCTGTGCTCGTCAACGCCAGCAACAACAACAGCCCTCATATTCA
GTTGATTTATAACTTAACGCTATGTGAGCTGAATGGCACAGATTGGCTGGCACAAAAA

TTTGACTGGGCAGTGGAGACTTTTGTCATCTTCCCCGTGTTGACTCACATTGTTTCCTAT
GGGGCACTCACCACCAGCCATTTCCTTGACACAGTTGGTCTGGCCACTGTGTCCACCGC
CGGATATTATCACGGGCGGTATGTCTTGAGTAGCATTTACGCAGTCTGTGCTCTGGCTG
CGCTGATTTGCTTTGTCATTAGGCTTGCGAAGAACTGCATGTCCTGGCGCTACTCTTGT
ACCAGATATACCAACTTCCTTCTGGACACTAAGGGCAGACTCTATCGTTGGCGGTCGCC
CGTCATTGTGGAGAAAGGGGGTAAGGTTGAGGTCGAAGGTCACCTCATCGACCTCAAA
AGAGTTGTGCTTGATGGTTCCGCGGCAACCCCTTTAACCAGAGTTTCAGCGGAACAATG
GGGTCGTCTCTAG

>JQ715697.1_CHINA-NVDC-GD2

ATGTTGGGGAAGTGCTTGACCGCGTGCTGTTGCTCACGATTGCTTTTTTTGTGGTGTATC
GTGCCGTTCTATCTTGCTGTGCTCGTCAACGCCAGCAACAACAACAGCTCTCATATTCA
GTTGATTTATAACTTAACGCTATGTGAGCTGAATGGCACAGATTGGCTGGCACAAAAA
TTTGACTGGGCAGTGGAGACTTTTGTCATCTTCCCCGTTTTGACTCACATTGTTTCCTAT
GGGGCACTCACTACCAGCCATTTCCTTGACACAGTTGGTCTGGCCACTGTGTCCACCGC
TGGATATTATCACAAGCGGTATGTCTTGAGTAGCATTTACGCAGTCTGTGCTCTGGCTG
CGCTGATTTGCTTTGTCATTAGGCTTGCGAAGAACTGCATGTCCTGGCGCTACTCTTGT
ACCAGATATACCAACTTCCTCCTGGACACTAAGGGCAGACTCTATCGTTGGCGGTCGCC
CATCATTGTAGAGAAGGGGGGTAAGGTTGAGGTCGAAGGTCACCTGATCGACCTCAAG
AGAGTTGTGCTTGATGGTTCCGTGGCAACCCCTTTAACCAGAGTTTCAGCGGAACGATG
GGGTCGTCTATAG

>JQ663556.1_CHINA-10-10QN

ATGTTGGGGAAGTGCTTGACCGCGTGCTGTTGCTCGCGATTGCTTTTTTTGTGGTGTATC
GTGCCGTTCTATCTTGCTGTGCTCGCCAACGCCAGCAACAGCAACAGCTCTCATATTCA
GTTGATTTATAACTTAACGCTATGTGAGCTGAATGGCACAGATTGGCTGGCACAAAAA
TTTGACTGGGCAGTGGAGACTTTTGTCATTTTCCCCGTGTTGACTCACATTGTTTCCTAT
GGAGCACTCACCACCAGCCATTTCCTTGACACAGTTGGTCTAGCCACTGTGTCCACCGC
CGGATATTATCACGGGCGGTATGTCTTGAGTAGCATTTACGCAGTCTGTGCTCTGGCTG
CGCTGATTTGCTTTGTCATTAGGCTTGCGAAGAACTGCATGTCCTGGCGCTACTCTTGT
ACCAGATACACCAACTTCCTTCTGGACACTAAGGGCAGACTCTATCGTTGGCGGTCACC
CGTCATTGTGGAGAAAGGGGGTAAGGTTGAGGTCGGAGGTCACCTGATCGACCTCAAG
AGAGTTGTGCTTGATGGTTCCGCGGCAACCCCTTTAACCAGAGTTTCAGCGGAACTATG
GGGTCGTCTCTAG

>JQ663545.1_CHINA-10-10BJ-5

ATGTTGGGGAAATGCTTGACCGCGTGCTGTTGCTCGCGATTGCTTTTTTTGTGGTGTATC
GTGCCGTTCTATCTTGCTGTGCTCGCCAACGCCAGCAACAGCAACAGCTCTCATATTCA
GTTGATTTATAACTTAACGCTATGTGAGCTGAATGGCACAGATTGGCTGGCACAAAAA
TTTGACTGGGCAGTGGAGACTTTTGTCATCTTCCCCGTGTTGACTCACATTGTTTCCTAT
GGAGCACTCACCACCAGCCATTTCCTTGACACAGTTGGTCTAGCCACTGTGTCCACCGC
CGGATATTATCACGGGCGGTATGTCTTGAGTAGCATTTACGCAGTCTGTGCTCTGGCTG
CGCTGATTTGCTTTGTCATTAGGCTTGCGAAGAACTGCATGTCCTGGCGCTACTCTTGT
ACCAGATATACCAACTTCCTTCTGGACACTAAGGGCAGACTCTATCGTTGGCGGTCACC
CGTCATTGTGGAGAAAGGGGGTAAGGTTGAGGTCGAAGGTCACCTGATCGACCTCAAG
AGAGTTGTGCTTGATGGTTCCGCGGCAACCCCTTTAACCAGAGTTTCAGCGGAACAATG
GGGTCGTCTTTAG
>HQ315837.1_CHINA-SY0909
ATGTTGGGGAAGTGCTTGACCGCGTGCTGTTGCTCGCGATTGCCTTTTTTGTGGTGTATC
GTGCCGTTCTATCTTGCTGTGCTCGCCAACGCCAGCAACAACAACAGCTCTCATATTCA
GTTGATTTATAACTTAACGCTATGTGAGCTGAATGGCACAGATTGGCTGGCACAAAAA
TTTGACTGGGCAGTGGAGACTTTTGTCATCTTCCCCGTGTTGACTCACATTGTTTCCTAT
GGGGCACTCACCACCAGCCATTTCCTTGACACAGTTGGTTTGGCCACTGTGTCCACCGC
CGGATATTATCACGGGCGGTATGTCTTGAGTAGCATTTACGCGGTCTGTGCTCTGGCTG
CGCTGATTTGCTTTGTCATTAGGCTTGCGAAGAACTGCATGTCCTGGCGCTACTCTTGC
ACCAGATATACCAACTTCCTTCTGGACACTAAGGGCAGACTCTATCGTTGGCGGTCGCC
CGTCATTGTGGAGAAAGGGGGTAAGGTTGAGGTCGAAGGTCACCTGATCGACCTCAAG
AGAGTTGTGCTTGATGGTTCCGCGGCAACCCCTTTAACCAGAGTTTCAGCGGAACAATG
GGGTCGTCTCTAG
>GQ857656.1_CHINA-SX-1
ATGTTGGGGAAGTGCTTGACCGCGTGCTGTTGCTCGCGATTGCTTTTTTTGTGGTGTATC
GTGCCGTTCTATCTTGCTGTGCTCGCCAACGCCAGCAACAACAACAGCTCTCATATTCA
GTTGATTTATAACTTAACGCTATGTGAGCTGAATGGCACAGATTGGCTGGCACAAAAA
TTTGACTGGGCAGTGGAGACTTTTGTCATCTTCCCCGTGTTGACTCACATTGTTTCCTAT
GGAGCACTCACCACCAGCCATTTCCTTGACACAGTTGGTCTAGCCACTGTGTCTACCGC
CGGATATTATCACGGGCGGTATGTCTTGAGTAGCATTTACGCAGTCTGTGCTCTGGCTG
CGCTGATTTGCTTTGTCATTAGGCTTGCGAAGAACTGCATGTCCTGGCGCTACTCTTGT
ACCAGATATACCAACTTCCTTCTGGACACTAAGGGCAGACTCTATCGTTGGCGGTCACC
CGTCATTGTAGAGAAAGGGGGTAAGGTTGAGGTCGAAGGTCACCTGATCGACCTCAAG

AGAGTTGTGCTTGATGGTTCCGCGGCAACCCCTTTAACCAGAGTTTCAGCGGAACAATG
GGGTCGTCTCTAG
>GQ374442.1_CHINA-GDBY1
ATGTTGGGGAAGTGCTTGACCGCGTGCTGTTGCTCGCGATTGCTTTTTTTGTGGTGTATC
GTGCCGTTCTATCTTGCTGTGCTCGTCAACGCCAGCAACAACAACAGCTCTCATATTCA
GTTGATTTATAACTTGACGCTATGTGAGCTGAATGGCACAGATTGGCTGGCACAAAAA
TTTGACTGGGCAGTGGAGACTTTTGTCATCTTCCCCGTGTTGACTCACATTGTTTCCTAT
GGGGCACTCACCACCAGCCATTTCCTTGACACAGCTGGTCTGGCCACTGTGTCCACCGC
CGGATATTATCACGGGCGGTATGTCTTGAGTAGCATTTACGCAGTCTGTGCTCTGGCTG
CGCTGATTTGCTTTGTCATTAGGCTTGCGAAGAACTGCATGTCCTGGCGCTACTCTTGT
ACCAGATATACCAACTTCCTTCTGGACACTAAGGGCAGACTCTATCGTTGGCGGTCGCC
CGTCATTGTGGAGAAAGGGGGTAAGGTTGAGGTCGAAGGTCACCTGATCGACCTCAAG
AGAGTTGTGCTTGATGGTTCCGCGGCAACCCCTTTAACCAGAGTTTCAGCGGAACAATG
GGGTCGTCTCTAG
>FJ889130.1_CHINA-CWZ-1-F3
ATGTTGGGGAAGTGCTTGACCGCGTGCTGTTGCTCGCGATTGCTTTTTTTGTGGTGTATC
GTGCCGTTCTATCTTGCTGTGCTCGTCAACGCCAGCAACAACAACAGCTCTCATATTCA
GTCGATTTATAACTTAACGTTATGTGAGCTGAATGGCACAGATTGGCTGGCACAAAAA
TTTGACTGGGCAGTGGAGACTTTTGTCATCTTCCCCGTGTTGACTCACATTGTTTCCTAT
GGGGCACTCACCACCAGCCATTTCCTTGACACAGTTGGTCTGGCCACTGTGTCCACCGC
CGGATATTATCACGGGCGGTATGTCTTGAGTAGCATTTACGCAGTCTGTGCTCTGGCTG
CGCTGATTTGCTTTGTCATTAGGCTTGCGAAGAACTGCATGTCCTGGCGCTACTCTTGT
ACCAGATATACCAACTTCCTTCTGGATACTAAGGGCAGACTCTATCGTTGGCGGTCGCC
CGTCATTGTGGAGAAAGGGGGTAAGGTTGAGGTCGAAGGTCACCTGATCGACCTCAAG
AGAGTTGTGCTTGATGGTTCCGCGGCAACCCCTTTAACCAGAGTTTCAGCGGAACAATG
GGGTCGTCTCTAG
>EU939312.1_CHINA-JSyx
ATGTTGGGGAAGTGCTTGACCGCGTGCTGTTGCTCGCGATTGCTTTTTTTGTGGTGTATC
GTGCCGTTCTATCTTGCTGTGCTCGTCAACGCCAGCAACAACAACAGCTCTCATATTCA
GTTGATTTATAACTTAACGCTATGTGAGCTGAATGGCACAGATTGGCTGGCACAAAAA
TTTGACTGGGCAGTGGAGACTTTTGTCATCTTCCCCGTGTTGACTCACATTGTTTCCTAT
GGGGCACTCACCACCAGCCATTTCCTTGACACAGTTGGTCTGGCCACTGTGTCCACCGC
CGGATATTATCACGGGCGGTATGTCTTGAGTAGCATTTACGCAGTCTGTGCTCTGGCTG
CGCTGATTTGCTTTGTCATTAGGCTTGCGAAGAACTGCATGTCCTGGCGCTACTCTTGT

ACCAGATATACCAACTTCCTTCTGGACACTAAGGGCAGACTCTATCGTTGGCGGTCGCC
CGTCATTGTGGAGAAAGGGGGTAAGGTTGAGGTCGAAGGTCACCTGATCGACCTCAAG
AGAGTTGTGCTTGATGGTTCCGCGGCAACCCCTTTAACCAGAGTTTCAGCGGAACAATG
GGGTCGTCTCTAG
>EU807840.1_CHINA-CH-1R
ATGTTGGGGAAATACTTGACCACGGGCTGCTGCTCGCGATTGCTTTCTTTGTGGTGTAT
CGTGCCGTTCTGTTTTGCTGTGCTCGTCAACGCCAACAGCAACAGCAGCTCTCAATTTC
AGTTGATTTATAACTTGACGCTATGTGAGCTGAATGGCACAGATTGGCTGGCTAACAA
ATTTGACTGGGCAGTGGAGACTTTTGTCATCTTTCCCGTGTTGACTCACATTGTGTCCTA
TGGGGCACTCACCACCAGCCATTTCCTTGACACAGTTGGTCTGGTCACTGTGTCCACCG
CCGGGTTTTATCACGGGCGGTATGTCTTGAGTAGCATCTACGCGGTCTGTGCTCTGGCT
GCGTTGATTTGCTTCGTCATTAGGCTTGCGAAGAACTGCATGTCCTGGCGCTACTCTTG
TACCAGATATACCAACTTCCTTCAGGACACTAAGGGCAGACTCTATCGTTGGCGGTCGC
CCGTTATTGTAGAGAAAGGGGGTAAGGTTGAGGTCGAGGGTCACCTGATCGACCTCAA
AAGAGTTGTGCTTGATGGTTCCGTGGCAACCCCTTTAACCAGAGTTTCAGCGGAACAAT
GGGGTCGTCTCTAG
>EU262603.1_CHINA-Em2007
ATGTTGGGGAAATGCTTGACCGCGGGCTGTTGCTCGCGATTGCCTTTTTTGTGGTGTAT
CGTGCCGTTCTGGTTTGTTGTGCTCGTCAACGCCAACAGAACCAGCAGTTCTCATTTTC
AGTTGATTTATAACTTGACGCTATGTGAGCTGAATGGCACAGATTGGCTGGCAGACAA
ATTTGACTGGGCAGTAGAGACTTTTGTCATCTTCCCCGTGTTGACTCACATTGTTTCCTA
TGGGGCACTCACCACCAGCCATTTCCTTGACACAGTTGGTCTGATCACAGTGTCCACCG
CCGGGTTTTACCATGGGCGGTATGTCTTGAGTAGCATCTACGCAGTCTGCGCTCTGGCT
GCGTTGATTTGCTTCATCATTAGGCTTGCGAAGAACTGCATGTCCTGGCGCTACTCTTG
CACCAGATATACCAACTTCCTTCTGGACACTAAGGGCAGACTCTATCGTTGGCGGTCGC
CCGTTATTGTGGAGAAAGGAGGTAAGGTTGAGGTCGAGGGTCACCTGATCGATCTCAA
AAGAGTTGTGCTTGATGGTTCTGCGGCAACCCCTTTAACCAGAGTTTCAGCGGAACAAT
GGGGTCGTCCTTAG
>EF635006.1_CHINA-HUN4
ATGTTGGGGAAGTGCTTGACCGCGTGCTGTTGCTCGCGATTGCTTTTTTTGTGGTGTATC
GTGCCGTTCTATCTTGCTGTGCTCGTCAACGCCAGCAACAACAACAGCTCTCATATTCA
GTTGATTTATAACTTAACGCTATGTGAGCTGAATGGCACAGATTGGCTGGCACAAAAA
TTTGACTGGGCAGTGGAGACTTTTGTCATCTTCCCCGTGTTGACTCACATTGTTTCCTAT
GGGGCACTCACCACCAGCCATTTCCTTGACACAGTTGGTCTGGCCACTGTGTCCACCGC

CGGATATTATCACGGGCGGTATGTCTTGAGTAGCATTTACGCAGTCTGTGCTCTGGCTG
CGCTGATTTGCTTTGTCATTAGGCTTGCGAAGAACTGCATGTCCTGGCGCTACTCTTGT
ACCAGATATACCAACTTCCTTCTGGACACTAAGGGCAGACTCTATCGTTGGCGGTCGCC
CGTCATTGTGGAGAAAGGGGGTAAGGTTGAGGTCGAAGGTCACCTGATCGACCTCAAG
AGAGTTGTGCTTGATGGTTCCGCGGCAACCCCTTTAACCAGAGTTTCAGCGGAACAATG
GGGTCGTCTCTAG
>EF112445.1_CHINA-JXA1
ATGTTGGGGAAGTGCTTGACCGCGTGCTGTTGCTCGCGATTGCTTTTTTTGTGGTGTATC
GTGCCGTTCTATCTTGCTGTGCTCGTCAACGCCAGCAACAACAACAGCTCTCATATTCA
GTTGATTTATAACTTAACGCTATGTGAGCTGAATGGCACAGATTGGCTGGCACAAAAA
TTTGACTGGGCAGTGGAGACTTTTGTCATCTTCCCCGTGTTGACTCACATTGTTTCCTAT
GGGGCACTCACCACCAGCCATTTCCTTGACACAGTTGGTCTGGCCACTGTGTCCACCGC
CGGATATTATCACGGGCGGTATGTCTTGAGTAGCATTTACGCAGTCTGTGCTCTGGCTG
CGCTGATTTGCTTTGTCATTAGGCTTGCGAAGAACTGCATGTCCTGGCGCTACTCTTGT
ACCAGATATACCAACTTCCTTCTGGACACTAAGGGCAGACTCTATCGTTGGCGGTCGCC
CGTCATTGTGGAGAAAGGGGGTAAGGTTGAGGTCGAAGGTCACCTGATCGACCTCAAG
AGAGTTGTGCTTGATGGTTCCGCGGCAACCCCTTTAACCAGAGTTTCAGCGGAACTATG
GGGTCGTCTCTAG
>DQ473474.1_COREA_DEL_SUR-LMY
ATGTTGGGGAAATGCTTGACCGCGGGCTGTTGCTCGCAATTGCTTTTTTTGTGGTGTAT
CGTGCCGTCTTGTTTTGTTGCGATCGTCAGCGCCAACAACAGCAGCAGCTCAAATTTAC
AGCTGATTTACAACTTGACGCTATGTGAGCTGAATGGCACAGATTGGCTAGCTAACAG
ATTTGACTGGGCGGTGGAGTGTTTTGTTATTTTTCCTGTATTGACTCACATTGTCTCTTA
TGGTGCCCTAACCACTAGCCACTTCCTTGACACAGTCGGTTTGGTCACTGTGTCTACCG
CCGGATTTGTTCACGGGCGGTATGTTTTGAGTAGCATTTACGCGGTCTGTGCCCTGGCT
GCGTTGATTTGCTTCGTCATTAGGCTTGCGAAGAATTGCATGTCCTGGCGCTACTCATG
TACCAGATATACCAACTTTCTTCTAGATACCAAGGGCAGACTCTACCGTTGGCGGTCGC
CTGTCATTATAGAGAAAAGGGGCAAAGTTGAGGTCGAGGGTCAACTAATCGACCCCAA
AAGAGTTGTGCTTGATGGTTCCCCGGCAACCCCTGTAACCAGAGTTTCAGCGGAACAA
TGGGGTCATCCTTAG
>DQ306879.1_EE.UU-99-3584
ATGTTGGGGAAATGCTTGACCGCGGGCTGTTGCTCGCAATTGCCTTTTTTGTGGTGTAT
CGTGCCGTTCTGTTTTGCTGCGCTCGTCAACGCCGGCAGCAACAGCAGCTCCCATTTAC
AGTTGATTTATAACCTGACGATATGTGAGCTGAATGGCACAGATTGGCTGAACGACAA

ATTTGATTGGGCGGTGGAGACTTTTGTCATCTTTCCTGTGTTGACTCACATTGTTTCMTA
TGGCGCCCTCACCACCAGCCATTTCCTTGACACAGTCGGTCTAGTTACTGTGTCTACCG
CCGGATATTACCATAGGCGGTATGTATTGAGTAGCATTTACGCTGTCTGTGCCCTGGCT
GCGTTGATTTGCTTCGTCATCAGGTTGACGAAGAATTGCATGTCCTGGCGCTACTCATG
CACCAGATATACTAACTTTCTTTTGGACACCAAGGGCAGACTCTATCGTTGGCGGTCAC
CCGTCATCATAGAGAAAGGGGGCAAAGTTGAGGTTGAAGGTCACCTGATCGACCTCAA
GAGAGTTGTGCTTGACGGTTCCGCGGCAACCCCCGTAACCAAAGTTTCAGCAGAACGA
TGGGGTCGTCCCTAG
>DQ306878.1_EE.UU-98-3403
ATGTTGGGGAAATGCTTGACCGCGGGCTGTTGCTCGCAATTGCCTTTTTTGTGGTGTAT
CGTGCCGTTCTGTTTTGCTGCGCTCGTCGACGCCAGCAGCAACAGCAGCTCCCATTTAC
AGTTGATTTATAACCTGACGATATGTGAGCTGAATGGCACAGATTGGCTGAACAACAA
ATTTGATTGGGCGGTGGAGACTTTTGTCATCTTTCCTGTGTTGACTCACATTGTTTCCTA
TGGCGCCCTCACCACCAGCCATTTCCTTGACACAGTCGGTCTGGTTACTGTGTCTACCG
CCGGATATTACCATGGGCGGTATGTATTGAGTAGCATTTACGCTGTCTGTGCCCTGGCT
GCGTTGATTTGCTTCGTCATTAGGTTGACGAAGAATTGCATGTCCTGGCGCTACTCATG
CACCAGATATACCAACTTTCTTTTGGACACCAAGGGCAGACTCTATCGTTGGCGGTCGC
CCGTCATCATAGAGAAAGGGGGCAAAGTTGAGGTTGAAGGTCACCTGATCGACCTCAA
GAGAGTTGTGCTTGACGGTTCCGCGGCAACCCCCGTAACCAAAGTTTCAGCAGAACGA
TGGGGTCGTCCCTAG
>DQ306877.1_EE.UU-98-3298
ATGTTGGGGAAATGCTTGACCGCGGGCTGTTGCTCGCAATTGCCTTTTTTGTGGTGTAT
CGTGCCGTTCTGTTTTGCTGCGCTCGTCAACGCCAGCAGCAACAGCAGCTCCCATTTAC
AGTTGATTTATAACCTGACGATATGTGAGCTGAATGGCACAGATTGGCTGAACGACAA
ATTTGATTGGGCGGTGGAGACTTTTGTCATCTTTCCTGTGTTGACTCACATTGTTTCCTA
TGGCGCCCTCACCACCAGCCATTTCCTTGACACAGTCGGTCTGGTTACTGTGTCTACCG
CCGGATATTACCATGGGCGGTATGTATTGAGTAGCATTTACGCTGTCTGTGCCCTGGCT
GCGTTGATTTGCTTCGTCATCAGGTTGACGAAGAATTGCATGTCCTGGCGCTACTCATG
CACCAGATATACCAACTTTCTTTTGGACACCAAGGGCAGACTCTATCGTTGGCGGTCAC
CCGTCATCATAGAGAAAGGGGGCAAAGTTGAGGTTGAAGGTCACCTGATCGACCTCAA
GAGAGTTGTGCTTGACGGTTCCGCGGCAACCCCCGTAACCAAAGTTTCAGCAGAACGA
TGGGGTCGTCCCTAG
>DQ176019.1__EE.UU-MN184A

ATGTTGGGGAAATGCTTGACCGCGGGCTATTGCTCGCAATTGCTTTTTTTGTGGTGTAT
CGTGCCGTTCTGTCTTGCTGCGCTCGTCAACGCCGACAGCAACAGCAGCTCCCATTTAC
AGTTGATTTATAAMTTAACGATATGTGAGCTGAATGGCACAGACTGGCTGAACAATCA
TTTTAGTTGGGCAGTGGAGACTTTCGTTATCTTTCCTGTGTTGACTCATATTGTTTCCTA
CGGCGCCCTCACTACCAGCCACCTCCTTGACACGGTCGGCCTGATCACTGTGTCCACCG
CCGGATACTGCCATAAGCGGTATGTCTTGAGTAGCATCTATGCTGTCTGCGCCCTGGCT
GCGCTGATTTGCTTCGTCATCAGGTTGACGAAAAATTGTATGTCCTGGCGCTACTCATG
TACCAGATATACCAACTTTCTTCTGGACACCAAGGGCAGACTCTATCGCTGGCGGTCAC
CCGTCATCATAGAGAAAAGGGGTAAAATTGAGGTTGGAGGTGACCTGATCGACCTCAA
GAGAGTTGTGCTTGATGGTTCCGCGGCAACCCCTGTAACCAAAGTTTCAGCGGAACAA
TGGGGTCGTCCTTAG
>AY424271.1_EE.UU-JA142
ATGTTGGGGAGATGCTTGACCGCGGGCTGTTGCTCGCGATTGCTTTCTTTGTGGTGTAT
CGTGCCATTTTGTTTTGCTGCGCTCGTCAACGCCAACAGCAACAGCAGCTCTCATCTTC
AGTTGATTTACAACTTGACGCTATGTGAGCTGAATGGCACAGATTGGCTGAAAGACAA
ATTTGATTGGGCAGTGGAGACTTTTGTCATCTTTCCCGTGTTGACTCACATTGTCTCATA
TGGTGCACTCACCACTAGCCATTTCCTTGACACAGTCGGTCTGGTTACTGTGTCTACCG
CCGGGTTCTACCACGGGCGGTATGTTCTGAGTAGCATCTACGCGGTCTGCGCTCTGGCC
GCATTGATTTGCTTCGTCATTAGGCTTGCGAAGAACTGCATGTCCTGGCGCTACTCTTG
TACCAGATATACTAACTTCCTTCTGGACACTAAGGGCAGACTCTATCGCTGGCGGTCGC
CCGTTATCATAGAGAAAGGGGGTAAGGTTGAGGTCGAAGGTCACCTGATCGACCTCAA
AAGAGTTGTGCTTGATGGTTCCGTGGCAACCCCTTTAACCAGAGTTTCAGCGGAACAAT
GGGGTCGTCTTTAG
>AY032626.1_CHINA-CH-1a
ATGTTGGGGAAATGCTTGACCACGGGCTGTTGCTCGCGATTGCTTTCTTTGTGGTGTAT
CGTGCCGTTCTGTTTTGCTGTGCTCGTCAACGCCAACAGCAACAGCAGCTCTCATTTTC
AGTTGATTTATAACTTGACGCTATGTGAGCTGAATGGCACAGATTGGCTGGCTAACAA
ATTTGACTGGGCAGTGGAGACTTTTGTCATCTTTCCCGTGTTGACTCACATTGTTTCCTA
TGGGGCACTCACCACCAGCCATTTCCTTGACACAGTTGGTCTGGTCACTGTGTCCACCG
CCGGGTTTTATCACGGGCGGTATGTCTTGAGTAGCATCTACGCGGTCTGTGCTCTGGCT
GCGTTGATTTGCTTCGTCATTAGGCTTGCGAAGAACTGCATGTCCTGGCGCTACTCTTG
TACCAGATATACCAACTTCCTTCTGGACACTAAGGGCAGACTCTATCGTTGGCGGTCGC
CCGTTATTGTAGAGAAAGGGGGTAAGGTTGAGGTCGAGGGTCACCTGATCGACCTCAA

AAGAGTTGTGCTTGATGGTTCCGTGGCAACCCCTTTAACCAGAGTTTCAGCGGAACAAT
GGGGTCGTCTCTAG
>AF494042.1_EE.UU-P129
ATGTTGGGGAAATGCTTGACCGCGGGCTGTTGCTCGCGATTGCTTTCTTTGTGGTGTAT
CGTGCCGTTCTGTTTTGCTGTGCTCGGCAGCGCCAACAGCAGCAGCAGCTCTCATTTTC
AGTTGATTTATAACTTGACGCTATGTGAGCTGAATGGCACAGATTGGCTGGCAGAAAA
ATTTGATTGGGCAGTGGAGACTTTTGTCATCTTTCCCGTGTTGACTCACATTGTTTCCTA
TGGTGCACTCACCACCAGCCATTTCCTTGACACAGTTGGTCTGGTTACTGTGTCCACCG
CCGGGTTTTATCACGGGCGGTATGTCTTGAGTAGCATCTACGCGGTCTGTGCTCTGGCT
GCGTTGATTTGCTTCGTTATTAGGCTTGCGAAGAACTGCATGTCCTGGCGCTACTCTTGT
ACCAGATATACCAACTTCCTTCTGGACACTAAGGGCAGACTCTATCGTTGGCGGTCGCC
CGTTATCATAGAAAAAGGGGGTAAGGTTGAGGTCGAAGGTCACCTGATCGACCTCAAA
AGAGTTGTGCTTGATGGTTCCGTGGCAACCCCTTTAACCAGAGTTTCAGCGGAACAATG
GGGTCGTCTCTAG
>AF331831.1_CHINA-BJ-4
ATGTTGGAGAAATGCTTGACCGCGGGCTGTTGCTCGCAATTGCTTTCTTTGTGGTGTAT
CGTGCCGTTCTGTTTTGCTGCGCTCGCCAACGCCAGCAACGACAGCAGCTCCCATCTAC
AGCTGATTTACAACTTGACGCTATGTGAGCTGAATGGCACAGATTGGCTAGCTAACAA
ATTTGATTGGGCAGTGGAGAGTTTTGTCATCTTTCCCGTTTTGACTCACATTGTCTCCTA
TGGTGCCCTCACTACCAGCCATTTCCTTGACACGGTCGCTTTAGTCACTGTGTCTACCGC
CGGGTTTGTTCACGGGCGGTATGTCCTATGTAGCATCTACGCGGTCTGTGCCCTGGCTG
CGTTGACTTGCTTCGTCATTAGGTTTGCAAAGAATTGCATGTCCTGGCGCTACGCGTGT
ACCAGATATACCAACTTTCTTCTGGACACTAAGGGCGGACTCTATCGTTGGCGGTCGCC
TGTCATCATAGAGAAAAGGGGCAAAGTTGAGGTCGAAGGTCATCTGATCGACCTCAAA
AGAGTTGTGCTTGATGGTTCCGTGGCAACCCCTATAACCAGAGTTTCAGCGGAACAAT
GGGGTCGTCCTTAG
>AF325691.1_EE.UU-NVSL_97-7985_IA_1-4-2
ATGTTGGAGAAATGCTTGACCGCGGGCTGTTGCTTGCGATTGCCTTCTTTGTGGTGTAT
CGTGCCGTTCTGTTTTGCTGTGCTCGTCAACGCCAACAACAGCAGCAGCTCCCATTTTC
AGTCGATTTATAACTTAACGCTATGTGAGCTGAATGGCACAGAATGGCTGAGTGAGAA
ATTTGATTGGGCAGTGGAGACTTTTGTCATCTTTCCCGTGTTAACTCACATTGTTTCCTA
TGGTGCACTCACCACCAGCCATTTCCTTGACACAGTTGGTCTGGTTACTGTGTCCACCG
CCGGGTTTCTCCACAGGCGGTATGTCTTGAGCAGCGTCTACGCGGTCTGTGCTCTGGCT
GCGTTGATTTGCTTCATCATTAGGCTTGCGAAGAACTGCATGTCCTGGCGCTACTCTTG

TACCAGATATACCAACTTTCTTCTGGACACTAAGGGCAAACTCTATCGTTGGCGGTCGC
CCGTTATCATAGAGAAAGGGGGCAGGGTTGAGGTCGAAGGTCACCTGATCGACCTCAA
AAGAGTTGTGCTTGATGGTTCCGCGGCAACCCCTTTAACCAGAGTTTCAGCGGAACAAT
GGGGTCGTCTCTAG
>AF184212.1_CHINA-SP
ATGTTGGGGAAATGCTTGACCGCGGGTTGCTGCTCGCGATTGCTTTCTTTTTGGTGTATC
GTGCCGTTCTGTTTTGCTGTGCTCGTCAACGCCAGCTACAGCAGCAGCTCTCATTTACA
GTTGATTTATAACTTGACGCTATGTGAGCTGAATGGTACAGATTGGCTGGCTAATAAAT
TTGATTGGGCAGTGGAGAGTTTTGTCATCTTTCCTGTGTTGACCCACATCGTTTCCTATG
GTGCACTAACCACCAGCCACTTCCTTGACACAGTTGGTCTGGTTACTGTGTCTACCGCC
GGGTTTTATCATGGGCGGTATGTCCTGAGTAGCATCTACGCGGTCTGTGCCCTGGCTGC
GTTAATTTGCTTCGTCATTAGGTTGGCGAAGAACTGTATGTCCTGGCGCTACTCATGCA
CCAGATACACCAACTTTCTTCTGGACACTAAGGGCAGACTCTATCGTTGGCGGTCGCCT
GTCATCATAGAGAAAGGGGGTAAGGTAGAGGTCGAAAGCCATCTGATCGACCTCAAA
AGAGTTGTGCTTGATGGGTCCGCGGCAACCCCTTTAACCAGAGTTTCAGCGGAACAAT
GGGGTCGTCCCTAG
>AF176348.2_CANADA-PA8
ATGTTGGGGAAATGCTTGACCGCGGGCTGGTGCTCGCAATTGCTTTCTTTGGGGTGTAT
CGTGCCGTTCTGTTTTGCTGTGCTCGCCAACGCCAGCAACGACAGCAGCTCCCATGTAC
AGCTGATTTACAACTTGACGCTATGTGAGCTGAATGGCACAGATTGGCTAGCTAACAA
ATTTGATTGGGCAGTGGAGAGTTTTGTCATCTTTCCCGTTTTGACTCACATTGTCTCCTA
TGGTGCCCTCACTACCAGCCATTTCCTTGACACAGTCGCTTTAGTCACTGTGTCTACCGC
CGGGTTTGTTCACGGGCGGTATGTCCTAAGTAGCATCTACGCGGTCTGTGCCCTGGCTG
CGTTGACTTGCTTCGTCATTAGGTTTGCAAAGAATTGCATGTCCTGGCGCTACGCGTGT
ACCAGATATACCAACTTTCTTCTGGACACTAAGGGCAGACTCTATCGTTGGCGGTCGCC
TGTCATCATAGAGAAAAGGGGCAAAGTTGAGGTCGAAGGTCATCTGATCGACCTCAAA
AGAGTTGTGCTTGATGGTTCCGTGGCAACCCCTATAACCAGAGTTTCAGCGGAACAAT
GGGGTCGCCCTTAG
>AB288356.1_EE.UU-EDRD-1
ATGTTGGGGAAATGCTTGACCGCGGGCTGTTGCTCGCGATTGCCTTTTTTGTGGTGTAT
CGTGCCGTTCTGTCTTGCTGCGCTCGTCAACGCCAGCGACAGCAGCAGCTCCCATTTAC
AGTTGATTTATAACCTGACGCTATGTGAGCTGAATGGCACAGATTGGCTGGCTGACAA
ATTTGATTGGGCAGTGGAGAGTTTTGTCATCTTTCCCGTGTTGACTCACATTGTTTCTTA
CTGCGCCCTCACTACCAGCCACTTCCTTGACACAGTTGGTCTGGTCGCTGTGTCTACCG

CCGGGTTTTACCACGGGCGGTATGTTCTGAGTAGCATCTATGCGGTCTGTGCCCTGGCT
GCGTTGGTTTGCTTCGTCATCAGATTGACGAAGAATTGCATGTCCTGGCGCTACTCATG
TACCAGATATACCAACTTTCTTCTGGATACCAAGGGCAGACTCTATCGTTGGCGGTCGC
CCGTCATCATAGAGAAAGGGGGTAAGGTTGAGGTTGAAGGTCATCTGATCGACCTCAA
GAGAGTTGTGCTTGATGGTTCCGCGGCAACCCCTATAACCAAAGTTTCAGCGGAACAA
TGGGGTCATCCCTAG
>MH791378.1_PERU-2015/15
ATGTTGGGGAAATGCTTGACCGCGGGCTGCTGCTCGCAATTGCTTTTTTTGTGGTGTAT
CGTGCCGTTCTGTTTTGTTGCGCTCGTCAACGCCAACAACAGCAGCAGCTCCCATTTAC
AGTTGATTTATAACCTGACGATATGTGAGCTGAATGGCACAGATTGGCTAAATAAAAG
TTTTGATTGGGCGGTGGAGACCTTCGTTATCTTTCCTGTGTTGACTCATATTGTCTCCTA
TGGCGCCCTCACCACCAGCCATTTCCTTGACACAGTCGGCCTGATCACCGTGTCTGCCG
CCGGATATTACCACAGGCGGTATGTCTTGAGTAGCATTTACGCCGTCTGCGCCCTGGCT
GCGTTAACTTGCTTCGTCATCAGGCTAACAAAAAATTGTATGTCCTGGCGTTACTCATG
CACCAGGTACACTAACTTTCTTCTGGACACCAAGGGCAAACTCTATCGTTGGCGGTCTC
CTGTCATCATAGAGAAAGGGGGTAAAATTGAGGTCGAAGGTCACCTGATCGACCTCAA
GAGAGTTGTGCTTGACGGTTCCGCGGCAACCCCTGTAACCAAAGTTTCAGCGGAACAA
TGGGGTCGTCCTTAG
>MH791376.1_PERU-2015/13
ATGTTGGGGAAATGCTTGACCGCGGGCTGCTGCTCGCAATTGCTTTTTTTGTGGTGTAT
CGTGCCGTTCTGTTTTGTTGCGCTCGTCAACGCCAACAACAGCAGCAGCTCCCATTTAC
AGTTGATTTATAACCTGACGATATGTGAGCTGAATGGCACAGATTGGCTAAATAAAAG
TTTTGATTGGGCGGTGGAGACCTTTGTTATCTTTCCTGTGTTGACTCATATTGTCTCCTA
TGGCGCCCTCACCACCAGCCATTTCCTTGACACAGTCGGCCTGATCACCGTGTCTGCCG
CCGGATATTACCACAGGCGGTATGTCTTGAGTAGCATTTACGCCGTCTGCGCCCTGGCT
GCGTTAACTTGCTTCGTCATCAGGCTAACAAAAAATTGTATGTCCTGGCGTTACTCATG
CACCAGGTACACTAACTATCTTCTGGACACCAAGGGCAAACTCTATCGTTGGCGGTCTC
CTGTCATCATAGAGAAAAGGGGTAAAATTGAGGTCGAAGGTCACCTGATCGACCTCAA
GAGAGTTGTGCTTGACGGTTCCGCGGCAACCCCTGTAACCAAAGTTTCAGCGGAACAA
TGGGGTCGTCCTTAG
>MH791379.1_PERU-2015/17
ATGTTGGGGAAATGCTTGACCGCGGGCTGCTGCTCGCAATTGCCTTTTTTGTGGTGTAT
CGTGCCGTTCTGTTTTGTTGCGCTCGTCAACGCCAACAACAGCAGCAGCTCTCATTTAC
AGTTGATTTATAACCTGACGATATGTGAGCTGAATGGCACAGATTGGCTAAATAAAAG

TTTTGATTGGGCGGTGGAGACCTTTGTTATCTTTCCTGTGTTGACTCATATTGTCTCCTA
TGGCGCCCTCACCACCAGCCATTTCCTTGACACAGTCGGCCTGATCACCGTGTCTGCCG
CCGGATACTACCACGGGCGGTATGTCTTGAGTAGCATTTACGCCGTCTGCGCCCTGGCT
GCGTTAACTTGCTTCGTCATCAGGCTAACAAAAAATTGTATGTCCTGGCGTTACTCATG
CACCAGATACACTAACTTTCTTCTGGACACCAAGGGCAAACTCTATCGTTGGCGGTCTC
CTGTCATCATAGAGAAAGGGGGTAAAATTGAGGTCGAAGGTCACCTGATCGACCTCAA
GAGAGTTGTGCTTGACGGTTCCGCGGCAACCCCTGTAACCAAAGTTTCAGCGGAACAA
TGGGGTCGTCCTTAG

>MH791383.1_PERU-2016/2

ATGTTGGGGAAATGCTTGACCGCGGGCTGCTGCTCGCAATTGCCTTTTTTGTGGTGTAT
CGTGCCGTTCTGTTTTGTTGCGCTCGTCAACGCCAACAACAGCAGCAGCTCCCATTTAC
AGTTGATTTATAACCTGACGATATGTGAGCTGAATGGCACAGATTGGCTAAATAAAAG
TTTTGATTGGGCGGTGGAGACCTTTGTTATCTTTCCTGTGTTGACTCATATTGTCTCCTA
TGGCGCCCTCACTACCAGCCATTTCCTTGACACAGTCGGCCTGATCACCGTGTCTGCCG
CCGGATACTACCACGGGCGGTATGTCTTGAGTAGCATTTACGCCGTCTGCGCCCTGGCT
GCGTTAACTTGCTTTGTCATCAGGCTAACAAAAAATTGTATGTCCTGGCGTTACTCATG
CACCAGATACACTAACTTTCTTCTGGACACCAAGGGCAAACTCTATCGTTGGCGGTCTC
CTGTCATCATAGAGAAAGGGGGTAAAATTGAGGTCGAAGGTCACCTGATCGACCTCAA
GAGAGTTGTGCTTGACGGTTCCGCGGCAACCCCTGTAACCAAAGTTTCAGCGGAACAA
TGGGGTCGTCCTTAG

>MH791385.1_PERU-2016/4

ATGTTGGGGAAATGCTTGACCGCGGGCTGCTGCTCGCAATTGCCTTTTTTGTGGTGTAT
CGTGCCGTTCTGTTTTGTTGCGCTCGTCAACGCCAACAACAGCAGCAGCTCCCATTTAC
AGTTGATTTATAACCTGACGATATGTGAGCTGAATGGCACAGATTGGCTAAATAAAAG
TTTTGATTGGGCGGTGGAGACCTTTGTTATTTTTCCTGTGTTGACTCATATTGTCTCCTA
TGGCGCCCTCACCACCAGCCATTTCCTTGACACAGTCGGCCTGATCACCGTGTCTGCCG
CCGGATATTACCACGGGCGGTATGTCTTGAGTAGCATTTACGCCGTCTGCGCCCTGGCT
GCGTTAACTTGCTTCGTCATCAGGCTAACAAAAAACTGTATGTCCTGGCGTTACTCATG
CACCAGATACACTAACTTTCTTCTGGACACCAAGGGCAAGCTCTATCGTTGGCGGTCTC
CTGTCATCATAGAGAAAGGGGGTAAAATTGAGGTCGAAGGTCACCTGATCGACCTCAA
GAGAGTTGTGCTTGACGGTTCCGCGGCAACCCCTGTAACCAAAGTTTCAGCGGAACAA
TGGGGTCGTCCTTAG

>MH791391.1_PERU-2017/7

ATGTTGGGGAAATGCTTGACCGCGGGCTGCTGCTCGCAATTGCTTTTTTTGTGGTGTAT
CGTGCCGTTCTGTTTTGTTGCGCTCGTCAACGCCAACAACAGCAGCAGCTCCCATTTAC
AGTTGATTTATAACCTGACGGTATGTGAGCTGAATGGCACAGATTGGCTAAATAGAAG
GTTTGATTGGGCGGTGGAGACCTTTGTTATCTTCCCTGTGTTGACTCATATTGTCTCCTA
TGGCGCCCTCACCACCAGCCATTTCCTTGACACAGTCGGCCTGATCACCGTGTCTGCCG
CTGGATATTACCACGGGCGGTATGTCTTGAGTAGCATTTACGCCGTCTGCGCCCTGGCT
GCGTTAACTTGCTTCGTCATCAGGCTAACAAAAAATTGTATGTCCTGGCGTTACTCATG
CACCAGGTACACCAACTATCTTCTGGACACCAAGGGCAAACTCTATCGTTGGCGGTCTC
CTGTCATCATAGAGAAAGGGGGTAAAATTGAGGTCGAAGGTCACCTGATCGACCTCAA
GAGAGTTGTGCTTGACGGTTCCGCGGCAACCCCTGTAACCAAAGTTTCAGCGGAACAA
TGGGGTCGTCCTTAG
>MH791386.1_PERU-2016/5
ATGTTGGGGAAATGCTTGACCGCGGGCTGCTGCTCGCAATTGCCTTTTTTGTGGTGTAT
CGTGCCGTTCTGTTTTGTTGTGCTCGTCAACGCCAACAACAGCAGCAGCTCCCATTTAC
AGTTGATTTATAACCTGACGATATGTGAGCTGAATGGCACAGATTGGCTAAATAAAAG
TTTTGATTGGGCGGTGGAGACCTTTGTCATCTTTCCTGTGTTGACTCACATTGTCTCCTA
TGGCGCCCTCACCACCAGCCATTTCCTTGACACAGTCGGCCTGATCACCGTGTCTGCCG
CCGGATATTACCACGGGCGGTATGTCTTGAGTAGCATTTACGCCGTCTGCGCCCTGGCT
GCGTTAACTTGCTTCGTTATCAGGCTAACAAAAAATTGTATGTCCTGGCGTTACTCATG
CACCAGATACACTAACTTTCTTCTGGACACCAAGGGCAAACTCTATCGTTGGCGGTCTC
CTGTCATCATAGAGAAAGGGGGCAAAATTGAGGTCGAAGGTCACCTGATCGACCTCAA
GAGAGTTGTGCTTGACGGTTCCGCGGCAACCCCTGTAACCAAAGTTTCAGCGGAACAA
TGGGGTCGTCCTTAG
>MH791388.1_PERU-2016/16
ATGTTGGGGAAATGCTTGACCGCGGGCTGCTGCTCGCAATTGCTTTTTTTGTGGTGTAT
CGTGCCGTTCTGTTTTGTTGCGCTCGTCAACGCCAACAACAGCAGCAGCTCCCATTTAC
AGTTGATTTATAACCTGACGATATGTGAGCTGAATGGCACAGATTGGCTAAATAAAAG
TTTTGATTGGGCGGTGGAGACCTTTGTTATCTTTCCTGTGTTGACTCATATTGTCTCCTA
TGGCGCCCTCACCACCAGCCATTTCCTTGACACAGTCGGCCTGATCACCGTGTCTGCCG
CCGGATATTACCACGGGCGGTATGTCTTGAGTAGCATTTACGCCGTCTGCGCCCTAGCT
GCGTTAACTTGCTTCGTCATCAGGCTAACGAAAAATTGTATGTCCTGGCGTTACTCATG
CACCAGATACACTAACTTTCTTCTGGACACCAAGGGCAAACTCTATCGTTGGCGGTCTC
CTGTCATCATAGAGAAAGGGGGTAAAATTGAGGTCGAAGGTCACCTGATCGACCTCAA

GAGAGTTGTGCTTGACGGTTCCGCGGCAACCCCTGTAACCAAAGTTTCAGCGGAACAA
TGGGGTCGTCCTTAG
>MH791390.1_PERU-2017/6
ATGTTGGGGAAATGCTTGACCGCGGGCTGCTGCTCGCAATTGCCTTTTTTGTGGTGTAT
CGTGCCGTTCTGTTTTGTTGCGCTCGTCAACGCCAACAACAGCAGCAGCTCCCATTTAC
AGTTGATTTATAACCTGACGATATGTGAGCTGAATGGCACAGATTGGCTAAATAAAAG
TTTTGATTGGGCGGTGGAGACCTTTGTTATCTTTCCTGTGTTGACTCATATTGTCTCCTA
TGGCGCCCTCACCACCAGCCATTTCCTTGACACAGTCGGCCTGATCACCGTGTCTGCCG
CCGGATATTACCACGGGCGGTATGTCTTGAGTAGCATTTACGCCGTCTGCGCCCTGGCT
GCGTTAATTTGCTTCGTCATCAGGCTAACAAAAAATTGTATGTCCTGGCGTTACTCATG
CACCAGGTACACCAACTTTCTTCTGGACACCAAGGGCAAACTCTATCGTTGGCGGTCTC
CTGTCATCATAGAGAAAGGGGGTAAAATTGAGGTCGAAGGTCACCTGATCGACCTCAA
GAGAGTTGTGCTTGACGGTTCCGCGGCAACCCCTGTAACCAAAGTTTCAGCGGAACAA
TGGGGTCGTCCTTAG
>48_Montana2019
GTGTTGGGGAAATGCTTGACCGCGGGCTGCTGCTCGCAATTGCCTTTTTTGTGGTGTAT
CGTGCCGTTCTGTCTTGTTGTGCTCGTCAACGCCAACAACAGCAGCAGCTCCCATTTAC
AGTTGATTTATAACCTGACGATATGTGAGCTGAATGGCACAGATTGGCTAAACAAAAG
TTTTGATTGGGCGGTGGAGACCTTTGTCATCTTTCCTGTGTTGACTCATATTGTCTCCTA
TGGCGCCCTTACCACCAGTCATTTCCTTGACACAGTCGGCCTGATCACCGTGTCTGCCG
CCGGATATTACCACGGGCGGTATGTCTTGAGTAGCATTTACGCCGTCTGCGCCTTAGCC
GCGTTAATTTGCTTCATCATCAAGCTAACAAAAAATTGTATGTCCTGGCGTTACTCATG
CACCAGGTACACTAATTTTCTTCTGGACACCAAGGGCAAACTCTATCGTTGGCGGTCTC
CTGTCATCATAGAGAAAGGGGGTAAAGTTGAGGTCCAAGGTCACCTGATAGACCTCAA
AAGAGTTGTGCTTGACGGTTCCGCGGCTACCCCTGTAACCAAAGTTTCAGCGGAACAA
TGGGGTCGTCCTTAG
>47_Montana2019
GTGTTGGGGAAATGCTTGACCGCGGGCTGCTGCTCGCAATTGCCTTTTTTGTGGTGTAT
CGTGCCGTTCTGTTTTGTTGTGCTCGTCAACGCCAACAACAGCAGCAGCTCCCATTTGC
AGTTGATTTATAACCTGACGATATGTGAGCTGAATGGCACAGATTGGCTAAACGAACA
TTTTGATTGGGCGGTGGAGACCTTTGTCATCTTTCCTGTGTTGACTCACATTGTCTCCTA
TGGTGCCCTCACCACCAGCCATTTCCTTGACACAGTCGGCCTGATCACCGTGTCTACCG
CCGGATATTACCACGGGCGGTATGTCTTGAGTAGCATTTACGCCGTCTGCGCCCTAGCT
GCGTTAACTTGCTTCATCATCAGGCTAACGAAAAACTGTATGTCCTGGCGTTACTCATG

CACCAGGTACACTAATTTTCTTCTGGACACCAAGGGCAAACTCTATCGTTGGCGGTCTC
CTGTCATCATAGAGAAAGGGGGTAAAGTTGAGGTCGAAGGTCACCTTATCGACCTCAA
GAGAGTTGTACTTGACGGTTCCGCGGCTACCCCTGTAACCAAAGTTTCAGCGGAACAA
TGGGGTCGTCCTTAG
>46_Montana2019
GTGTTGGGGAAATGCTTGACCGCGGGCTGCTGCTCGCAATTGCCTTTTTTGTGGTGTAT
CGTGCCGTTCTGTTTTGTTGTGCTCGTCAACGCCAACAACAGCAGCAGCTCCCATTTAC
AGTTGATTTATAACCTGACGATATGTGAGCTGAATGGCACAGATTGGCTAAACGAACA
TTTTGATTGGGCGGTGGAGACCTTTGTCATCTTTCCTGTGTTGACTCACATTGTCTCCTA
TGGTGCCCTCACCACCAGCCATTTCCTTGACACAGTCGGCCTGATCACCGTGTCTACCG
CCGGATATTACCACGGGCGGTATGTCTTGAGTAGCATTTACGCCGTCTGCGCCCTAGCT
GCGTTAACTTGCTTCATCATCAGGCTAACGAAAAACTGTATGTCCTGGCGTTACTCATG
CACCAGGTACACTAATTTTCTTCTGGACACCAAGGGCAAACTCTATCGTTGGCGGTCTC
CTGTCATCATAGAGAAAGGGGGTAAAGTTGAGGTCGAAGGTCACCTGATCGACCTCAA
GAGAGTTGTACTTGACGGTTCCGCGGCTACCCCTGTAACCAAAGTTTCAGCGGAACAA
TGGGGTCGTCCTTAG
>45_Montana2019
GTGTTGGGGAAATGCTTGACCGCGGGCTGCTGCTCGCAATTGCCTTTTTTGTGGTGTAT
CGTGCCGTTCTGTTTTGTTGTGCTCGTCAACGCCAACAACAGCAGCAGCTCCCATTTAC
AGTATGTTTATAACCTGACGATATGTGAGCTGAATGGCACAGATTGGCTAAACGTTCAT
TTTGATTGGGCGGTGGAGACCTTTGTCATCTTTCCTGTGTTGACTCACATTGTCTCCTTT
GGTGCCCTTACCACCAGCCATTTCCTTGACACATTCGGCCTGATCACCGTGTCTACCGC
CGGATATTACCACGGGCGGTATGTCTTGAGTAGCATTTACGCCGTCTGCGCCCTAGCCG
CGTTAATTTGCTTCATCATCAAGCTAACAAAAAATTGTATGTCCTGGCGTTACTCATGC
ACCAGGTACACTAATTTTCATCTGGACACCAAGGGCAAACTCTATCGTTGGCGGTCTCC
TGTCATCATAGAGAAAGGGGGTAAAGTTGAGGTCCAAGGTCACCTGATAGACCTCAAA
AGAGTTGTGCTTGACGGTTCCGCGGCTACCCCTGTAACCAAAGTTTCAGCGGAACAAT
GGGGTCGTCCTTAG
>44_Montana2019
ATGTTGGGGAAATGCTTGACCGCGGGCTGCTGCTCGCAATTGCCTTTTTTGTGGTGTAT
CGTGCCGTTCTGTTTTGTTGCGCTCGTCAACGCCGGCAACAGCATCAGCTCCCATTTAC
AGGTGATTTATAACCTGACGATATGTGAGCTGAATGGCACAGATTGGCTAAATAAAAG
CTTTGATTGGGCGGTGGAGACCTTTGTTATCTTTCCTGTGGTGACTCATATTGTCTCCTA
TGGCGCCTTCACCACCAGCCATTTCCTTGACACAGTCGGCCTGATCACCGTGTCTGCCG

CCGGATATTACCCCGGACGGTATGTTTTGAGTAGCATTTACGCCGTTTGCGCCCTGGAT
GCGTTAACTTGCTTCGTCATCAGGCTAACAAAAAATTGTATGTCCTGGCGTTATTCATG
CACCAGGTACAGTAATTTTCTTCTGGACACCAAGGGCAAACTCTATCGTTGGCGGTCTC
CTGTCATCATAGAGAAAGGGGGTAAAATAGAGGTAGAAGGTCACATGATCGACCTCAA
GAGAGTTGTACTTGACGGTTCCGCGGCTACCCCTGTAACCAAAGTTTCAGCGGAACAA
TGGGGTCGTCCTTAG
>43_Montana2019
ATGTTGGGGAAATGCTTGACCGCGGGCTGCTGCTCGCAATTGCCTTTTTTGTGGTGTAT
CGTGCCGTTCTGTTTTGTTGCGCTCGTCAACGCCGGCAACAGCATCAGCTCCCATTTAC
AGGTGATTTATAACCTGACGATATGTGAGCTGAATGGCACAGATTGGCTAAATAAAAG
CTTTGATTGGGCGGTGGAGACCTTTGTTATCTTTCCTGTGTTGACTCATATTGTCTCCTA
TGGCGCCCTCACCACCAGCCATTTCCTTGACACAGTCGGCCTGATCACCGTGTCTGCCG
CCGGATATTACCACGGACGGTATGTTTTGAGTAGCATTTACGCCGTCTGCGCCCTGGCT
GCATTAACTTGCTTCGTCATCAGGCTAACAAAAAATTGTATGTCCTGGCGTTATTCATG
CACCAGGTACACTAATTTTCTTCTGGACACCAAGGGCAAACTCTATCCTTGGCGGTCTC
CTGTCATCATAAAGAAAGGGGGGAAAATAGAGGTAGAAGGTCACATGATCGACCTCA
AGAGAGTTGTACTTGACGGTTCCGCGGCTACCCCTGTAACCAAAGTTTCAGCGGAACA
ATGGGGTCGTCCTTAG
>42_Montana2019
ATGTTGGGGAAATGCTTGACCGCGGGCTGCTGCTCGCAATTGCCTTTTTTGTGGTGTAT
CGTGCCATTCTGTTTTGTTGCGCTCGTCAACACCCGCAACACCAGCATCTCCCTTTTACT
TTTGATTTATAACCTGACGATATGTGAGCTGAATGGCACAGATTGGCTAAATAAAAGCT
TTGATTGGGCGGTGGAGACCTTTGTTATCTTTCCTGTGTTGACTCATATTGTCTCCTATG
GCGCCCTCACCACCAGCCATTTCCTTGACACAGTCGGCCTGATCACCGTGTCTGCCGCC
GGATATTACCACGGACGGTATGTTTTAAGTAGCATTTACGCCGTCTGCGCCCTGGCTGC
GTTAACTTGCTTCGTCATCTCGCTATCTAAAAATTGTATGTCCTGGCGTTATTCTTGCAC
CAAGTACACTACTTTTCTTCTGGACACCAAGTGCAATCTCTCTCGCGGAAGTTCTCCTG
TGATGATACACCGCTGGGGTTAAAACGAAGTCTCATGTTTCCTGTCCCTCTTCAAAAGA
GTTGTGCTTGACGGTTCCGCGGCTACCCCTGTAACCAAAGTTTCAGCGGAACAATGGG
GTCGTCCTTAG
>41_Montana2019
ATGTTGGGGAAATGCTTGACCGCGGGCTGCTGCTCGCAATTGCCTTTTTTGTGGTGTAT
CGTGCCGTTCTGTTTTGTTGCGCTCGTCAACGCCAGCAACAACAGCAGCTCCCATTTAC
TGTTGATTTATAACCTGACGATATGTGAGCTGAACGGCACAGATTGGCTAAATAAAAG

TTTTGATTGGGCGGTGGAGACCTTTGTTATCTTTCCTGTGTTGACTCATATTGTCTCCTA
TGGCGCCCTCACCACCAGCCATTTCCTTGACACAGTCGGCCTGATCACCGTGTCTGCCG
CCGGATATTACCACGGACGGTATGTTTTGAGTAGCATTTACGCCGTCTGCGCCCTGGCT
GCGTTAACTTGCTTCATCATCAGGCTAACAAAAAATTGTATGTCCTGGCGTTACTCATG
CACCAGGTACACTAATTTTCTTCTGGACACCAAGGGCAAACTCTATCGTTGGCGGTCTC
CTGTCATCATAGAGAAAGGGGGTAAAGTTGACGTCGAAGGTCACCTGATCGACCTCAA
GAGAGTTGTACTTGACGGTTCCGCGGCTACCCCTGTAACCAAAGTTTCAGCGGAACAA
TGGGGTCGTCCTTAG
>40_Montana2019
ATGTTGGGGAAATGCTTGACCGCGGGCTGCTGCTCGCAATTGCCTTTTTTGTGGTGTAT
CGTGCCGTTCTGTTTTGTTGCGCTCGTCAACGCCAGCAACAACAGCAGCTCCCATTTAC
AGTTGATTTATAACCTGACGATATGTGAGCTGAACGGCACAGATTGGCTAAATAAAAG
TTTTGATTGGGCGGTGGAGACCTTTGTTATCTTTCCTGTGTTGACTCATATTGTCTCCTA
TGGCGCCCTCACCACCAGCCATTTCCTTGACACAGTCGGCCTGATCACCGTGTCTGCCG
CCGGATATTACCACGGACGGTATGTTTTGAGTAGCATTTACGCCGTCTGCGCCCTGGCT
GCGTTAACTTGCTTCATCATCAGGCTAACAAAAAATTGTATGTCCTGGCGTTACTCATG
CACCAGGTACACTAATTTTCTTCTGGACACCAAGGGCAAACTCTATCGTTGGCGGTCTC
CTGTCATCATAGAGAAAGGGGGTAAAGTTGAGGTCGAAGGTCACCTGATCGACCTCAA
GAGAGTTGTACTTGACGGTTCCGCGGCTACCCCTGTAACCAAAGTTTCAGCGGAACAA
TGGGGTCGTCCTTAG
>39_Montana2019
ATGTTGGGGAAATGCTTGACCGCGGGCTGCTGCGCGCGGGGGCTTTTTTGGTGGTGGAT
CGATAAGTTCTGTTTGGTTTCGCTCGTCAATGCCGAGAACAGCAGCAGCTCCCATTTAA
GATAAATCCATATTCGGGCGGTATGCGGTCTGAATGGCACAAATTGGCTAAATAGAAG
GTTTGATTGGGCGGTGTAGACCTTTGTTGTCTTTCCTGTGTTGACTCATATTGTCTCCTA
TGGCGCCCTCACCACCAGCCATTTCCTTGACACAGTCGGTTTGATTAAGGTGTCTGCCG
CCGGATATTACCACGGGCGGCATGTCAAAAGTAGCATTTACGCCGTGTGCGCCCTGGC
TGCGATGGGCCGATTTGTCATCAGACTAACAAAAAATTGCACCTCCGGGCGGGACTCA
CGCACCAGATACGCTAACTATCTTCTGGACACCAAGGGCAAACTATATCGTTGGCGGT
CTCCTGTCATCATAGAGAAAGGGGGTAAAATTGAGGTCGAAGGTCACCTGATCGACCT
CAAGAGAGTTGTGCTTGACGGTTCCGCGGCGACTCCTGTACCCTAAGTTTCAGCGGAAC
AATGGGGTCGTCCTTAG
>38_Montana2019

ATGTTGGGGAAATGCTTGACCGCGGGCTGCTGCGCGCGGGGGCTTTTTTGGTGGTGGAT
CGATAAGTTCTGTTTGGTTTCGCTCGTCAATGCCGAGAACAGCAGCAGCTCCCATTTAA
GATAAATCCATATTCGGGCGGTATGCGGTCTGAATGGCACAAATTGGCTAAATAGAAG
GTTTGATTGGGCGGTGTAGACCTTTGTTGTCTTTCCTGTGTTGACTCATATTGTCTCCTA
TGGCGCCCTCACCACCAGCCATTTCCTTGACACAGTCGGTTTGATGAAGGTGTCTGCCG
CCGGATATTACCACGGGCGGCATGTCAAAAGTAGCATTTACGCCGTGTGCGCCCTGGC
TGCGATGGGCCGATTTGTCATCAGACTAACAAAAAATTGTACCTCCGGGCGGGACTCA
CGCACCAGATACGCTAACTATCTTCTGGACACTAAGGGCAAACTATATCGTTGGCGGTC
TCCTGTCATCATAGAGAAAGGGGGTAAAATTGAGGTCGAAGGTCACCTGATCGACCTC
AAGAGAGTTGTGCTTGACGGTTCCGCGGCGACTCCTGTAACCAAAGTTTCAGCGGAAC
AATGGGGTCGTCCTTAG
>37_Montana2019
ATGTTGGGGAAATGCTTGACCGCGGGCTGCTGCGCGCGGGGGCTTTTTTGGTGGTGGAT
CGATAAGTTCTGTTTGGTTTCGCTCGTCAATGCCGAGAACAGCAGCAGCTCCCATTTAA
GATAAATCCATATTCGGGCGGTATGCGGTCTGAATGGCACAAATTGGCTAAATAGAAG
GTTTGATTGGGCGGTGTAGACCTTTGTTGTCTTTCCTGTGTTGACTCATATTGTCTCCTA
TGGCGCCCTCACCACCAGCCATTTCCTTGACACAGTCGGTTTGATTAAGGTGTCTGCCG
CCGGATATTACCACGGACGGCATGTCAAAAGTAGCATTTACGCCGTGTGCGCCCTGGC
TGCGATGGGCCGATTTGTCATCAGACAACGAAAAAATTGCACCTCCGGGCGGGACTCA
CGCACCAGATACGCTAACTATCTTCTGGACACCAAGGGCAAACTATATCGTTGGCGGT
CTCCTGTCATCATAGAGAAAGGGGGCAAAATTGAGGTTGTCTTTCACCTGATCGACCTC
AAGAGAGTTGTGCTTGACGGTTCCGCGGCGACTCCTGTACCCTAAGTTTCAGCGGAAC
AATGGGGTCGTCCTTAG
>36_Montana2019
ATGTTGGGGAAATGCTTGACCGCGGGCTGCTGCGCGCGGGGGCTTTTTTGGTGGTGGAT
CGATAAGTTCTGTTTGGTTTCGCTCGTCAATGCCGAGAACAGCAGCAGCTCCCATTTAA
GATAAATCCATATTCGGGCGGTATGCGGTCTGAATGGCACAAATTGGCTAAATAGAAG
GTTTGATTGGGCGGTGTAGACCTTTGTTGTCTTTCCTGTGTTGACTCATATTGTCTCCTA
TGGCGCCCTCACCACCAGCCATTTCCTTGACACAGTCGGTTTGATGAAGGTGTCTGCCG
CCGGATATTACCACGGGCAGCGTGTCAAAAGTAGCATTTACGCCGTGTGCGCCCTGGC
TGCGATGGGCCGATTTGTCATCAGACAACGAAAAAATTGTACCTCCGGGCGGGACTCA
CGCACCAGATACGCTAACTATCTTCTGGACACTAAGGGCAAACTATATCGTTGGCGGTC
TCCTGTCATCATAGAGAAAGGGGGTAAAATTGAGGTTGGAGGTCACCTGATCGACCTC

AAGAGAGTTGTGCTTGACGGTTCCGCGGCGACTCCTGTAACCAAAGTTTCAGCGGAAC
AATGGGGTCGTCCTTAG
>35_Montana2019
ATGTTGGGGAAATGCTTGACCGCGGGCTGCTGCTCGCAATTGCTTTTTTTGTGGTGTAT
CGTGCCGTTCTGTTTTGTTGCGCTCGTCAACGCCAGCAACAGCAGCAGCTCCCATTTTC
AGTTGATTTATAACCTGACGATATGCGAGCTGAATGGCACAGATTGGCTAAATAGAAG
TTTTGATTGGGCGGTAGAGACCTTTGTTATCTTTCCTGTGTTGACTCATATTGTCTCCTA
TGGCGCCCTCACCACCAGCCATTTCCTTGACACAGTCGGCCTGATCACCGTGTCTGCCG
CCGGATATTACCACGGGCGGTATGTCTTGAGTAGCATTTATGCCGTCTGCGCCCTGGCT
GCGTTAACTTGCTTTGTCATCAGGCTAACAAAAAATTGCATGTCCTGGCGTTACTCGTG
CACCAGGTACACTAACTATCTTCTGGACACTAAGGGCAAACTCTATCGTTGGCGGTCTC
CTGTCATCATAGAGAAAGGGGGTAAAATTGAGGTCGAAGGTCACCTGATCGACCTCAA
GAGAGTTGTGCTTGACGGTTCCGCGGCAACCCCTGTAACCAAAGTTTCAGCGGAACAA
TGGGGTCGTCCTTAG
>34_Montana2019
ATGTTGGGGAAATGCTTGACCGCGGGCTGCTGCTCGCAATTGCTTTTTTTGTGGTGTAT
CGTGCCGTTCTGTTTTGTTGCGCTCGTCAACGCCAGCAACAGCAGCAGCTCCCATTTTC
AGTTGATTTATAACCTGACGATATGCGAGCTGAATGGCACAGATTGGCTAAATAGAAG
TTTTGATTGGGCGGTAGAGACCTTTGTTATCTTTCCTGTGTTGACTCATATTGTCTCCTA
TGGCGCCCTCACCACCAGCCATTTCCTTGACACAGTCGGCCTGATCACCGTGTCTGCCG
CCGGATATTACCACGGGCGGTATGTCTTGAGTAGCATTTATGCCGTCTGCGCCCTGGCT
GCGTTAACTTGCTTTGTCATCAGGCTAACAAAAAATTGCATGTCCTGGCGTTACTCGTG
CACCAGGTACACTAACTATCTTCTGGACACTAAGGGCAAACTCTATCGTTGGCGGTCTC
CTGTCATCATAGAGAAAGGGGGTAAAATTGAGGTCGAAGGTCACCTGATCGACCTCAA
GAGAGTTGTGCTTGACGGTTCCGCGGCAACCCCTGTAACCAAAGTTTCAGCGGAACAA
TGGGGTCGTCCTTAG
>33_Montana2019
ATGTTGGGGAAATGCTTGACCGCGGGCTGCTGCTCGCAATTGCCTTTTTTGTGGTGTAT
CGTGCCGTTCTGTTTTGTTGTGCTCGTCAACGCCAACAACAGCAACAGCTCCCATTTAC
AGTTGATTTATAACCTGACGATATGTGAGCTGAATGGCACAGATTGGCTAAATAAAAG
TTTTGATTGGGCGGTGGAGACCTTTGTTATCTTTCCTGTGTTGACTCATATTGTCTCCTA
CGGCGCCCTCACCACCAGCCATTTCCTTGACACAGTCGGCCTGATCACCGTGTCTGCCG
CCGGATACTACCACGGACGGTATGTCTTAAGTAGCATTTACGCCGTCTGCGCCATGGCT
GCGTTAACTTGCTTCGTCATCAGGCTAACAAAAAATTGTATGTCCTGGCGTTACTCATG

TACCAGGTACACTAATTTTCTTCTGGACACCAAGGGCAAACTCTATCGTTGGCGGTCTC
CTGTCATCATAGAGAAAGGGGGTAAAGTTGAGGTCGAAGGTCACCTGATCGACCTCAA
GAGAGTTGTGCTTGACGGTTCCGCGGCAACCCCTGTAACCAAAGTTTCAGCGGAACAA
TGGGGTCGTCCTTAG
>32_Montana2019
ATGTTGGGGAAATGCTTGACCGCGGGCTGCTGCTCGCAATTGCCTTTTTTGTGGTGTAT
CGTGCCGTTCTGTTTTGTTGTGCTCGTCAACGCCAACAACAGCAACAGCTCCCATTTAC
AGTTGATTTATAACCTGACGATATGTGAGCTGAATGGCACAGATTGGCTAAATAAAAG
TTTTGATTGGGCGGTGGAGACCTTTGTTATCTTTCCTGTGTTGACTCATATTGTCTCCTA
CGGCGCCCTCACCACCAGCCATTTCCTTGACACAGTCGGCCTGATCACCGTGTCTGCCG
CCGGATACTACCACGGACGGTATGTCTTAAGTAGCATTTACGCCGTCTGCGCCATGGCT
GCGTTAACTTGCTTCGTCATCAGGCTAACAAAAAATTGTATGTCCTGGCGTTACTCATG
CACCAGATACACTAATTTTCTTCTGGACACCAAGGGCAAACTCTATCGTTGGCGGTCTC
CTGTCATCATAGAGAAAGGGGGTAAAGTTGAGGTCGAAGGTCACCTGATCGACCTCAA
GAGAGTTGTGCTTGACGGTTCCGCGGCAACCCCTGTAACCAAAGTTTCAGCGGAACAA
TGGGGTCGTCCTTAG
>31_Montana2019
ATGTTGGGGAAATGCTTGACCGCGGGCTGCTGCTCGCAATTGCCTTTTTTGTGGTGTAT
CGTGCCGTTCTGTTTTGTTGTGCTCGTCAACGCCAACAACAGCAACAGCTCCCATTTAC
AGTTGATTTATAACCTGACGATATGTGAGCTGAATGGCACAGATTGGCTAAATAAAAG
TTTTGATTGGGCGGTGGAGACCTTTGTTATCTTTCCTGTGTTGACTCATATTGTCTCCTA
CGGCGCCCTCACCACCAGCCATTTCCTTGACACAGTCGGCCTGATCACCGTGTCTGCCG
CCGGATACTACCACGGACGGTATGTCTTAAGTAGCATTTACGCCGTCTGCGCCATGGCT
GCGTTAACTTGCTTCGTCATCAGGCTAACAAAAAATTGTATGTCCTGGCGTTACTCATG
TACCAGGTACACTAATTTTCTTCTGGACACCAAGGGCAAACTCTATCGTTGGCGGTCTC
CTGTCATCATAGAGAAAGGGGGTAAAGTTGAGGTCGAAGGTCACCTGATCGACCTCAA
GAGAGTTGTGCTTGACGGTTCCGCGGCAACCCCTGTAACCAAAGTTTCAGCGGAACAA
TGGGGTCGTCCTTAG
>30_Montana2019
ATGTTGGGGAAATGCTTGACCGCGGGCTGCTGCTCGCAATTGCCTTTTTTGTGGTGTAT
CGTGCCGTTCTGTTTTGTTGTGCTCGTCAACGCCAACAACAGCAACAGCTCCCATTTAC
AGTTGATTTATAACCTGACGATATGTGAGCTGAATGGCACAGATTGGCTAAATGAAAG
TTTTGATTGGGCGGTGGAGACCTTTGTTATCTTTCCTGTGTTGACTCATATTGTCTCCTA
CGGCGCCCTCACCACCAGCCATTTCCTTGACACAGTCGGCCTGATCACCGTGTCTGCCG

CCGGATACTACCACGGACGGTATGTCTTAAGTAGCATTTACGCCGTCTGCGCCATGGCT
GCGTTAACTTGCTTCGTCATCAGGCTAACAAAAAATTGTATGTCCTGGCGTTACTCATG
TACCAGGTACACTAATTTTCTTCTGGACACCAAGGGCAAACTCTATCGTTGGCGGTCTC
CTGTCATCATAGAGAAAGGGGGTAAAGTTGAGGTCGAAGGTCACCTGATCGACCTCAA
CAGAGTTGTGCTTGACGGTTCCGCGGCAACCCCTGTAACCAAAGTTTCAGCGGAACAA
TGGGGTCGTCCTTAG
>29_Montana2019
ATGTTGGGGAAATGCTTGACCGCGGGCTGCTGCTCGCAATTGCTTTTTTTGTGGTGTAT
CGTGCCGTTCTGTTTTGTTGCGCTCGTCAACGCCAGCAACAGCAGCAGCTCCCATTTTC
AGTTGATTTATAACCTGACGATATGCGAGCTGAATGGCACAGATTGGCTAAATAGAAA
TTTTGATTGGGCGGTAGAGACCTTTGTTATCTTTCCTGTGTTGACTCATATTGTCTCCTA
TGGCGCCCTCACCACCAGCCATTTCCTTGACACAGTCGGCCTGATCACCGTGTCTGCCG
CCGGATATTACCACGGGCGGTATGTCTTGAGTAGCATTTATGCCGTCTGCGCCCTGGCT
GCATTAATTTGCTTTGTCATCAGGCTAACAAAAAATTGCATGTCCTGGCGTTACTCGTG
CACCAGGTACACTAACTATCTTCTGGACACTAAGGGCAAACTCTATCGTTGGCGGTCTC
CTGTCATCATAGAGAAAGGGGGTAAAATTGAGGTCGAAGGTCACCTGATCGACCTCAA
GAGAGTTGTGCTTGACGGTTCCGCGGCAACCCCTGTAACCAAAGTTTCAGCGGAACAA
TGGGGTCGTCCTTAG
>28_Montana2019
ATGTTGGGGAAATGCTTGACCGCGGGCTGCTGCTCGCAATTGCTTTTTTTGTGGTGTAT
CGTGCCGTTCTGTTTTGTTGCGCTCGTCAACGCCAGCAACAGCAGCAGCTCCCATTTTC
AGTTGATTTATAACCTGACGATATGCGAGCTGAATGGCACAGATTGGCTAAATAGAAG
TTTTGATTGGGCGGTAGAGACCTTTGTTATCTTTCCTGTGTTGACTCATATTGTCTCCTA
TGGCGCCCTCACCACCAGCCATTTCCTTGACACAGTCGGCCTGATCACCGTGTCTGCCG
CCGGATATTACCACGGGCGGTATGTCTTGAGTAGCATTTATGCCGTCTGCGCCCTGGCT
GCGTTAACTTGCTTTGTCATCAGGCTAACAAAAAATTGCATGTCCTGGCGTTACTCGTG
CACCAGGTACACTAACTATCTTCTGGACACTAAGGGCAAACTCTATCGTTGGCGGTCTC
CTGTCATCATAGAGAAAGGGGGTAAAATTGAGGTCGAAGGTCACCTGATCGACCTCAA
GAGAGTTGTGCTTGACGGTTCCGCGGCAACCCCTGTAACCAAAGTTTCAGCGGAACAA
TGGGGTCGTCCTTAG
>27_Montana2019
ATGTTGGGGAAATGCTTGACCGCGGGCTGCTGCTCGCAATTGCTTTTTTTGTGGTGTAT
CGTGCCGTTCTGTTTTGTTGCGCTCGTCAACGCCAGCAACAGCAGCAGCTCCCATTTTC
AGTTGATTTATAACCTGACGATATGCGAGCTGAATGGCACAGATTGGCTAAATAGAAG

TTTTGATTGGGCGGTAGAGACCTTTGTTATCTTTCCTGTGTTGACTCATATTGTCTCCTA
TGGCGCCCTCACCACCAGCCATTTCCTTGACACAGTCGGCCTGATCACCGTGTCTGCCG
CCGGATATTACCACGGGCGGTATGTCTTGAGTAGCATTTATGCCGTCTGCGCCCTGGCT
GCGTTAACTTGCTTTGTCATCAGGCTAACAAAAAATTGCATGTCCTGGCGTTACTCGTG
CACCAGGTACACTAACTATCTTCTGGACACTAAGGGCAAACTCTATCGTTGGCGGTCTC
CTGTCATCATAGAGAAAGGGGGTAAAATTGAGGTCGAAGGTCACCTGATCGACCTCAA
GAGAGTTGTGCTTGACGGTTCCGCGGCAACCCCTGTAACCAAAGTTTCAGCGGAACAA
TGGGGTCGTCCTTAG
>26_Montana2019
ATGTTGGGGAAATGCTTGACCGCGGGCTGCTGCTCGCAATTGCTTTTTTTGTGGTGTAT
CGTGCCGTTCTGTTTTGTTGCGCTCGTCAACGCCAGCAACAGCAGCAGCTCCCATTTTC
AGTTGATTTATAACCTGACGATATGCGAGCTGAATGGCACAGATTGGCTAAATAGAAG
TTTTGATTGGGCGGTAGAGACCTTTGTTATCTTTCCTGTGTTGACTCATATTGTCTCCTA
TGGCGCCCTCACCACCAGCCATTTCCTTGACACAGTCGGCCTGATCACCGTGTCTGCCG
CCGGATATTACCACGGGCGGTATGTCTTGAGTAGCATTTATGCCGTCTGCGCCCTGGCT
GCGCTAACTTGCTTTGTCATCAGGCTAACAAAAAATTGCATGTCCTGGCGTTACTCGTG
CACCAGGTACACTAACTATCTTCTGGACACTAAGGGCAAACTCTATCGTTGGCGGTCTC
CTGTCATCATAGAGAAAGGGGGTAAAATTGAGGTCGAAGGTCACCTGATCGACCTCAA
GAGAGTTGTGCTTGACGGTTCCGCGGCAACCCCTGTAACCAAAGTTTCAGCGGAACAA
TGGGGTCGTCCTTAG
>25_montana2019
ATGTTGGGGAAATGCTTGACCGCGGGCTGCTGCTCGCAATTGCTTTTTTTGTGGTGTAT
CGTGCCGTTCTGTTTTGTTGCGCTCGTCAACGCCAGCAACAGCAGCAGCTCCCATTTTC
AGTTGATTTATAACCTGACGATATGCGAGCTGAATGGCACAGATTGGCTAAATAGAAG
TTTTGATTGGGCGGTAGAGACCTTTGTTATCTTTCCTGTGTTGACTCATATTGTCTCCTA
TGGCGCCCTCACCACCAGCCATTTCCTTGACACAGTCGGCCTGATCACCGTGTCTGCCG
CCGGATATTACCACGGGCGGTATGTCTTGAGTAGCATTTATGCCGTCTGCGCCCTGGCT
GCGTTAACTTGCTTTGTCATCAGGCTAACAAAAAATTGCATGTCCTGGCGTTACTCGTG
CACCAGGTACACTAACTATCTTCTGGACACTAAGGGCAAACTCTATCGTTGGCGGTCTC
CTGTCATCATAGAGAAAGGGGGTAAAATTGAGGTCGAAGGTCACCTGATCGACCTCAA
GAGAGTTGTGCTTGACGGTTCCGCGGCAACCCCTGTAACCAAAGTTTCAGCGGAACAA
TGGGGTCGTCCTTAG
