## Supplementary material for "Genetic Variability, N-Glycosylation Sites, and Recombination Events in the ORF5 (GP5) Gene of Lineage 1A (NADC34) *Betaarterivirus americense* Strains from Lima, Peru Molecular Variability of the GP5 Glycoprotein in *Betaarterivirus americense*": Archivos complementarios: Supplementary Material 2. ORF5 lineage 1A sequences used for RDP v4.101 recombination analysis..pdf

>MF326985.1_Linaje 1A(NADC34)

ATGTTGGGGAAATGCTTGACCGCGGGCTGCTGCTCGCAATTGCCTTTTTTGTGGTGTATCGTGCCGTTCTGT
TTTGTTGCGCTCGTCAACGCCAACAACAGCAGCAGCTCCCATTTACAGTTGATTTATAACCTGACGATATGT
GAGCTGAATGGCACAGATTGGCTAAATAAAAGTTTTGATTGGGCGGTGGAGACCTTTGTTATTTTTCCTGTG
TTGACTCATATTGTCTCCTATGGCGCCCTCACCACCAGCCATTTCCTTGACACAGTCGGCCTGATCACCGTGT
CTGCCGCCGGATATTACCACGGGCGGTATGTCTTGAGTAGCATTTACGCCGTCTGCGCCCTGGCTGCGTTAA
CTTGCTTCGTCATCAGGCTAACAAAAAATTGTATGTCCTGGCGTTACTCATGCACCAGATATACTAACTTTCT
TCTGGACACCAAGGGCAAACTCTATCGTTGGCGGTCTCCTGTCATCATAGAGAAAGGGGGTAAAATTGAGG
TCGAAGGTCACCTGATCGACCTCAAGAGAGTTGTGCTTGACGGTTCCGCGGCAACCCCTGTAACCAAAGTTT
CAGCGGAACAATGGGGTCGTCCTTAG

>EF536003.1_Linaje 5A(VR2332)

ATGTTGGAGAAATGCTTGACCGCGGGCTGTTACTCGCAATTGCTTTCTTTGTGGTGTATCGTGCCGTTCTGTT
TTGCTGTGCTCGTCAACGCCAGCAACGACAGCAGCTCCCATCTACAGCTGATTTACAACTTGACGCTATGTG
AGCTGAATGGCACAGATTGGCTAGCTAACAAATTTGATTGGGCAGTGGAGAGTTTTGTCATCTTTCCCGTTT
TGACTCACATTGTCTCCTATGGTGCCCTCACTACTAGCCATTTCCTTGACACAGTCGCTTTAGTCACTGTGTCT
ACCGCCGGGTTTGTTCACGGGCGGTATGTCCTAAGTAGCATCTACGCGGTCTGTGCCCTGGCTGCGTTGACT
TGCTTCGTCATTAGGTTTGCAAAGAATTGCATGTCCTGGCGCTACGCGTGTACCAGATATACCAACTTTCTTC
TGGACACTAAGGGCAGACTCTATCGTTGGCGGTCGCCTGTCATCATAGAGAAAAGGGGCAAAGTTGAGGT
CGAAGGTCATCTGATCGACCTCAAAAGAGTTGTGCTTGATGGTTCCGTGGCAACCCCTATAACCAGAGTTTC
AGCGGAACAATGGGGTCGTCCTTAG

>AF066183.4_Linaje 5A(RespPRRS_MLV)

ATGTTGGAGAAATGCTTGACCGCGGGCTGTTGCTCGCAATTGCTTTCTTTGTGGTGTATCGTGCCGTTCTGT
TTTGCTGTGCTCGCCAACGCCAGCAACGACAGCAGCTCCCATCTACAGCTGATTTACAACTTGACGCTATGT
GAGCTGAATGGCACAGATTGGCTAGCTAACAAATTTGATTGGGCAGTGGAGAGTTTTGTCATCTTTCCCGTT
TTGACTCACATTGTCTCCTATGGTGCCCTCACTACCAGCCATTTCCTTGACACAGTCGCTTTAGTCACTGTGTC
TACCGCCGGGTTTGTTCACGGGCGGTATGTCCTAAGTAGCATCTACGCGGTCTGTGCCCTGGCTGCGTTGAC
TTGCTTCGTCATTAGGTTTGCAAAGAATTGCATGTCCTGGCGCTACGCGTGTACCAGATATACCAACTTTCTT
CTGGACACTAAGGGCGGACTCTATCGTTGGCGGTCGCCTGTCATCATAGAGAAAAGGGGCAAAGTTGAGG
TCGAAGGTCATCTGATCGACCTCAAAAGAGTTGTGCTTGATGGTTCCGTGGCAACCCCTATAACCAGAGTTT
CAGCGGAACAATGGGGTCGTCCTTAG

>DQ988080.1_Linaje 8A(Ingelvac_ATP)

ATGTTGGGGAGATGCTTGACCGCGGGCTGTTGCTCGCGATTGCTTTCTTTGTGGTGTATCGTGCCATTTTGT
TTTGCTGCGCTCGTCAACGCCAACAGCAACAGCAGCTCTCATCTTCAGTTAATTTACAACTTGACGCTATGTG
AGCTGAATGGCACAGATTGGCTGAAAGACAAATTTGATTGGGCATTGGAGACTTTTGTCATCTTTCCCGTGT
TGACTCACATTGTCTCATATAGTGCACTCACCACTAGCCATTTCCTTGACACAGTCGGTCTGGTTACTGTGTC
TACTGCCGGGTTCTACCACGGGCGGTATGTTCTGAGTAGCATCTACGCGGTCTGCGCTCTGGCCGCATTGAC
TTGCTTCGTCATTAGGCTTGCGAAGAACTGCATGTCCTGGCGCTACTCTTGTACCAGATATACTAACTTCCTT
CTGGACACTAAGGGCAGACTCTATCGCTGGCGGTCGCCCGTTATCATAGAGAAAGGGGGTAAGGTTGAGG
TCGAAGGTCACCTGATCGACCTCAAAAGAGTTGTGCTTGATGGTTCCGTGGCAACCCCTTTAACCAGAGTTT
CAGCGGAACAATGGGGTCGTCTTTAG

>JN660150.1_Linaje 1B(NADC31)

ATGTTGGGGAAATGCTCGACCGCGGGCTGCTGCTCGCCATTGCTTTTTTTGTGGTGTATCGTGCCGTCCTGT
TTAGTTGCGCTCGTCAACGCCAGCAAAAACAACAGCTCCCATTTACAGTCGATTTATAACCTGACGATATGT
GAGCTGAATGGCACAGATTGGCTAAATAAAAAATTTGACTGGGCAGTGGAGACCTTTGTCATTTTTCCTGTA
TTGACTCATATCGTCTCCTATGGTGCCCTCACCACCAGCCATTTCCTTGACGCAGTCGGTTTGGTCATCGTGT
CCACCGCCGGATACTATCACGGGCGGTATGTCCTGAGCAGCATTTACGCTGTCTGCGCCTTGGCCGCGCTG
ATTTGCTTCGCCATCAGGTTAACGAAAAACTGCATGTCCTGGCGCTACTCATGTACTAGGTATACTAACTTTC
TTCTAGACACCAAGGGCAAACTCTATCGTTGGCGGTCTCCCGTCATCATAGAAAAAAAAGGGAAAATCGAG
GTCAACGGTCACTTGATCGACCTCAAGAGAGTTGTGCTTGATGGTTCCGCAGCAACCCCTGTAACCAAAGTT
TCAGCGGAACAATGGGGTCATCCTTAG

>MF279253.1_Linaje 8D(DMZC181)

ATGTTGGGGAATTGCTTGACAGCTGTTTGTTGCTCGCGATTGTTTTCTTTGTGGTATATCGTGCCGTTTTGTT
TTGCTGTGCTCGTCAACGCCAACARCAGCAACAGCTCTCATTTTCAGTTGATTTATAACTTGACGATATGCGA
GCTGAATGGCACAGACTGGCTGAGCCACAACTTTGATTGGGCCGTGGAGACTTTTGTCATTTTTCCCGTGCT
AACTCACATCGTTTCCTATGGTGCACTCACCACCAGCCACTTCCTCGACACGGTTGGTCTGGTTACAGTGTCC
ACCGCCGGGTATTACCACGGGCGGTATGTCTTGAGCAGCATCTATGCAATCTGTGCCTTAGCAGCATTGACT
TGCTTCTTCATTAGGCTTACGAAAAATTGCATGTCCTGGCGTTACTCTTGTACCAGATATACCAACTTCCTTCT
AGACACTAAGGGCAGGCTCTATCGTTGGCGGTCTCCCGTCATCATAGAGAAAGGGGGTAAGGTTGAGGTT
GAAGGTCACCTAATCGACCTCAAAAGAGTCGTGCTTGATGGTTCCGCGGCAACYCCTTTAACCAAAGTTTCA
GCGGAGCAATGGGGTCGTCCCTAG

>KF611905.1_Linaje 1C(HENAN-XINX)

ATGTTGGTGAAATGCTTGGCCGCGGGTTGTTGCTCGCAATTGCCTTTTTTGTGGTGTATCGTGCCGTTCTATT
TTGCTGCGCTCGTCAACGCCAACAGCAACAGCAGCTCCCATCTACAGTTGATTTATAACCTGACGATATGTG
AGCTGAATGGTACGGATTGGCTGGGATCAAATTTTGATTGGGCAGTGGAGACTTTCGTCATCTTTCCTGTAT
TGACTCATATTGTCTCTTACGGCGCCCTTACCACTAGCCATTTTCTTGACACGGTCGGCCTGATCACTGTGTC
CACCGCCGGATATTTTCACAAGCGGTATGTGTTGAGTAGCATCTACGCTGTCTGTGCCCTGGCTGCGTTGGT
TTGCTTCGCCATTAGGTTGGCAAAAAATTGCATGTCCTGGCGCTACTCATGCACCAGATATACCAATTTTCTT
CTGGACACTAAGGGCAAACTCTACCGCTGGCGGTCACCCGTCATCATAGAGAAGGAGGGTAAAGTTGATG
TGGGTGGTCACTTAATCGACCTCAAGAGAGTTGTGCTTGATGGTTCCGCGGCAACCCCTGTAACCAAGATTT
CAGCGGAACAATGGGGTCGTCCATAG

>JX512910.2_ Linaje 8E(EE.UU-SRV07)

ATGTTGGGGAAGTGCTTGACCGCGTGCTGTTGCTCGCGATTGCTTTTTTTGTGGTGTATCGTGCCGTTCTATC
TTGCTGTGCTCGTCAACGCCAGCAACAACAACAGCCCTCATATTCAGTTGATTTATAACTTAACGCTATGTGA
GCTGAATGGCACAGATTGGCTGGCACAAAAATTTGACTGGGCAGTGGAGACTTTTGTCATCTTCCCCGTGTT
GACTCACATTGTTTCCTATGGGGCACTCACCACCAGCCATTTCCTTGACACAGTTGGTCTGGCCACTGTGTCC
ACCGCCGGATATTATCACGGGCGGTATGTCTTGAGTAGCATTTACGCAGTCTGTGCTCTGGCTGCGCTGATT
TGCTTTGTCATTAGGCTTGCGAAGAACTGCATGTCCTGGCGCTACTCTTGTACCAGATATACCAACTTCCTTC
TGGACACTAAGGGCAGACTCTATCGTTGGCGGTCGCCCGTCATTGTGGAGAAAGGGGGTAAGGTTGAGGT
CGAAGGTCACCTCATCGACCTCAAAAGAGTTGTGCTTGATGGTTCCGCGGCAACCCCTTTAACCAGAGTTTC
AGCGGAACAATGGGGTCGTCTCTAG

>EU807840.1_Linaje 9D(CHINA-CH-1R)

ATGTTGGGGAAATACTTGACCACGGGCTGCTGCTCGCGATTGCTTTCTTTGTGGTGTATCGTGCCGTTCTGT
TTTGCTGTGCTCGTCAACGCCAACAGCAACAGCAGCTCTCAATTTCAGTTGATTTATAACTTGACGCTATGTG
AGCTGAATGGCACAGATTGGCTGGCTAACAAATTTGACTGGGCAGTGGAGACTTTTGTCATCTTTCCCGTGT
TGACTCACATTGTGTCCTATGGGGCACTCACCACCAGCCATTTCCTTGACACAGTTGGTCTGGTCACTGTGTC
CACCGCCGGGTTTTATCACGGGCGGTATGTCTTGAGTAGCATCTACGCGGTCTGTGCTCTGGCTGCGTTGAT
TTGCTTCGTCATTAGGCTTGCGAAGAACTGCATGTCCTGGCGCTACTCTTGTACCAGATATACCAACTTCCTT
CAGGACACTAAGGGCAGACTCTATCGTTGGCGGTCGCCCGTTATTGTAGAGAAAGGGGGTAAGGTTGAGG
TCGAGGGTCACCTGATCGACCTCAAAAGAGTTGTGCTTGATGGTTCCGTGGCAACCCCTTTAACCAGAGTTT
CAGCGGAACAATGGGGTCGTCTCTAG

>DQ473474.1_Linaje 5B(COREA_DEL_SUR-LMY)

ATGTTGGGGAAATGCTTGACCGCGGGCTGTTGCTCGCAATTGCTTTTTTTGTGGTGTATCGTGCCGTCTTGT
TTTGTTGCGATCGTCAGCGCCAACAACAGCAGCAGCTCAAATTTACAGCTGATTTACAACTTGACGCTATGT
GAGCTGAATGGCACAGATTGGCTAGCTAACAGATTTGACTGGGCGGTGGAGTGTTTTGTTATTTTTCCTGTA
TTGACTCACATTGTCTCTTATGGTGCCCTAACCACTAGCCACTTCCTTGACACAGTCGGTTTGGTCACTGTGT
CTACCGCCGGATTTGTTCACGGGCGGTATGTTTTGAGTAGCATTTACGCGGTCTGTGCCCTGGCTGCGTTGA
TTTGCTTCGTCATTAGGCTTGCGAAGAATTGCATGTCCTGGCGCTACTCATGTACCAGATATACCAACTTTCT
TCTAGATACCAAGGGCAGACTCTACCGTTGGCGGTCGCCTGTCATTATAGAGAAAAGGGGCAAAGTTGAG
GTCGAGGGTCAACTAATCGACCCCAAAAGAGTTGTGCTTGATGGTTCCCCGGCAACCCCTGTAACCAGAGT
TTCAGCGGAACAATGGGGTCATCCTTAG

>DQ306879.1_Linaje 1E(EE.UU-99-3584)

ATGTTGGGGAAATGCTTGACCGCGGGCTGTTGCTCGCAATTGCCTTTTTTGTGGTGTATCGTGCCGTTCTGT
TTTGCTGCGCTCGTCAACGCCGGCAGCAACAGCAGCTCCCATTTACAGTTGATTTATAACCTGACGATATGT
GAGCTGAATGGCACAGATTGGCTGAACGACAAATTTGATTGGGCGGTGGAGACTTTTGTCATCTTTCCTGT
GTTGACTCACATTGTTTCMTATGGCGCCCTCACCACCAGCCATTTCCTTGACACAGTCGGTCTAGTTACTGTG
TCTACCGCCGGATATTACCATAGGCGGTATGTATTGAGTAGCATTTACGCTGTCTGTGCCCTGGCTGCGTTG
ATTTGCTTCGTCATCAGGTTGACGAAGAATTGCATGTCCTGGCGCTACTCATGCACCAGATATACTAACTTTC
TTTTGGACACCAAGGGCAGACTCTATCGTTGGCGGTCACCCGTCATCATAGAGAAAGGGGGCAAAGTTGA
GGTTGAAGGTCACCTGATCGACCTCAAGAGAGTTGTGCTTGACGGTTCCGCGGCAACCCCCGTAACCAAAG
TTTCAGCAGAACGATGGGGTCGTCCCTAG

>DQ176019.1_Linaje 1F(EE.UU-MN184A)

ATGTTGGGGAAATGCTTGACCGCGGGCTATTGCTCGCAATTGCTTTTTTTGTGGTGTATCGTGCCGTTCTGT
CTTGCTGCGCTCGTCAACGCCGACAGCAACAGCAGCTCCCATTTACAGTTGATTTATAAMTTAACGATATGT
GAGCTGAATGGCACAGACTGGCTGAACAATCATTTTAGTTGGGCAGTGGAGACTTTCGTTATCTTTCCTGTG
TTGACTCATATTGTTTCCTACGGCGCCCTCACTACCAGCCACCTCCTTGACACGGTCGGCCTGATCACTGTGT
CCACCGCCGGATACTGCCATAAGCGGTATGTCTTGAGTAGCATCTATGCTGTCTGCGCCCTGGCTGCGCTGA
TTTGCTTCGTCATCAGGTTGACGAAAAATTGTATGTCCTGGCGCTACTCATGTACCAGATATACCAACTTTCT
TCTGGACACCAAGGGCAGACTCTATCGCTGGCGGTCACCCGTCATCATAGAGAAAAGGGGTAAAATTGAG
GTTGGAGGTGACCTGATCGACCTCAAGAGAGTTGTGCTTGATGGTTCCGCGGCAACCCCTGTAACCAAAGT
TTCAGCGGAACAATGGGGTCGTCCTTAG

>AF325691.1_Linaje 8B (EE.UU-NVSL_97-7985_IA_1-4-2)

ATGTTGGAGAAATGCTTGACCGCGGGCTGTTGCTTGCGATTGCCTTCTTTGTGGTGTATCGTGCCGTTCTGT
TTTGCTGTGCTCGTCAACGCCAACAACAGCAGCAGCTCCCATTTTCAGTCGATTTATAACTTAACGCTATGTG
AGCTGAATGGCACAGAATGGCTGAGTGAGAAATTTGATTGGGCAGTGGAGACTTTTGTCATCTTTCCCGTG
TTAACTCACATTGTTTCCTATGGTGCACTCACCACCAGCCATTTCCTTGACACAGTTGGTCTGGTTACTGTGTC
CACCGCCGGGTTTCTCCACAGGCGGTATGTCTTGAGCAGCGTCTACGCGGTCTGTGCTCTGGCTGCGTTGAT
TTGCTTCATCATTAGGCTTGCGAAGAACTGCATGTCCTGGCGCTACTCTTGTACCAGATATACCAACTTTCTT
CTGGACACTAAGGGCAAACTCTATCGTTGGCGGTCGCCCGTTATCATAGAGAAAGGGGGCAGGGTTGAGG
TCGAAGGTCACCTGATCGACCTCAAAAGAGTTGTGCTTGATGGTTCCGCGGCAACCCCTTTAACCAGAGTTT
CAGCGGAACAATGGGGTCGTCTCTAG

>AF184212.1_Linaje 7(CHINA-SP)

ATGTTGGGGAAATGCTTGACCGCGGGTTGCTGCTCGCGATTGCTTTCTTTTTGGTGTATCGTGCCGTTCTGTT
TTGCTGTGCTCGTCAACGCCAGCTACAGCAGCAGCTCTCATTTACAGTTGATTTATAACTTGACGCTATGTGA
GCTGAATGGTACAGATTGGCTGGCTAATAAATTTGATTGGGCAGTGGAGAGTTTTGTCATCTTTCCTGTGTT
GACCCACATCGTTTCCTATGGTGCACTAACCACCAGCCACTTCCTTGACACAGTTGGTCTGGTTACTGTGTCT
ACCGCCGGGTTTTATCATGGGCGGTATGTCCTGAGTAGCATCTACGCGGTCTGTGCCCTGGCTGCGTTAATT
TGCTTCGTCATTAGGTTGGCGAAGAACTGTATGTCCTGGCGCTACTCATGCACCAGATACACCAACTTTCTTC
TGGACACTAAGGGCAGACTCTATCGTTGGCGGTCGCCTGTCATCATAGAGAAAGGGGGTAAGGTAGAGGT
CGAAAGCCATCTGATCGACCTCAAAAGAGTTGTGCTTGATGGGTCCGCGGCAACCCCTTTAACCAGAGTTTC
AGCGGAACAATGGGGTCGTCCCTAG

>AF176348.2_Linaje L5A(CANADA-PA8)

ATGTTGGGGAAATGCTTGACCGCGGGCTGGTGCTCGCAATTGCTTTCTTTGGGGTGTATCGTGCCGTTCTGT
TTTGCTGTGCTCGCCAACGCCAGCAACGACAGCAGCTCCCATGTACAGCTGATTTACAACTTGACGCTATGT
GAGCTGAATGGCACAGATTGGCTAGCTAACAAATTTGATTGGGCAGTGGAGAGTTTTGTCATCTTTCCCGTT
TTGACTCACATTGTCTCCTATGGTGCCCTCACTACCAGCCATTTCCTTGACACAGTCGCTTTAGTCACTGTGTC
TACCGCCGGGTTTGTTCACGGGCGGTATGTCCTAAGTAGCATCTACGCGGTCTGTGCCCTGGCTGCGTTGAC
TTGCTTCGTCATTAGGTTTGCAAAGAATTGCATGTCCTGGCGCTACGCGTGTACCAGATATACCAACTTTCTT
CTGGACACTAAGGGCAGACTCTATCGTTGGCGGTCGCCTGTCATCATAGAGAAAAGGGGCAAAGTTGAGG
TCGAAGGTCATCTGATCGACCTCAAAAGAGTTGTGCTTGATGGTTCCGTGGCAACCCCTATAACCAGAGTTT
CAGCGGAACAATGGGGTCGCCCTTAG

>AB288356.1_Linaje 4(EE.UU-EDRD-1)

ATGTTGGGGAAATGCTTGACCGCGGGCTGTTGCTCGCGATTGCCTTTTTTGTGGTGTATCGTGCCGTTCTGT
CTTGCTGCGCTCGTCAACGCCAGCGACAGCAGCAGCTCCCATTTACAGTTGATTTATAACCTGACGCTATGT
GAGCTGAATGGCACAGATTGGCTGGCTGACAAATTTGATTGGGCAGTGGAGAGTTTTGTCATCTTTCCCGT
GTTGACTCACATTGTTTCTTACTGCGCCCTCACTACCAGCCACTTCCTTGACACAGTTGGTCTGGTCGCTGTG
TCTACCGCCGGGTTTTACCACGGGCGGTATGTTCTGAGTAGCATCTATGCGGTCTGTGCCCTGGCTGCGTTG
GTTTGCTTCGTCATCAGATTGACGAAGAATTGCATGTCCTGGCGCTACTCATGTACCAGATATACCAACTTTC
TTCTGGATACCAAGGGCAGACTCTATCGTTGGCGGTCGCCCGTCATCATAGAGAAAGGGGGTAAGGTTGA
GGTTGAAGGTCATCTGATCGACCTCAAGAGAGTTGTGCTTGATGGTTCCGCGGCAACCCCTATAACCAAAG
TTTCAGCGGAACAATGGGGTCATCCCTAG

>MH791378.1_Linaje 1A(PERU-2015/15)

ATGTTGGGGAAATGCTTGACCGCGGGCTGCTGCTCGCAATTGCTTTTTTTGTGGTGTATCGTGCCGTTCTGT
TTTGTTGCGCTCGTCAACGCCAACAACAGCAGCAGCTCCCATTTACAGTTGATTTATAACCTGACGATATGT
GAGCTGAATGGCACAGATTGGCTAAATAAAAGTTTTGATTGGGCGGTGGAGACCTTCGTTATCTTTCCTGTG
TTGACTCATATTGTCTCCTATGGCGCCCTCACCACCAGCCATTTCCTTGACACAGTCGGCCTGATCACCGTGT
CTGCCGCCGGATATTACCACAGGCGGTATGTCTTGAGTAGCATTTACGCCGTCTGCGCCCTGGCTGCGTTAA
CTTGCTTCGTCATCAGGCTAACAAAAAATTGTATGTCCTGGCGTTACTCATGCACCAGGTACACTAACTTTCT
TCTGGACACCAAGGGCAAACTCTATCGTTGGCGGTCTCCTGTCATCATAGAGAAAGGGGGTAAAATTGAGG
TCGAAGGTCACCTGATCGACCTCAAGAGAGTTGTGCTTGACGGTTCCGCGGCAACCCCTGTAACCAAAGTTT
CAGCGGAACAATGGGGTCGTCCTTAG

>MH791376.1_Linaje 1A(PERU-2015/13)

ATGTTGGGGAAATGCTTGACCGCGGGCTGCTGCTCGCAATTGCTTTTTTTGTGGTGTATCGTGCCGTTCTGT
TTTGTTGCGCTCGTCAACGCCAACAACAGCAGCAGCTCCCATTTACAGTTGATTTATAACCTGACGATATGT
GAGCTGAATGGCACAGATTGGCTAAATAAAAGTTTTGATTGGGCGGTGGAGACCTTTGTTATCTTTCCTGTG
TTGACTCATATTGTCTCCTATGGCGCCCTCACCACCAGCCATTTCCTTGACACAGTCGGCCTGATCACCGTGT
CTGCCGCCGGATATTACCACAGGCGGTATGTCTTGAGTAGCATTTACGCCGTCTGCGCCCTGGCTGCGTTAA
CTTGCTTCGTCATCAGGCTAACAAAAAATTGTATGTCCTGGCGTTACTCATGCACCAGGTACACTAACTATCT
TCTGGACACCAAGGGCAAACTCTATCGTTGGCGGTCTCCTGTCATCATAGAGAAAAGGGGTAAAATTGAGG
TCGAAGGTCACCTGATCGACCTCAAGAGAGTTGTGCTTGACGGTTCCGCGGCAACCCCTGTAACCAAAGTTT
CAGCGGAACAATGGGGTCGTCCTTAG

>MH791379.1_Linaje 1A(PERU-2015/17)

ATGTTGGGGAAATGCTTGACCGCGGGCTGCTGCTCGCAATTGCCTTTTTTGTGGTGTATCGTGCCGTTCTGT
TTTGTTGCGCTCGTCAACGCCAACAACAGCAGCAGCTCTCATTTACAGTTGATTTATAACCTGACGATATGTG
AGCTGAATGGCACAGATTGGCTAAATAAAAGTTTTGATTGGGCGGTGGAGACCTTTGTTATCTTTCCTGTGT
TGACTCATATTGTCTCCTATGGCGCCCTCACCACCAGCCATTTCCTTGACACAGTCGGCCTGATCACCGTGTC
TGCCGCCGGATACTACCACGGGCGGTATGTCTTGAGTAGCATTTACGCCGTCTGCGCCCTGGCTGCGTTAAC
TTGCTTCGTCATCAGGCTAACAAAAAATTGTATGTCCTGGCGTTACTCATGCACCAGATACACTAACTTTCTT
CTGGACACCAAGGGCAAACTCTATCGTTGGCGGTCTCCTGTCATCATAGAGAAAGGGGGTAAAATTGAGGT
CGAAGGTCACCTGATCGACCTCAAGAGAGTTGTGCTTGACGGTTCCGCGGCAACCCCTGTAACCAAAGTTT
CAGCGGAACAATGGGGTCGTCCTTAG

>MH791383.1_Linaje 1A(PERU-2016/2)

ATGTTGGGGAAATGCTTGACCGCGGGCTGCTGCTCGCAATTGCCTTTTTTGTGGTGTATCGTGCCGTTCTGT
TTTGTTGCGCTCGTCAACGCCAACAACAGCAGCAGCTCCCATTTACAGTTGATTTATAACCTGACGATATGT
GAGCTGAATGGCACAGATTGGCTAAATAAAAGTTTTGATTGGGCGGTGGAGACCTTTGTTATCTTTCCTGTG
TTGACTCATATTGTCTCCTATGGCGCCCTCACTACCAGCCATTTCCTTGACACAGTCGGCCTGATCACCGTGT
CTGCCGCCGGATACTACCACGGGCGGTATGTCTTGAGTAGCATTTACGCCGTCTGCGCCCTGGCTGCGTTAA
CTTGCTTTGTCATCAGGCTAACAAAAAATTGTATGTCCTGGCGTTACTCATGCACCAGATACACTAACTTTCT
TCTGGACACCAAGGGCAAACTCTATCGTTGGCGGTCTCCTGTCATCATAGAGAAAGGGGGTAAAATTGAGG
TCGAAGGTCACCTGATCGACCTCAAGAGAGTTGTGCTTGACGGTTCCGCGGCAACCCCTGTAACCAAAGTTT
CAGCGGAACAATGGGGTCGTCCTTAG

>MH791385.1_Linaje 1A(PERU-2016/4)

ATGTTGGGGAAATGCTTGACCGCGGGCTGCTGCTCGCAATTGCCTTTTTTGTGGTGTATCGTGCCGTTCTGT
TTTGTTGCGCTCGTCAACGCCAACAACAGCAGCAGCTCCCATTTACAGTTGATTTATAACCTGACGATATGT
GAGCTGAATGGCACAGATTGGCTAAATAAAAGTTTTGATTGGGCGGTGGAGACCTTTGTTATTTTTCCTGTG
TTGACTCATATTGTCTCCTATGGCGCCCTCACCACCAGCCATTTCCTTGACACAGTCGGCCTGATCACCGTGT
CTGCCGCCGGATATTACCACGGGCGGTATGTCTTGAGTAGCATTTACGCCGTCTGCGCCCTGGCTGCGTTAA
CTTGCTTCGTCATCAGGCTAACAAAAAACTGTATGTCCTGGCGTTACTCATGCACCAGATACACTAACTTTCT
TCTGGACACCAAGGGCAAGCTCTATCGTTGGCGGTCTCCTGTCATCATAGAGAAAGGGGGTAAAATTGAGG
TCGAAGGTCACCTGATCGACCTCAAGAGAGTTGTGCTTGACGGTTCCGCGGCAACCCCTGTAACCAAAGTTT
CAGCGGAACAATGGGGTCGTCCTTAG

>MH791391.1_Linaje 1A(PERU-2017/7)

ATGTTGGGGAAATGCTTGACCGCGGGCTGCTGCTCGCAATTGCTTTTTTTGTGGTGTATCGTGCCGTTCTGT
TTTGTTGCGCTCGTCAACGCCAACAACAGCAGCAGCTCCCATTTACAGTTGATTTATAACCTGACGGTATGT
GAGCTGAATGGCACAGATTGGCTAAATAGAAGGTTTGATTGGGCGGTGGAGACCTTTGTTATCTTCCCTGT
GTTGACTCATATTGTCTCCTATGGCGCCCTCACCACCAGCCATTTCCTTGACACAGTCGGCCTGATCACCGTG
TCTGCCGCTGGATATTACCACGGGCGGTATGTCTTGAGTAGCATTTACGCCGTCTGCGCCCTGGCTGCGTTA
ACTTGCTTCGTCATCAGGCTAACAAAAAATTGTATGTCCTGGCGTTACTCATGCACCAGGTACACCAACTATC
TTCTGGACACCAAGGGCAAACTCTATCGTTGGCGGTCTCCTGTCATCATAGAGAAAGGGGGTAAAATTGAG
GTCGAAGGTCACCTGATCGACCTCAAGAGAGTTGTGCTTGACGGTTCCGCGGCAACCCCTGTAACCAAAGT
TTCAGCGGAACAATGGGGTCGTCCTTAG

>MH791386.1_Linaje 1A(PERU-2016/5)

ATGTTGGGGAAATGCTTGACCGCGGGCTGCTGCTCGCAATTGCCTTTTTTGTGGTGTATCGTGCCGTTCTGT
TTTGTTGTGCTCGTCAACGCCAACAACAGCAGCAGCTCCCATTTACAGTTGATTTATAACCTGACGATATGTG
AGCTGAATGGCACAGATTGGCTAAATAAAAGTTTTGATTGGGCGGTGGAGACCTTTGTCATCTTTCCTGTGT
TGACTCACATTGTCTCCTATGGCGCCCTCACCACCAGCCATTTCCTTGACACAGTCGGCCTGATCACCGTGTC
TGCCGCCGGATATTACCACGGGCGGTATGTCTTGAGTAGCATTTACGCCGTCTGCGCCCTGGCTGCGTTAAC
TTGCTTCGTTATCAGGCTAACAAAAAATTGTATGTCCTGGCGTTACTCATGCACCAGATACACTAACTTTCTT
CTGGACACCAAGGGCAAACTCTATCGTTGGCGGTCTCCTGTCATCATAGAGAAAGGGGGCAAAATTGAGGT
CGAAGGTCACCTGATCGACCTCAAGAGAGTTGTGCTTGACGGTTCCGCGGCAACCCCTGTAACCAAAGTTT
CAGCGGAACAATGGGGTCGTCCTTAG

>MH791388.1_Linaje 1A(PERU-2016/16)

ATGTTGGGGAAATGCTTGACCGCGGGCTGCTGCTCGCAATTGCTTTTTTTGTGGTGTATCGTGCCGTTCTGT
TTTGTTGCGCTCGTCAACGCCAACAACAGCAGCAGCTCCCATTTACAGTTGATTTATAACCTGACGATATGT
GAGCTGAATGGCACAGATTGGCTAAATAAAAGTTTTGATTGGGCGGTGGAGACCTTTGTTATCTTTCCTGTG
TTGACTCATATTGTCTCCTATGGCGCCCTCACCACCAGCCATTTCCTTGACACAGTCGGCCTGATCACCGTGT
CTGCCGCCGGATATTACCACGGGCGGTATGTCTTGAGTAGCATTTACGCCGTCTGCGCCCTAGCTGCGTTAA
CTTGCTTCGTCATCAGGCTAACGAAAAATTGTATGTCCTGGCGTTACTCATGCACCAGATACACTAACTTTCT
TCTGGACACCAAGGGCAAACTCTATCGTTGGCGGTCTCCTGTCATCATAGAGAAAGGGGGTAAAATTGAGG
TCGAAGGTCACCTGATCGACCTCAAGAGAGTTGTGCTTGACGGTTCCGCGGCAACCCCTGTAACCAAAGTTT
CAGCGGAACAATGGGGTCGTCCTTAG

>MH791390.1_Linaje 1A(PERU-2017/6)

ATGTTGGGGAAATGCTTGACCGCGGGCTGCTGCTCGCAATTGCCTTTTTTGTGGTGTATCGTGCCGTTCTGT
TTTGTTGCGCTCGTCAACGCCAACAACAGCAGCAGCTCCCATTTACAGTTGATTTATAACCTGACGATATGT
GAGCTGAATGGCACAGATTGGCTAAATAAAAGTTTTGATTGGGCGGTGGAGACCTTTGTTATCTTTCCTGTG
TTGACTCATATTGTCTCCTATGGCGCCCTCACCACCAGCCATTTCCTTGACACAGTCGGCCTGATCACCGTGT
CTGCCGCCGGATATTACCACGGGCGGTATGTCTTGAGTAGCATTTACGCCGTCTGCGCCCTGGCTGCGTTAA
TTTGCTTCGTCATCAGGCTAACAAAAAATTGTATGTCCTGGCGTTACTCATGCACCAGGTACACCAACTTTCT
TCTGGACACCAAGGGCAAACTCTATCGTTGGCGGTCTCCTGTCATCATAGAGAAAGGGGGTAAAATTGAGG
TCGAAGGTCACCTGATCGACCTCAAGAGAGTTGTGCTTGACGGTTCCGCGGCAACCCCTGTAACCAAAGTTT
CAGCGGAACAATGGGGTCGTCCTTAG

>48_Montana2019 (Linaje 1A)

GTGTTGGGGAAATGCTTGACCGCGGGCTGCTGCTCGCAATTGCCTTTTTTGTGGTGTATCGTGCCGTTCTGT
CTTGTTGTGCTCGTCAACGCCAACAACAGCAGCAGCTCCCATTTACAGTTGATTTATAACCTGACGATATGT
GAGCTGAATGGCACAGATTGGCTAAACAAAAGTTTTGATTGGGCGGTGGAGACCTTTGTCATCTTTCCTGT
GTTGACTCATATTGTCTCCTATGGCGCCCTTACCACCAGTCATTTCCTTGACACAGTCGGCCTGATCACCGTG
TCTGCCGCCGGATATTACCACGGGCGGTATGTCTTGAGTAGCATTTACGCCGTCTGCGCCTTAGCCGCGTTA
ATTTGCTTCATCATCAAGCTAACAAAAAATTGTATGTCCTGGCGTTACTCATGCACCAGGTACACTAATTTTC
TTCTGGACACCAAGGGCAAACTCTATCGTTGGCGGTCTCCTGTCATCATAGAGAAAGGGGGTAAAGTTGAG
GTCCAAGGTCACCTGATAGACCTCAAAAGAGTTGTGCTTGACGGTTCCGCGGCTACCCCTGTAACCAAAGTT
TCAGCGGAACAATGGGGTCGTCCTTAG

>47_Montana2019 (Linaje 1A)

GTGTTGGGGAAATGCTTGACCGCGGGCTGCTGCTCGCAATTGCCTTTTTTGTGGTGTATCGTGCCGTTCTGT
TTTGTTGTGCTCGTCAACGCCAACAACAGCAGCAGCTCCCATTTGCAGTTGATTTATAACCTGACGATATGT
GAGCTGAATGGCACAGATTGGCTAAACGAACATTTTGATTGGGCGGTGGAGACCTTTGTCATCTTTCCTGTG
TTGACTCACATTGTCTCCTATGGTGCCCTCACCACCAGCCATTTCCTTGACACAGTCGGCCTGATCACCGTGT
CTACCGCCGGATATTACCACGGGCGGTATGTCTTGAGTAGCATTTACGCCGTCTGCGCCCTAGCTGCGTTAA
CTTGCTTCATCATCAGGCTAACGAAAAACTGTATGTCCTGGCGTTACTCATGCACCAGGTACACTAATTTTCT
TCTGGACACCAAGGGCAAACTCTATCGTTGGCGGTCTCCTGTCATCATAGAGAAAGGGGGTAAAGTTGAGG
TCGAAGGTCACCTTATCGACCTCAAGAGAGTTGTACTTGACGGTTCCGCGGCTACCCCTGTAACCAAAGTTT
CAGCGGAACAATGGGGTCGTCCTTAG

>46_Montana2019 (Linaje 1A)

GTGTTGGGGAAATGCTTGACCGCGGGCTGCTGCTCGCAATTGCCTTTTTTGTGGTGTATCGTGCCGTTCTGT
TTTGTTGTGCTCGTCAACGCCAACAACAGCAGCAGCTCCCATTTACAGTTGATTTATAACCTGACGATATGTG
AGCTGAATGGCACAGATTGGCTAAACGAACATTTTGATTGGGCGGTGGAGACCTTTGTCATCTTTCCTGTGT
TGACTCACATTGTCTCCTATGGTGCCCTCACCACCAGCCATTTCCTTGACACAGTCGGCCTGATCACCGTGTC
TACCGCCGGATATTACCACGGGCGGTATGTCTTGAGTAGCATTTACGCCGTCTGCGCCCTAGCTGCGTTAAC
TTGCTTCATCATCAGGCTAACGAAAAACTGTATGTCCTGGCGTTACTCATGCACCAGGTACACTAATTTTCTT
CTGGACACCAAGGGCAAACTCTATCGTTGGCGGTCTCCTGTCATCATAGAGAAAGGGGGTAAAGTTGAGGT
CGAAGGTCACCTGATCGACCTCAAGAGAGTTGTACTTGACGGTTCCGCGGCTACCCCTGTAACCAAAGTTTC
AGCGGAACAATGGGGTCGTCCTTAG

>45_Montana2019 (Linaje 1A)

GTGTTGGGGAAATGCTTGACCGCGGGCTGCTGCTCGCAATTGCCTTTTTTGTGGTGTATCGTGCCGTTCTGT
TTTGTTGTGCTCGTCAACGCCAACAACAGCAGCAGCTCCCATTTACAGTATGTTTATAACCTGACGATATGTG
AGCTGAATGGCACAGATTGGCTAAACGTTCATTTTGATTGGGCGGTGGAGACCTTTGTCATCTTTCCTGTGT
TGACTCACATTGTCTCCTTTGGTGCCCTTACCACCAGCCATTTCCTTGACACATTCGGCCTGATCACCGTGTCT
ACCGCCGGATATTACCACGGGCGGTATGTCTTGAGTAGCATTTACGCCGTCTGCGCCCTAGCCGCGTTAATT
TGCTTCATCATCAAGCTAACAAAAAATTGTATGTCCTGGCGTTACTCATGCACCAGGTACACTAATTTTCATC
TGGACACCAAGGGCAAACTCTATCGTTGGCGGTCTCCTGTCATCATAGAGAAAGGGGGTAAAGTTGAGGTC
CAAGGTCACCTGATAGACCTCAAAAGAGTTGTGCTTGACGGTTCCGCGGCTACCCCTGTAACCAAAGTTTCA
GCGGAACAATGGGGTCGTCCTTAG

>44_Montana2019 (Linaje 1A)

ATGTTGGGGAAATGCTTGACCGCGGGCTGCTGCTCGCAATTGCCTTTTTTGTGGTGTATCGTGCCGTTCTGT
TTTGTTGCGCTCGTCAACGCCGGCAACAGCATCAGCTCCCATTTACAGGTGATTTATAACCTGACGATATGT
GAGCTGAATGGCACAGATTGGCTAAATAAAAGCTTTGATTGGGCGGTGGAGACCTTTGTTATCTTTCCTGTG
GTGACTCATATTGTCTCCTATGGCGCCTTCACCACCAGCCATTTCCTTGACACAGTCGGCCTGATCACCGTGT
CTGCCGCCGGATATTACCCCGGACGGTATGTTTTGAGTAGCATTTACGCCGTTTGCGCCCTGGATGCGTTAA
CTTGCTTCGTCATCAGGCTAACAAAAAATTGTATGTCCTGGCGTTATTCATGCACCAGGTACAGTAATTTTCT
TCTGGACACCAAGGGCAAACTCTATCGTTGGCGGTCTCCTGTCATCATAGAGAAAGGGGGTAAAATAGAG
GTAGAAGGTCACATGATCGACCTCAAGAGAGTTGTACTTGACGGTTCCGCGGCTACCCCTGTAACCAAAGT
TTCAGCGGAACAATGGGGTCGTCCTTAG

>43_Montana2019 (Linaje 1A)

ATGTTGGGGAAATGCTTGACCGCGGGCTGCTGCTCGCAATTGCCTTTTTTGTGGTGTATCGTGCCGTTCTGT
TTTGTTGCGCTCGTCAACGCCGGCAACAGCATCAGCTCCCATTTACAGGTGATTTATAACCTGACGATATGT
GAGCTGAATGGCACAGATTGGCTAAATAAAAGCTTTGATTGGGCGGTGGAGACCTTTGTTATCTTTCCTGTG
TTGACTCATATTGTCTCCTATGGCGCCCTCACCACCAGCCATTTCCTTGACACAGTCGGCCTGATCACCGTGT
CTGCCGCCGGATATTACCACGGACGGTATGTTTTGAGTAGCATTTACGCCGTCTGCGCCCTGGCTGCATTAA
CTTGCTTCGTCATCAGGCTAACAAAAAATTGTATGTCCTGGCGTTATTCATGCACCAGGTACACTAATTTTCT
TCTGGACACCAAGGGCAAACTCTATCCTTGGCGGTCTCCTGTCATCATAAAGAAAGGGGGGAAAATAGAGG
TAGAAGGTCACATGATCGACCTCAAGAGAGTTGTACTTGACGGTTCCGCGGCTACCCCTGTAACCAAAGTTT
CAGCGGAACAATGGGGTCGTCCTTAG

>42_Montana2019 (Linaje 1A)

ATGTTGGGGAAATGCTTGACCGCGGGCTGCTGCTCGCAATTGCCTTTTTTGTGGTGTATCGTGCCATTCTGT
TTTGTTGCGCTCGTCAACACCCGCAACACCAGCATCTCCCTTTTACTTTTGATTTATAACCTGACGATATGTGA
GCTGAATGGCACAGATTGGCTAAATAAAAGCTTTGATTGGGCGGTGGAGACCTTTGTTATCTTTCCTGTGTT
GACTCATATTGTCTCCTATGGCGCCCTCACCACCAGCCATTTCCTTGACACAGTCGGCCTGATCACCGTGTCT
GCCGCCGGATATTACCACGGACGGTATGTTTTAAGTAGCATTTACGCCGTCTGCGCCCTGGCTGCGTTAACT
TGCTTCGTCATCTCGCTATCTAAAAATTGTATGTCCTGGCGTTATTCTTGCACCAAGTACACTACTTTTCTTCT
GGACACCAAGTGCAATCTCTCTCGCGGAAGTTCTCCTGTGATGATACACCGCTGGGGTTAAAACGAAGTCT
CATGTTTCCTGTCCCTCTTCAAAAGAGTTGTGCTTGACGGTTCCGCGGCTACCCCTGTAACCAAAGTTTCAGC
GGAACAATGGGGTCGTCCTTAG

>41_Montana2019 (Linaje 1A)

ATGTTGGGGAAATGCTTGACCGCGGGCTGCTGCTCGCAATTGCCTTTTTTGTGGTGTATCGTGCCGTTCTGT
TTTGTTGCGCTCGTCAACGCCAGCAACAACAGCAGCTCCCATTTACTGTTGATTTATAACCTGACGATATGTG
AGCTGAACGGCACAGATTGGCTAAATAAAAGTTTTGATTGGGCGGTGGAGACCTTTGTTATCTTTCCTGTGT
TGACTCATATTGTCTCCTATGGCGCCCTCACCACCAGCCATTTCCTTGACACAGTCGGCCTGATCACCGTGTC
TGCCGCCGGATATTACCACGGACGGTATGTTTTGAGTAGCATTTACGCCGTCTGCGCCCTGGCTGCGTTAAC
TTGCTTCATCATCAGGCTAACAAAAAATTGTATGTCCTGGCGTTACTCATGCACCAGGTACACTAATTTTCTT
CTGGACACCAAGGGCAAACTCTATCGTTGGCGGTCTCCTGTCATCATAGAGAAAGGGGGTAAAGTTGACGT
CGAAGGTCACCTGATCGACCTCAAGAGAGTTGTACTTGACGGTTCCGCGGCTACCCCTGTAACCAAAGTTTC
AGCGGAACAATGGGGTCGTCCTTAG

>40_Montana2019 (Linaje 1A)

ATGTTGGGGAAATGCTTGACCGCGGGCTGCTGCTCGCAATTGCCTTTTTTGTGGTGTATCGTGCCGTTCTGT
TTTGTTGCGCTCGTCAACGCCAGCAACAACAGCAGCTCCCATTTACAGTTGATTTATAACCTGACGATATGT
GAGCTGAACGGCACAGATTGGCTAAATAAAAGTTTTGATTGGGCGGTGGAGACCTTTGTTATCTTTCCTGTG
TTGACTCATATTGTCTCCTATGGCGCCCTCACCACCAGCCATTTCCTTGACACAGTCGGCCTGATCACCGTGT
CTGCCGCCGGATATTACCACGGACGGTATGTTTTGAGTAGCATTTACGCCGTCTGCGCCCTGGCTGCGTTAA
CTTGCTTCATCATCAGGCTAACAAAAAATTGTATGTCCTGGCGTTACTCATGCACCAGGTACACTAATTTTCT
TCTGGACACCAAGGGCAAACTCTATCGTTGGCGGTCTCCTGTCATCATAGAGAAAGGGGGTAAAGTTGAGG
TCGAAGGTCACCTGATCGACCTCAAGAGAGTTGTACTTGACGGTTCCGCGGCTACCCCTGTAACCAAAGTTT
CAGCGGAACAATGGGGTCGTCCTTAG

>39_Montana2019 (Linaje 1A)

ATGTTGGGGAAATGCTTGACCGCGGGCTGCTGCGCGCGGGGGCTTTTTTGGTGGTGGATCGATAAGTTCTG
TTTGGTTTCGCTCGTCAATGCCGAGAACAGCAGCAGCTCCCATTTAAGATAAATCCATATTCGGGCGGTATG
CGGTCTGAATGGCACAAATTGGCTAAATAGAAGGTTTGATTGGGCGGTGTAGACCTTTGTTGTCTTTCCTGT
GTTGACTCATATTGTCTCCTATGGCGCCCTCACCACCAGCCATTTCCTTGACACAGTCGGTTTGATTAAGGTG
TCTGCCGCCGGATATTACCACGGGCGGCATGTCAAAAGTAGCATTTACGCCGTGTGCGCCCTGGCTGCGAT
GGGCCGATTTGTCATCAGACTAACAAAAAATTGCACCTCCGGGCGGGACTCACGCACCAGATACGCTAACT
ATCTTCTGGACACCAAGGGCAAACTATATCGTTGGCGGTCTCCTGTCATCATAGAGAAAGGGGGTAAAATT
GAGGTCGAAGGTCACCTGATCGACCTCAAGAGAGTTGTGCTTGACGGTTCCGCGGCGACTCCTGTACCCTA
AGTTTCAGCGGAACAATGGGGTCGTCCTTAG

>38_Montana2019 (Linaje 1A)

ATGTTGGGGAAATGCTTGACCGCGGGCTGCTGCGCGCGGGGGCTTTTTTGGTGGTGGATCGATAAGTTCTG
TTTGGTTTCGCTCGTCAATGCCGAGAACAGCAGCAGCTCCCATTTAAGATAAATCCATATTCGGGCGGTATG
CGGTCTGAATGGCACAAATTGGCTAAATAGAAGGTTTGATTGGGCGGTGTAGACCTTTGTTGTCTTTCCTGT
GTTGACTCATATTGTCTCCTATGGCGCCCTCACCACCAGCCATTTCCTTGACACAGTCGGTTTGATGAAGGTG
TCTGCCGCCGGATATTACCACGGGCGGCATGTCAAAAGTAGCATTTACGCCGTGTGCGCCCTGGCTGCGAT
GGGCCGATTTGTCATCAGACTAACAAAAAATTGTACCTCCGGGCGGGACTCACGCACCAGATACGCTAACT
ATCTTCTGGACACTAAGGGCAAACTATATCGTTGGCGGTCTCCTGTCATCATAGAGAAAGGGGGTAAAATT
GAGGTCGAAGGTCACCTGATCGACCTCAAGAGAGTTGTGCTTGACGGTTCCGCGGCGACTCCTGTAACCAA
AGTTTCAGCGGAACAATGGGGTCGTCCTTAG

>37_Montana2019 (Linaje 1A)

ATGTTGGGGAAATGCTTGACCGCGGGCTGCTGCGCGGGGGCTTTTTGGTGGTGGATCGATAAGTTCTG
TTTGGTTTCGCTCGTCAATGCCGAGAACAGCAGCAGCTCCCATTTAAGATAAATCCATATTCGGGCGGTATG
CGGTCTGAATGGCACAAATTGGCTAAATAGAAGGTTTGATTGGGCGGTGTAGACCTTTGTTGTCTTTCCTGT
GTTGACTCATATTGTCTCCTATGGCGCCCTCACCACCAGCCATTTCCTTGACACAGTCGGTTTGATTAAGGTG
TCTGCCGCCGGATATTACCACGGACGGCATGTCAAAAGTAGCATTTACGCCGTGTGCGCCCTGGCTGCGAT
GGGCCGATTTGTCATCAGACAACGAAAAAATTGCACCTCCGGGCGGGACTCACGCACCAGATACGCTAACT
ATCTTCTGGACACCAAGGGCAAACTATATCGTTGGCGGTCTCCTGTCATCATAGAGAAAGGGGGCAAAATT
GAGGTTGTCTTTCACCTGATCGACCTCAAGAGAGTTGTGCTTGACGGTTCCGCGGCGACTCCTGTACCCTAA
GTTTCAGCGGAACAATGGGGTCGTCCTTAG

>36_Montana2019 (Linaje 1A)

ATGTTGGGGAAATGCTTGACCGCGGGCTGCTGCGCGGGGGCTTTTTGGTGGTGGATCGATAAGTTCTG
TTTGGTTTCGCTCGTCAATGCCGAGAACAGCAGCAGCTCCCATTTAAGATAAATCCATATTCGGGCGGTATG
CGGTCTGAATGGCACAAATTGGCTAAATAGAAGGTTTGATTGGGCGGTGTAGACCTTTGTTGTCTTTCCTGT
GTTGACTCATATTGTCTCCTATGGCGCCCTCACCACCAGCCATTTCCTTGACACAGTCGGTTTGATGAAGGTG
TCTGCCGCCGGATATTACCACGGGCAGCGTGTCAAAAGTAGCATTTACGCCGTGTGCGCCCTGGCTGCGAT
GGGCCGATTTGTCATCAGACAACGAAAAAATTGTACCTCCGGGCGGGACTCACGCACCAGATACGCTAACT
ATCTTCTGGACACTAAGGGCAAACTATATCGTTGGCGGTCTCCTGTCATCATAGAGAAAGGGGGTAAAATT
GAGGTTGGAGGTCACCTGATCGACCTCAAGAGAGTTGTGCTTGACGGTTCCGCGGCGACTCCTGTAACCAA
AGTTTCAGCGGAACAATGGGGTCGTCCTTAG

>35_Montana2019 (Linaje 1A)

ATGTTGGGGAAATGCTTGACCGCGGGCTGCTGCTCGCAATTGCTTTTTTTGTGGTGTATCGTGCCGTTCTGT
TTTGTTGCGCTCGTCAACGCCAGCAACAGCAGCAGCTCCCATTTTCAGTTGATTTATAACCTGACGATATGC
GAGCTGAATGGCACAGATTGGCTAAATAGAAGTTTTGATTGGGCGGTAGAGACCTTTGTTATCTTTCCTGTG
TTGACTCATATTGTCTCCTATGGCGCCCTCACCACCAGCCATTTCCTTGACACAGTCGGCCTGATCACCGTGT
CTGCCGCCGGATATTACCACGGGCGGTATGTCTTGAGTAGCATTTATGCCGTCTGCGCCCTGGCTGCGTTAA
CTTGCTTTGTCATCAGGCTAACAAAAAATTGCATGTCCTGGCGTTACTCGTGCACCAGGTACACTAACTATCT
TCTGGACACTAAGGGCAAACTCTATCGTTGGCGGTCTCCTGTCATCATAGAGAAAGGGGGTAAAATTGAGG
TCGAAGGTCACCTGATCGACCTCAAGAGAGTTGTGCTTGACGGTTCCGCGGCAACCCCTGTAACCAAAGTTT
CAGCGGAACAATGGGGTCGTCCTTAG

>34_Montana2019 (Linaje 1A)

ATGTTGGGGAAATGCTTGACCGCGGGCTGCTGCTCGCAATTGCTTTTTTTGTGGTGTATCGTGCCGTTCTGT
TTTGTTGCGCTCGTCAACGCCAGCAACAGCAGCAGCTCCCATTTTCAGTTGATTTATAACCTGACGATATGC
GAGCTGAATGGCACAGATTGGCTAAATAGAAGTTTTGATTGGGCGGTAGAGACCTTTGTTATCTTTCCTGTG
TTGACTCATATTGTCTCCTATGGCGCCCTCACCACCAGCCATTTCCTTGACACAGTCGGCCTGATCACCGTGT
CTGCCGCCGGATATTACCACGGGCGGTATGTCTTGAGTAGCATTTATGCCGTCTGCGCCCTGGCTGCGTTAA
CTTGCTTTGTCATCAGGCTAACAAAAAATTGCATGTCCTGGCGTTACTCGTGCACCAGGTACACTAACTATCT
TCTGGACACTAAGGGCAAACTCTATCGTTGGCGGTCTCCTGTCATCATAGAGAAAGGGGGTAAAATTGAGG
TCGAAGGTCACCTGATCGACCTCAAGAGAGTTGTGCTTGACGGTTCCGCGGCAACCCCTGTAACCAAAGTTT
CAGCGGAACAATGGGGTCGTCCTTAG

>33_Montana2019 (Linaje 1A)

ATGTTGGGGAAATGCTTGACCGCGGGCTGCTGCTCGCAATTGCCTTTTTTGTGGTGTATCGTGCCGTTCTGT
TTTGTTGTGCTCGTCAACGCCAACAACAGCAACAGCTCCCATTTACAGTTGATTTATAACCTGACGATATGTG
AGCTGAATGGCACAGATTGGCTAAATAAAAGTTTTGATTGGGCGGTGGAGACCTTTGTTATCTTTCCTGTGT
TGACTCATATTGTCTCCTACGGCGCCCTCACCACCAGCCATTTCCTTGACACAGTCGGCCTGATCACCGTGTC
TGCCGCCGGATACTACCACGGACGGTATGTCTTAAGTAGCATTTACGCCGTCTGCGCCATGGCTGCGTTAAC
TTGCTTCGTCATCAGGCTAACAAAAAATTGTATGTCCTGGCGTTACTCATGTACCAGGTACACTAATTTTCTT
CTGGACACCAAGGGCAAACTCTATCGTTGGCGGTCTCCTGTCATCATAGAGAAAGGGGGTAAAGTTGAGGT
CGAAGGTCACCTGATCGACCTCAAGAGAGTTGTGCTTGACGGTTCCGCGGCAACCCCTGTAACCAAAGTTT
CAGCGGAACAATGGGGTCGTCCTTAG

>32_Montana2019 (Linaje 1A)

ATGTTGGGGAAATGCTTGACCGCGGGCTGCTGCTCGCAATTGCCTTTTTTGTGGTGTATCGTGCCGTTCTGT
TTTGTTGTGCTCGTCAACGCCAACAACAGCAACAGCTCCCATTTACAGTTGATTTATAACCTGACGATATGTG
AGCTGAATGGCACAGATTGGCTAAATAAAAGTTTTGATTGGGCGGTGGAGACCTTTGTTATCTTTCCTGTGT
TGACTCATATTGTCTCCTACGGCGCCCTCACCACCAGCCATTTCCTTGACACAGTCGGCCTGATCACCGTGTC
TGCCGCCGGATACTACCACGGACGGTATGTCTTAAGTAGCATTTACGCCGTCTGCGCCATGGCTGCGTTAAC
TTGCTTCGTCATCAGGCTAACAAAAAATTGTATGTCCTGGCGTTACTCATGCACCAGATACACTAATTTTCTT
CTGGACACCAAGGGCAAACTCTATCGTTGGCGGTCTCCTGTCATCATAGAGAAAGGGGGTAAAGTTGAGGT
CGAAGGTCACCTGATCGACCTCAAGAGAGTTGTGCTTGACGGTTCCGCGGCAACCCCTGTAACCAAAGTTT
CAGCGGAACAATGGGGTCGTCCTTAG

>31_Montana2019 (linaje 1A)

ATGTTGGGGAAATGCTTGACCGCGGGCTGCTGCTCGCAATTGCCTTTTTTGTGGTGTATCGTGCCGTTCTGT
TTTGTTGTGCTCGTCAACGCCAACAACAGCAACAGCTCCCATTTACAGTTGATTTATAACCTGACGATATGTG
AGCTGAATGGCACAGATTGGCTAAATAAAAGTTTTGATTGGGCGGTGGAGACCTTTGTTATCTTTCCTGTGT
TGACTCATATTGTCTCCTACGGCGCCCTCACCACCAGCCATTTCCTTGACACAGTCGGCCTGATCACCGTGTC
TGCCGCCGGATACTACCACGGACGGTATGTCTTAAGTAGCATTTACGCCGTCTGCGCCATGGCTGCGTTAAC
TTGCTTCGTCATCAGGCTAACAAAAAATTGTATGTCCTGGCGTTACTCATGTACCAGGTACACTAATTTTCTT
CTGGACACCAAGGGCAAACTCTATCGTTGGCGGTCTCCTGTCATCATAGAGAAAGGGGGTAAAGTTGAGGT
CGAAGGTCACCTGATCGACCTCAAGAGAGTTGTGCTTGACGGTTCCGCGGCAACCCCTGTAACCAAAGTTT
CAGCGGAACAATGGGGTCGTCCTTAG

>30_Montana2019 (Linaje 1A)

ATGTTGGGGAAATGCTTGACCGCGGGCTGCTGCTCGCAATTGCCTTTTTTGTGGTGTATCGTGCCGTTCTGT
TTTGTTGTGCTCGTCAACGCCAACAACAGCAACAGCTCCCATTTACAGTTGATTTATAACCTGACGATATGTG
AGCTGAATGGCACAGATTGGCTAAATGAAAGTTTTGATTGGGCGGTGGAGACCTTTGTTATCTTTCCTGTGT
TGACTCATATTGTCTCCTACGGCGCCCTCACCACCAGCCATTTCCTTGACACAGTCGGCCTGATCACCGTGTC
TGCCGCCGGATACTACCACGGACGGTATGTCTTAAGTAGCATTTACGCCGTCTGCGCCATGGCTGCGTTAAC
TTGCTTCGTCATCAGGCTAACAAAAAATTGTATGTCCTGGCGTTACTCATGTACCAGGTACACTAATTTTCTT
CTGGACACCAAGGGCAAACTCTATCGTTGGCGGTCTCCTGTCATCATAGAGAAAGGGGGTAAAGTTGAGGT
CGAAGGTCACCTGATCGACCTCAACAGAGTTGTGCTTGACGGTTCCGCGGCAACCCCTGTAACCAAAGTTTC
AGCGGAACAATGGGGTCGTCCTTAG

>29_Montana2019 (Linaje 1A)

ATGTTGGGGAAATGCTTGACCGCGGGCTGCTGCTCGCAATTGCTTTTTTTGTGGTGTATCGTGCCGTTCTGT
TTTGTTGCGCTCGTCAACGCCAGCAACAGCAGCAGCTCCCATTTTCAGTTGATTTATAACCTGACGATATGC
GAGCTGAATGGCACAGATTGGCTAAATAGAAATTTTGATTGGGCGGTAGAGACCTTTGTTATCTTTCCTGTG
TTGACTCATATTGTCTCCTATGGCGCCCTCACCACCAGCCATTTCCTTGACACAGTCGGCCTGATCACCGTGT
CTGCCGCCGGATATTACCACGGGCGGTATGTCTTGAGTAGCATTTATGCCGTCTGCGCCCTGGCTGCATTAA
TTTGCTTTGTCATCAGGCTAACAAAAAATTGCATGTCCTGGCGTTACTCGTGCACCAGGTACACTAACTATCT
TCTGGACACTAAGGGCAAACTCTATCGTTGGCGGTCTCCTGTCATCATAGAGAAAGGGGGTAAAATTGAGG
TCGAAGGTCACCTGATCGACCTCAAGAGAGTTGTGCTTGACGGTTCCGCGGCAACCCCTGTAACCAAAGTTT
CAGCGGAACAATGGGGTCGTCCTTAG

>28_Montana2019 (Linaje 1A)

ATGTTGGGGAAATGCTTGACCGCGGGCTGCTGCTCGCAATTGCTTTTTTTGTGGTGTATCGTGCCGTTCTGT
TTTGTTGCGCTCGTCAACGCCAGCAACAGCAGCAGCTCCCATTTTCAGTTGATTTATAACCTGACGATATGC
GAGCTGAATGGCACAGATTGGCTAAATAGAAGTTTTGATTGGGCGGTAGAGACCTTTGTTATCTTTCCTGTG
TTGACTCATATTGTCTCCTATGGCGCCCTCACCACCAGCCATTTCCTTGACACAGTCGGCCTGATCACCGTGT
CTGCCGCCGGATATTACCACGGGCGGTATGTCTTGAGTAGCATTTATGCCGTCTGCGCCCTGGCTGCGTTAA
CTTGCTTTGTCATCAGGCTAACAAAAAATTGCATGTCCTGGCGTTACTCGTGCACCAGGTACACTAACTATCT
TCTGGACACTAAGGGCAAACTCTATCGTTGGCGGTCTCCTGTCATCATAGAGAAAGGGGGTAAAATTGAGG
TCGAAGGTCACCTGATCGACCTCAAGAGAGTTGTGCTTGACGGTTCCGCGGCAACCCCTGTAACCAAAGTTT
CAGCGGAACAATGGGGTCGTCCTTAG

>27_Montana2019 (Linaje 1A)

ATGTTGGGGAAATGCTTGACCGCGGGCTGCTGCTCGCAATTGCTTTTTTTGTGGTGTATCGTGCCGTTCTGT
TTTGTTGCGCTCGTCAACGCCAGCAACAGCAGCAGCTCCCATTTTCAGTTGATTTATAACCTGACGATATGC
GAGCTGAATGGCACAGATTGGCTAAATAGAAGTTTTGATTGGGCGGTAGAGACCTTTGTTATCTTTCCTGTG
TTGACTCATATTGTCTCCTATGGCGCCCTCACCACCAGCCATTTCCTTGACACAGTCGGCCTGATCACCGTGT
CTGCCGCCGGATATTACCACGGGCGGTATGTCTTGAGTAGCATTTATGCCGTCTGCGCCCTGGCTGCGTTAA
CTTGCTTTGTCATCAGGCTAACAAAAAATTGCATGTCCTGGCGTTACTCGTGCACCAGGTACACTAACTATCT
TCTGGACACTAAGGGCAAACTCTATCGTTGGCGGTCTCCTGTCATCATAGAGAAAGGGGGTAAAATTGAGG
TCGAAGGTCACCTGATCGACCTCAAGAGAGTTGTGCTTGACGGTTCCGCGGCAACCCCTGTAACCAAAGTTT
CAGCGGAACAATGGGGTCGTCCTTAG

>26_Montana2019 (Linaje 1A)

ATGTTGGGGAAATGCTTGACCGCGGGCTGCTGCTCGCAATTGCTTTTTTTGTGGTGTATCGTGCCGTTCTGT
TTTGTTGCGCTCGTCAACGCCAGCAACAGCAGCAGCTCCCATTTTCAGTTGATTTATAACCTGACGATATGC
GAGCTGAATGGCACAGATTGGCTAAATAGAAGTTTTGATTGGGCGGTAGAGACCTTTGTTATCTTTCCTGTG
TTGACTCATATTGTCTCCTATGGCGCCCTCACCACCAGCCATTTCCTTGACACAGTCGGCCTGATCACCGTGT
CTGCCGCCGGATATTACCACGGGCGGTATGTCTTGAGTAGCATTTATGCCGTCTGCGCCCTGGCTGCGCTAA
CTTGCTTTGTCATCAGGCTAACAAAAAATTGCATGTCCTGGCGTTACTCGTGCACCAGGTACACTAACTATCT
TCTGGACACTAAGGGCAAACTCTATCGTTGGCGGTCTCCTGTCATCATAGAGAAAGGGGGTAAAATTGAGG
TCGAAGGTCACCTGATCGACCTCAAGAGAGTTGTGCTTGACGGTTCCGCGGCAACCCCTGTAACCAAAGTTT
CAGCGGAACAATGGGGTCGTCCTTAG

>25_montana2019 (Linaje 1A)

ATGTTGGGGAAATGCTTGACCGCGGGCTGCTGCTCGCAATTGCTTTTTTTGTGGTGTATCGTGCCGTTCTGT
TTTGTTGCGCTCGTCAACGCCAGCAACAGCAGCAGCTCCCATTTTCAGTTGATTTATAACCTGACGATATGC
GAGCTGAATGGCACAGATTGGCTAAATAGAAGTTTTGATTGGGCGGTAGAGACCTTTGTTATCTTTCCTGTG
TTGACTCATATTGTCTCCTATGGCGCCCTCACCACCAGCCATTTCCTTGACACAGTCGGCCTGATCACCGTGT
CTGCCGCCGGATATTACCACGGGCGGTATGTCTTGAGTAGCATTTATGCCGTCTGCGCCCTGGCTGCGTTAA
CTTGCTTTGTCATCAGGCTAACAAAAAATTGCATGTCCTGGCGTTACTCGTGCACCAGGTACACTAACTATCT
TCTGGACACTAAGGGCAAACTCTATCGTTGGCGGTCTCCTGTCATCATAGAGAAAGGGGGTAAAATTGAGG
TCGAAGGTCACCTGATCGACCTCAAGAGAGTTGTGCTTGACGGTTCCGCGGCAACCCCTGTAACCAAAGTTT
CAGCGGAACAATGGGGTCGTCCTTAG
